# PERK inhibition rewires translational and CMGC protein kinase networks into an antiviral state

**DOI:** 10.1101/2025.10.08.681090

**Authors:** Mohammed Samer Shaban, Hendrik Schuele Weiser, Axel Weber, Johanna Meier-Soelch, Franziska Dort, Christin Mayr-Buro, Michael Poppe, Nadja Karl, Jochen Wilhelm, John Ziebuhr, Uwe Linne, Michael Kracht

**Affiliations:** Rudolf Buchheim Institute of Pharmacology, Justus Liebig University, Giessen, Germany; Institute of Medical Virology, Justus Liebig University, Giessen, Germany; Institute for Lung Health, Justus Liebig University Giessen, Giessen, Germany; Mass Spectrometry Facility of the Department of Chemistry, Philipps University, Marburg, Germany; German Center for Lung Research (DZL) and Universities of Giessen and Marburg Lung Center(UGMLC); Member of the Excellence Cluster Cardio-Pulmonary Institute (CPI), Giessen, Germany

**Keywords:** Coronavirus, kinome, PERK inhibitor, translatome, phospho-proteome

## Abstract

Protein kinases (PKs) are central regulators of cellular signaling, yet only a small fraction of the human kinome is targeted therapeutically, and kinase–substrate relationships remain incompletely defined. Here, we systematically characterize kinome regulation during human coronavirus 229E (HCoV-229E) infection across transcriptomic, translational, proteomic, and phospho-proteomic layers. We reveal that pharmacological inhibition of the ER stress sensor kinase PERK reprograms host protein biosynthesis and phospho-proteomic landscapes, simultaneously blocking viral nucleocapsid phosphorylation and modulating multiple host kinases. This rewiring antagonizes virus-induced translational shutdown, along with pronounced regulation of the CMGC kinase family, a pattern conserved in SARS-CoV and SARS-CoV-2 infected cells. Comparative analyses with PERK depletion distinguish on-target from off-target effects of PERK inhibition. Our findings uncover the kinome-scale consequences of PERK perturbation in coronavirus infection and demonstrate how the polypharmacology of PERK inhibitors can be harnessed to establish a potent antiviral state, revealing new avenues for host-directed antiviral strategies.

## Introduction

Protein kinases (PKs) are pivotal regulators of cellular states and constitute one of the most therapeutically relevant classes of drug targets in oncology and inflammatory diseases (Cohen *et al*, 2022). To date, the U.S. Food and Drug Administration (FDA) has approved 85 PK inhibitors (PKi) for these indications; however, no PKi has yet been developed for the treatment of viral or other infectious diseases (Roskoski, 2025).

A landmark kinome census identified 518 human kinases characterized by conserved catalytic domains that bind ATP and mediate the transfer of the γ-phosphate to specific substrates (Manning *et al*, 2002). This enzymatic superfamily, termed the kinome, constitutes approximately 2–3% of the ~20,000 human protein-coding genes (Roskoski, 2025). The kinome is subdivided into eukaryotic protein kinases (ePKs) and atypical kinases (aPKs, e.g. lipid kinases) (Kanev *et al*, 2019). Phylogenetic analyses classify the 479 ePKs into seven principal families: AGC, CAMK, CMGC, CKI, STE, TK, and TKL, along with a heterogeneous group termed “Other” kinases. These relationships are illustrated in the widely referenced kinome dendrogram (Manning *et al*., 2002; Wilson *et al*, 2018). Subsequent refinements of kinase classification, reflected in resources such as KinHub (http://www.kinhub.org/kinmap/) and Kinome.org (www.kinome.org<u>), now</u> catalogue between 536 and 710 confirmed or putative human kinases (Eid *et al*, 2017; Moret *et al*, 2021).

The 85 clinically approved small-molecule PKi target only 24 eukaryotic protein kinases in total, meaning that only ∼5% of the human kinome is currently accessible for therapeutic purposes (Roskoski, 2025). With limited exceptions, these agents act as ATP-competitive inhibitors, exploiting the conserved architecture of the kinase ATP-binding pocket. This structural conservation confers an inherent propensity for PKi to engage multiple kinases, a phenomenon known as kinase polypharmacology. While this lack of absolute specificity is often viewed as a challenge, it is now recognized as a class effect that can be therapeutically advantageous; indeed, more than a dozen approved PKi function as multi-kinase inhibitors (Fabian *et al*, 2005; Karaman *et al*, 2008; Roskoski, 2025).

Phosphorylation of serine (Ser), threonine (Thr), and tyrosine (Tyr) residues by ePKs is a pervasive post-translational modification, with ultra-deep phospho-proteomic profiling predicting its occurrence on up to 90% of proteins expressed within a single cell type (Sharma *et al*, 2014). To date, more than 200,000 phosphorylation sites have been experimentally identified in human proteins, although the true extent of the phospho-proteome remains unknown (Hornbeck *et al*, 2019; Ochoa *et al*, 2020). Despite remarkable advances in global phospho-proteomic technologies, the systematic identification and functional validation of regulatory phosphorylation sites remains an immense challenge, with a substantial fraction of the kinome’s functional landscape yet to be elucidated (Needham *et al*, 2019; Ochoa *et al*., 2020).

To date, only ~5% of the human phospho-proteome has been experimentally assigned to specific kinases (Jiang *et al*, 2025). A major advance in decoding kinase–substrate relationships emerged from the recent systematic annotation of kinases to their preferred ~90,000 phosphorylation sites, based on experimentally derived phosphorylation motifs covering approximately 84% of the human kinome (Johnson *et al*, 2023). This comprehensive Ser/Thr motif atlas now enables the high-resolution inference of activation or repression patterns across 303 Ser/Thr kinases directly from phospho-proteomic datasets (Johnson *et al*., 2023). In a recent landmark application, this approach delineated the regulatory kinase– substrate landscape in KRAS-mutant cancer cells exposed to an ERK MAP kinase inhibitor with unprecedented molecular resolution (Klomp *et al*, 2024). Comparable systematic studies of kinome perturbations in the context of viral infection are notably absent.

Coronavirus infection is characterized by the rapid reorganization of host subcellular architecture into specialized replicative organelles (ROs) that originate from the endoplasmic reticulum (ER) (Cortese *et al*, 2020; Zimmermann *et al*, 2023). This process is accompanied by activation of canonical stress kinase pathways, including JNK, p38, and NF-κB (Higgins *et al*, 2023; Mizutani *et al*, 2006; Mizutani *et al*, 2004, 2005; Neufeldt *et al*, 2022; Poppe *et al*, 2017). Recent work, including our own, has further shown that kinase-based ER sensors, most prominently the protein kinase RNA-dependent-like ER kinase (PERK), encoded by the eukaryotic translation initiation factor 2-alpha kinase 3 (*EIF2AK3*) gene, are strongly activated during coronavirus infection (Echavarria-Consuegra *et al*, 2021; Harding *et al*, 1999; Renner *et al*, 2024; Shaban *et al*, 2021). These findings suggest that ER stress acts as a central signaling hub that primes host cellular pathways at the earliest stages of infection.

Activated PERK is unique among eIF2α kinases in orchestrating both the unfolded protein response (UPR), an organelle-specific adaptive program that restores ER proteostasis, and the integrated stress response (ISR), which globally reprograms cellular biosynthetic capacity (Acosta-Alvear *et al*, 2025; Costa-Mattioli & Walter, 2020). The ISR generally acts to transiently stall translation under stress conditions, allowing recovery and adaptation (Karagoz *et al*, 2019). Translational arrest is mediated through phosphorylation of the translation initiation factor eIF2α by PERK or alternative eIF2α kinases, including PKR, HRI, and GCN2 (Hetz *et al*, 2020). Although these pathways are well-characterized, the downstream (patho)physiological consequences of UPR and ISR activation remain highly complex and incompletely understood (Marciniak *et al*, 2022; Urra & Hetz, 2017).

In this study, we systematically interrogated post-translational modifications (PTMs) and the regulation of the human kinome at transcriptional, translational, proteomic, and phospho-proteomic levels in cells infected with the human alpha coronavirus 229E (HCoV-229E) as a model system. By focusing on the consequences of pharmacological PERK inhibition, we reveal how a single kinase inhibitor simultaneously disrupts viral nucleocapsid (N) protein phosphorylation and modulates multiple host kinases, thereby rewiring the regulatory protein biosynthesis and phospho-proteomic landscape toward an antiviral state that counteracts virus-induced translational shutdown. The PERKi-sensitive CMGC kinase family emerges as particularly strongly regulated during infection, with this regulatory pattern conserved in SARS-CoV and SARS-CoV-2 infections. Comparative analyses with PERK-depleted cells delineate the on-target and off-target effects of PERKi on translation and kinase activation. These findings offer novel insights into the kinome-wide consequences of PERK perturbation in coronavirus-infected cells and highlight how the polypharmacology of PERK inhibitors can be strategically exploited to suppress RNA virus replication.

## Results

### Phosphorylation of HCoV-229E N protein links viral replication to host stress signaling

To dissect how host enzymes modify coronavirus proteins and shape the host response, we performed LC-MS/MS profiling of HCoV-229E–infected cells at 12 and 24 hours post infection (hpi). We detected extensive, partially overlapping acetylation, ubiquitination, and phosphorylation events across the host proteome as well as on 10 viral proteins, with the nucleocapsid (N) protein being the most extensively modified **(Fig. 1A, Supplementary Fig. 1A-B)**. Twenty-two phosphorylation(P)-sites were clustered within N, predicted to be targeted by more than 90 protein kinases spanning 12 kinase families **(Fig. 1B)**, highlighting the breadth of kinome involvement in the viral life cycle.

**Fig. 1.**
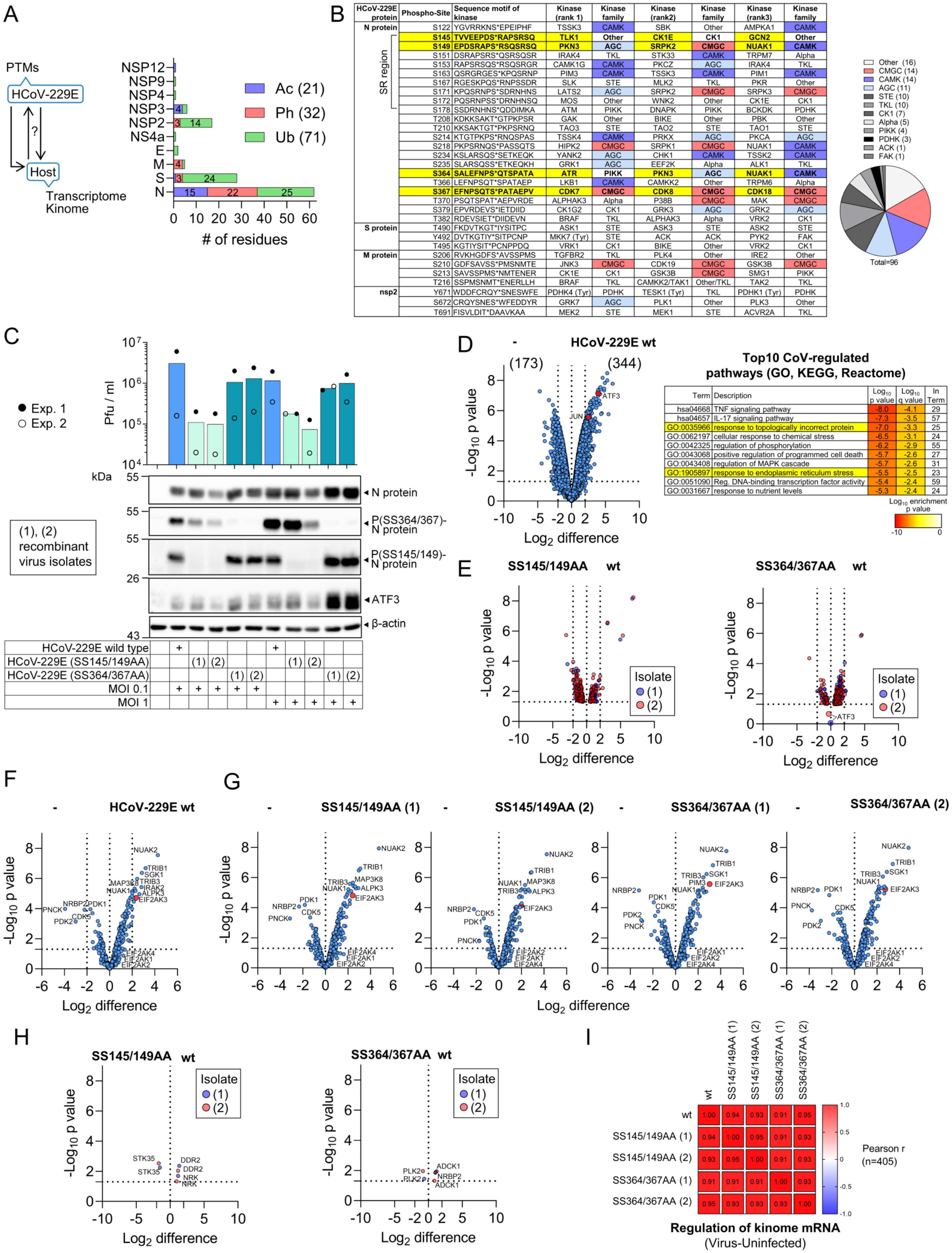
Regulation of the kinome transcriptome by wild type HCoV-229E and N protein phospho-site mutant viruses. (A) Left: Schematic summarizing the analysis of reciprocal effects of CoV protein posttranslational modifications on the HCoV-229E transcriptional host response. Right: Huh7 cells were infected for 12 h or 24 h with HCoV-229E at a multiplicity of infection (MOI) of 1 or were left untreated. Acetylated (Ac), ubiquitinylated (Ub) or phosphorylated (P) peptides were purified from trypsin-digested total cell extracts by anti acetyl or anti K-ε-GG antibodies or immobilized metal affinity columns (IMAC), respectively, and identified by LC-MS/MS. The bar graph summarizes the total number of modifications in CoV proteins detected at both time points post infection. (B) Table summarizing all centered phosphorylation sites of CoV proteins and the three most likely kinases (ranks 1-3) phosphorylating these sites according to motif enrichment analysis. The pie plot shows the kinase family distribution with CMGC, CAMK and AGC families highlighted in red, light or dark blue. Yellow colors highlight P-sites that were further analyzed using P-specific antibodies. (C) Huh7 cells were infected with recombinant HCoV-229E wild type or variants carrying tandem mutations in the middle (SS145/149AA) or C-terminal part (SS364/367AA) of the N protein. For each of the mutants two independent viral isolates (1, 2) were generated and used at low (0.1) or high MOI (1). The bar graph shows viral titers determined from supernatants 24 hpi of two independent experiments for both viral isolates. Whole cell extracts from the same cells were used for immunoblotting to monitor N protein or CoV-inducible ATF3 levels and the phosphorylation of N protein at SS145/149 or SS364/367 by double-purified polyclonal rabbit antibodies raised against di-phosphorylated peptides. Antibodies against β-actin were used to confirm equal loading. (D) Total RNA isolated from uninfected or HCoV-229E-infected cells (24 hpi, MOI=l) as shown in (C) was analyzed by microarrays carrying 60 k probes. All genes with annotated biotypes and a mean Log_2_ expression value of ≥ 4 were collected. The Volcano plot on the left shows mean Log_2_-fold changes and their significance from two biologically independent experiments. 344 or 173 genes were up- or down-regulated by more than 4-fold (LFC ≥ 2, -Log_10_ p value ≥ 1.3). The table on the right shows the top10 enriched pathway terms of all significantly differentially expressed genes (DEGs). Red dots highlight CoV-induced *ATF3* or *JUN* (encoding c-Jun) mRNA, while yellow colors highlight ER stress / UPR pathways, respectively. (E) Volcano plots of microarray results showing all DEGs (LFC ≥ or ≤ 0, -Log_10_ p value ≥ 1.3) from two biologically independent experiments comparing wild type HCoV-229E (wt) with P-site mutants from the infection experiments of isolates (1) and (2) (24 hpi, MOI=l) as shown in (C). (F-H) Volcano plots showing regulation of 405 protein kinases (out of 536) annotated within the kinhub data base (http://www.kinhub.org/index.html) (Eid *et al*., 2017) that were expressed at the mRNA level in wild type HCoV-229E (wt) (F), or in the P-site mutants compared to uninfected cells (G), or by directly comparing wild type CoV with mutants (H). The gene *EIF2AK3* encoding PERK is highlighted in red in (F-G). (I) Correlation matrix of CoV-induced changes in kinome expression compared to uninfected cells across all conditions described in (D) to (H). The heatmap shows pearson r.

We generated antibodies against tandem P-sites in the middle serine-rich region of the N protein, which is evolutionarily conserved in related α-coronaviruses, as well as against a largely unexplored and unique group of P-sites in the C-terminal region **(Supplementary Fig. 1C-D)**, and engineered recombinant HCoV-229E variants carrying alanine substitutions at either SS145/149 or SS364/367. Independent isolates of mutants with SS145/149 substitutions displayed reduced replication at both high and low multiplicity of infection (MOI), whereas SS364/367 mutants replicated comparably to wild type. Immunoblotting confirmed antibody specificity, loss of phosphorylation in mutants, and unexpectedly revealed elevated expression of ATF3, a highly consistently HCoV-229E-induced transcription factor downstream in the ER stress response with undefined roles in CoV replication, in cells infected with the SS364/367 variant **(Fig. 1C)**. These findings suggest that distinct P-sites in CoV proteins differentially regulate viral replication and host signaling.

Transcriptome analysis of infected cells revealed that wild-type HCoV-229E altered expression of 517 transcripts (> 4-fold at 24 hpi), including *ATF3* and *JUN* **(Fig. 1D)**. These strongly regulated transcripts mapped to stress-related, inflammatory and kinase pathways, particularly ER stress and the unfolded protein response (GO:0035966, GO:1905897), consistent with our prior analyses of HCoV-229E-infected Huh7 cells (Poppe *et al*., 2017). In contrast, infection with either N phospho-mutant caused minimal additional changes in host transcription **(Fig. 1E)**. Of the 536 human kinases annotated as human kinome in the kinhub data base (http://www.kinhub.org/)(Eid *et al*., 2017), 405 were expressed in Huh7 cells; 16 were strongly regulated (> 4-fold), including NUAK2, a kinase also regulated by SARS-CoV-2 (Prasad *et al*, 2023), and EIF2AK3 (encoding PERK), the latter being the sole differentially expressed member of the eIF2α kinase family **(Fig. 1F)**. PERK induction strongly implicated this kinase as the upstream driver for the activation of ATF3 and the ER stress / UPR pathways.

Comparison of wild-type and mutant virus infections revealed near-identical correlating kinase expression profiles, with only six kinases weakly altered **(Fig. 1G–I)**. Thus, N protein phosphorylation exerts selective effects on replication and ATF3 regulation without broadly altering kinome expression. Collectively, these data establish N phosphorylation as a functionally important viral modification and identify PERK as a key host kinase induced during HCoV-229E infection.

### Pharmacological inhibition of PERK suppresses N phosphorylation and viral replication

Given the induction of PERK during infection, we next examined whether pharmacological inhibition of this kinase perturbs coronavirus protein modifications, replication and signaling **(Fig. 2A)**. In line with our previous results, CoV infection at 12 hpi and 24 hpi, or short-term exposure to the prototypical ER stressor thapsigargin (Tg) activated PERK, as evidenced by its autophosphorylation causing retarded mobility upon SDS-PAGE, followed by increased serine 51 phosphorylation of the PERK substrate eIF2α **(Fig. 2B, Supplementary Fig. 2A)** (Shaban *et al*., 2021). Both events were abolished by the selective PERK inhibitor GSK2606414 (PERKi), confirming on-target activity **(Fig. 2B–C)**.

**Fig. 2.**
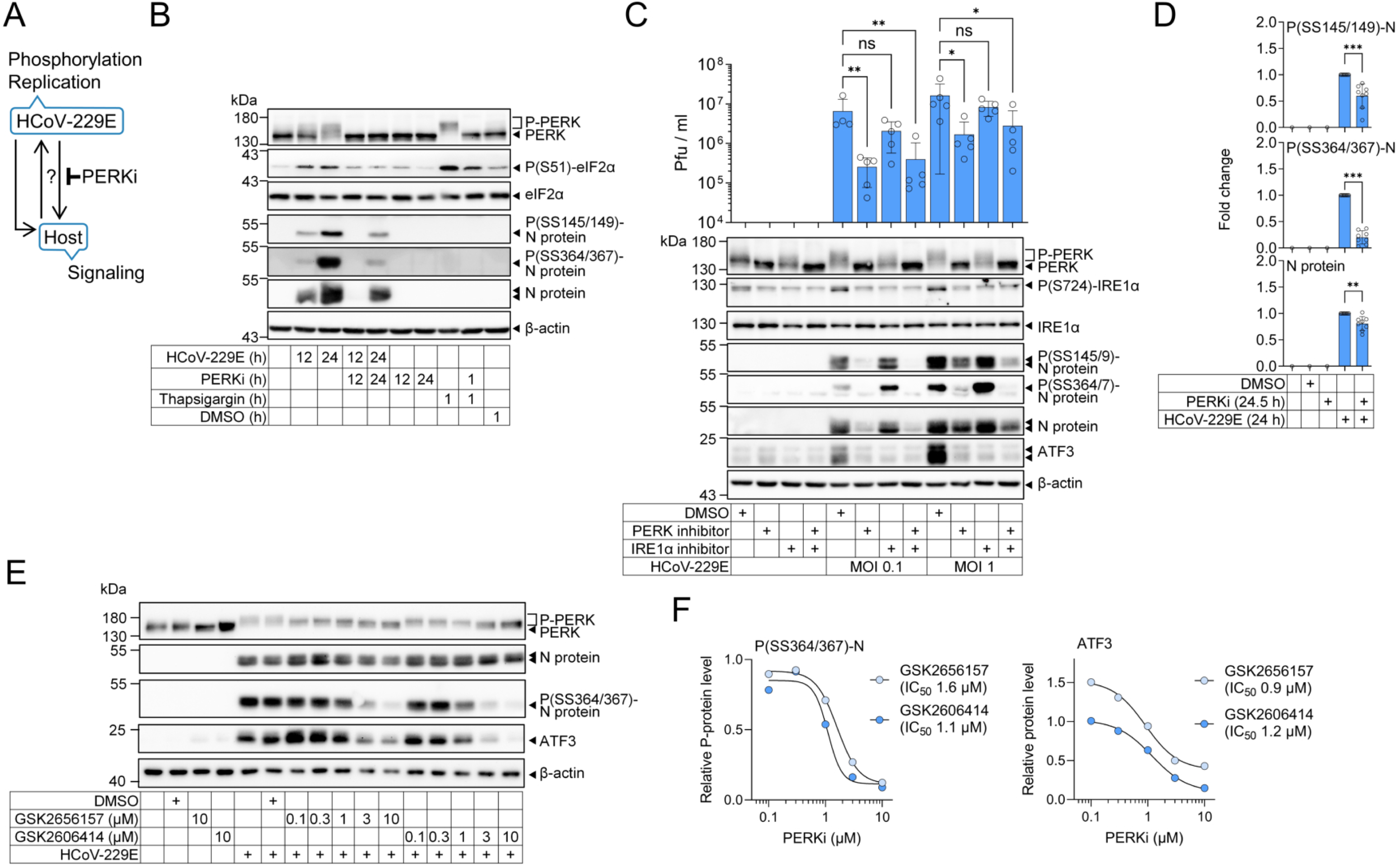
PERK inhibition suppresses N protein phosphorylation and viral replication. (A) Schematic summarizing the analysis of effects of PERK inhibitors (PERKi) on HCoV-229E protein modifications, replication and host signaling response. (B) Huh7 cells were left untreated or were pretreated for 30 minutes with PERK inhibitor (PERKi) GSK2606414 (10 µM) or solvent (DMSO) and then infected for 12 h or 24 h with HCoV-229E (MOI=l) or treated with thapsigargin (1 µM) for 1 h. Total cell extracts were analyzed for the phosphorylation or expression of the indicated proteins by Western blotting. Shown is one representative out of three independent experiments. (C) Huh7 cells were pretreated for 30 minutes with of PERK inhibitor GSK2606414 (10 µM), the IRE1α inhibitor KIRA6 (1 µM) or solvent (DMSO) and then infected for 24 h with HCoV-229E at MOI of 0.1 or 1 as indicated. Supernatants of cells were used to determine viral titers by plaque assays and whole cell extracts were used to analyze phosphorylation and expression of proteins by immunoblotting. The graph shows the mean titers ± s.d. from five biologically independent experiments, while the bottom panels show one representative out of five experiments. Asterisks indicate p values (*p ≤ 0.05, **p ≤ 0.01, ***p ≤ 0.001, ****p ≤ 0.0001) obtained by Kruskal Wallis tests. NS, not significant. (D) Summarizing quantification of PERKi effects on N protein expression and phosphorylation levels derived from data shown in (B-C) for cells infected with MOI of 1 in the 24 hpi condition. The graphs show relative mean changes ± s.d. from 8 independent experiments. Asterisks indicate p values (*p ≤ 0.05, **p ≤ 0.01, ***p ≤ 0.001, ****p ≤ 0.0001) obtained by Mann Whitney tests. (E) Huh7 cells were left untreated or were infected with HCoV-229E (MOI=l) for 24 h in the presence of increasing doses of PERK inhibitors GSK2606414 or GSK2656157 (0.1 to 10 µM) as indicated. The highest dose of each inhibitor and DMSO served as controls. Total cell extracts were analyzed for the phosphorylation or expression of the indicated proteins by Western blotting. Shown is one representative out of two independent experiments. (F) Relative levels of P(SS364/367)-N or ATF3 were quantified from the experiment shown in (E) and IC_50_ values were calculated from non-linear regression lines shown in the graphs.

PERKi treatment markedly reduced N protein expression and phosphorylation at 12 hpi, with stronger suppression of phosphorylation than protein abundance at 24 hpi **(Fig. 2B, Supplementary Fig. 2A)**. This effect was consistent across high and low MOI and translated into significantly reduced viral replication **(Fig. 2C, Supplementary Fig. 2B)**. By contrast, inhibition of IRE1α with KIRA6 did not affect N phosphorylation or replication, though both PERKi and KIRA6 attenuated CoV-induced ATF3 expression, highlighting distinct kinase-specific and shared ER stress–dependent regulatory axes **(Fig. 2C, Supplementary Fig. 2B)**. Notably, PERKi also suppressed IRE1α phosphorylation, indicating cross-talk between ER stress sensors, in line with (Chang *et al*, 2018; Ong *et al*, 2024).

Quantitative analyses from pooled data at 24 hpi revealed that PERKi reduced phosphorylation at N residues SS364/367 by 80% and at SS145/147 by 40%, while N protein expression declined by only ~20% **(Fig. 2D)**. These data suggest that PERKi disrupts both N kinase activity and viral protein synthesis. Dose-response experiments confirmed potent suppression of N phosphorylation, ATF3 induction, and replication by both GSK2606414 and the related compound GSK2656157 (Atkins *et al*, 2013; Axten *et al*, 2012; Axten *et al*, 2013; Shaban *et al*., 2021), with IC_50_ values of 1–2 µM and maximal inhibition at 10 µM **(Fig. 2E-F)**. Hence, this dose was chosen for subsequent experiments in our study.

Together, these findings demonstrate that PERK inhibition profoundly attenuates N phosphorylation, viral protein synthesis, and replication, establishing PERK as a critical host factor coupling coronavirus replication and N protein modifications to ER stress signaling.

### PERK inhibition restores coronavirus-suppressed translation

PERK is a key regulator of the integrated stress response and translation. We therefore asked whether the antiviral effects of PERK inhibitors were linked to regulation of the nascent proteome in HCoV-229E-infected cells.

To monitor protein synthesis, we labeled nascent polypeptides with puromycin or the alkyne analog O-propargyl-puromycin (OPP). Both incorporate into elongating peptide chains at the ribosomal peptidyl-transferase site, triggering premature release. Puromycin-labeled peptides can be detected in bulk by immunoblot, whereas OPP enables conjugation to biotin-azide via click chemistry in vitro and subsequent proteome-wide identification by LC–MS/MS (Aviner, 2020; Forester *et al*, 2018) **(Fig. 3A)**.

**Fig. 3.**
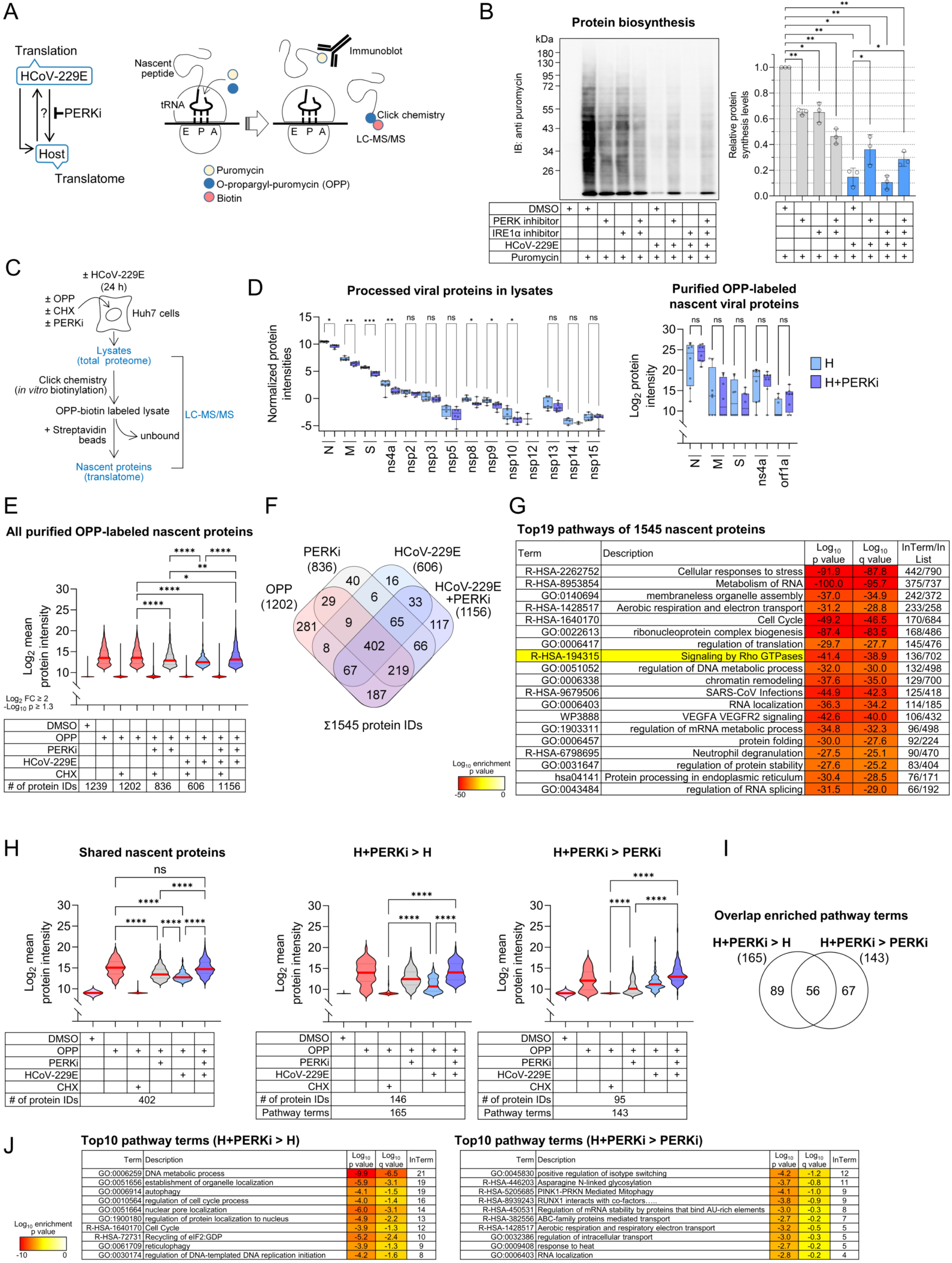
PERK inhibition antagonizes CoV-mediated translational shutdown. (A) Left: Schematic summarizing the analysis of effects of PERK inhibitors (PERKi) on translation of HCoV-229E and host proteins. Right: Cartoon visualizing the attachment of tyrosyl-tRNA mimetic puromycin or its derivative OPP to the C-terminal end of the growing polypepide chain during translational elongation, which enables monitoring bulk protein synthesis by anti puromycin antibodies or the identification of individual nascent polypeptides by biotin tagging, purification and mass spectrometry. (B) Huh7 cells were pretreated for 30 minutes with of PERK inhibitor GSK2606414 (10 µM), the IRE1α inhibitor KIRA6 (1 µM) or solvent (DMSO) and then infected for 24 h with HCoV-229E at MOI of 1. Puromycin (3µM) was added for the last 30 minutes as indicated and equal amounts of whole cell extracts were examined by immunoblotting using anti puromycin antibodies. The right panel shows one representative out of three immunoblot experiments. The graph shows mean levels of puromycinylation ± s.d. from three independent experiments. Asterisks indicate p values (*p ≤ 0.05, **p ≤ 0.01) obtained by one-way ANOVA. (C) Experimental design for assessing protein expression levels and the nascent proteome landscape by OPP metabolic labeling in CoV-infected cells. Huh7 cells were left untreated or were infected with HCoV-229E (MOI=1, 24 h) in the presence or absence of PERKi GSK2656157 (10 µM). For the last 2 h, cells were metabolically labeled with OPP (30 µM). Cycloheximide (CHX, 10 µg/ml) was added to one sample 1 h before OPP addition. All samples were lysed simultaneously and total cell extracts were subjected to click biochemistry using biotin azide as substrate. Biotinylated nascent polypeptides were purified by streptavidin affinity chromatography. Lysates and purified nascent protein fractions were digested by trypsin and peptides were identified and quantified by label-free LC-MS/MS analysis. (D) Protein intensity values derived from normalized cell lysates (left graph) or the nascent protein fractions (right graph) were analyzed for HCoV-229E structural (N, M, S), non-structural proteins (ns), translated and processed from open reading frame (orf) 1a, or accessory protein ns4a. Box plots show l^st^, 2^nd^ (median) and 3^rd^ quartiles and individual protein intensity values from four independent biological experiments and two technical replicates. Whiskers show minimum to maximum. NS, not significant. (E) Biotinylated and purified nascent proteins were identified from the conditions summarized in (C). For each quantified nascent protein, active translation was defined by the significant enrichment (LFC ≥ 2, -Log_10_ p value ≥ 1.3, student’s t-test) of mean protein intensity compared to values from CHX-treated cells, which represent the maximal translationally repressed state. As an additional negative control for unspecifically purified proteins, the OPP-labelled translatome of untreated cells (OPP only) was compared to the DMSO solvent control in the absence of OPP. Violin plots show the distribution of mean protein intensities of the indicated numbers of proteins in each fraction and the corresponding values in the presence of CHX from four independent biological experiments and two technical replicates. Asterisks indicate p values (*p ≤ 0.05, **p ≤ 0.01, ***p ≤ 0.001, ****p ≤ 0.0001) obtained by Kruskal Wallis tests. (F) Venn diagram showing the overlap of all 1545 nascent proteins identified across all conditions described in (E). (G) Top enriched pathway terms for the proteins shown in (F) alongside the number of identified components (In Term) of each pathway (In List). Yellow color highlights the Rho GTPase pathway. (H) Violin plots show the distribution of mean protein intensities of 402 actively translating proteins shared between all conditions according to the Venn diagram shown in (F) (left panel), or of nascent proteins sets which were significantly more enriched (-Log_10_ p value ≥ 1.3, student’s t-test) in PERKi-treated infected cells compared with infected cells only (middle panel), or with PERKi-treated cells only (right panel). Data are from four independent biological experiments and two technical replicates. Asterisks indicate p values (*p ≤ 0.05, **p ≤ 0.01, ***p ≤ 0.001, ****p ≤ 0.0001) obtained by Kruskal Wallis tests. (I-J) The Venn diagram shows overlapping and distinct pathway terms annotated to the nascent protein set upregulated in PERKi-treated infected cells compared to infected cells only, or PERKi-treated cells only, respectively (I). The tables in (J) show the top10 unique enriched pathway terms for each condition.

Anti puromycin antibodies confirmed that PERKi, but not KIRA6, partially rescued the coronavirus-driven translational shutdown. In contrast, in uninfected cells both compounds suppressed translation, either alone or in combination **(Fig. 3B)**.

To dissect how PERK inhibition alters translation, we enriched OPP-labeled nascent peptides by streptavidin affinity purification and quantified them by LC–MS/MS **(Fig. 3C, Supplementary Fig. 3)**. Most coronavirus structural and nonstructural proteins were reduced in total lysates upon PERKi treatment, however, their nascent peptide levels remained stable **(Fig. 3D).** This uncoupling indicates that PERKi does not block synthesis but instead impairs downstream maturation or stability of viral proteins.

We next benchmarked these nascent translation profiles against cycloheximide (CHX)-treated cells, where translational elongation is globally arrested by binding of CHX to the ribosomal 60S exit site (Dmitriev *et al*, 2020). Relative to this maximally repressed state, OPP labeling alone yielded ~1,200 enriched nascent proteins. This number fell sharply with PERKi or HCoV-229E infection (836 and 606 proteins, respectively), yet strikingly, PERKi treatment of infected cells restored both the number and intensity of nascent proteins to near-normal levels **(Fig. 3E)**.

Across all conditions, we identified 1,545 unique proteins, with only 25% (402) shared, underscoring extensive remodeling of the translational landscape under PERK inhibition, viral infection, or both **(Fig. 3F)**. Pathway analysis revealed enrichment in processes central to stress and infection, including RNA transcription, metabolism and localization, translation and folding, membraneless organelle assembly, energy metabolism, cell-cycle control, Rho GTPase signaling, and SARS-CoV-2 infection **(Fig. 3G)**.

Focusing on the 402 common nascent proteins, we observed broad suppression by both PERKi and HCoV-229E, with translation derepressed only when PERKi was applied to infected cells **(Fig. 3H, left)**. This pattern paralleled global protein synthesis measured by puromycin incorporation **(Fig. 3B)**. Two additional subsets of nascent proteins were selectively enriched: one induced only by PERKi in infected cells (165 proteins) and another enhanced by infection plus PERKi relative to PERKi alone (143 proteins) **(Fig. 3H, middle and right)**. These subsets mapped to distinct functional signatures: PERKi in infected cells preferentially upregulated proteins linked to nuclear processes, autophagy, cell-cycle regulation, and eIF2:GDP recycling, whereas in uninfected cells PERKi primarily affected proteins involved in N-glycosylation, mitochondrial function, and mRNA stability **(Fig. 3I–J)**.

Together, these findings show that PERK inhibition not only suppresses basal translation but paradoxically restores translation of large subsets of proteins silenced by HCoV-229E. This dual activity was unlikely to be explained solely by partial blockade of virus-induced eIF2α phosphorylation and instead pointed to additional mechanisms of drug action beyond the classical ISR.

### Proteome- and kinome-wide effects of PERK inhibition in CoV-infected cells

Although PERK itself was not predicted to directly phosphorylate the coronavirus N protein **(Fig. 1B)**, the ability of PERK inhibition to suppress multiple viral phenotypes prompted us to map the downstream networks that mediate these effects. We first profiled the proteomes of HCoV-229E–infected cells at 12 and 24 hpi, with or without PERK inhibition.

Infection alone induced widespread changes: 670 proteins were upregulated and 136 downregulated by at least twofold, with many more altered at lower thresholds. Viral structural proteins and nonstructural proteins dominated the most strongly induced group **(Fig. 4A–B)**. Notably, the overlap of differentially expressed proteins (DEPs) between 12 hpi and 24 hpi was minimal, underscoring continuous proteome remodeling during ongoing infection **(Fig. 4A–B)**. Enrichment analysis revealed time- and virus-specific regulation of pathways linked to membrane organization, vesicle transport, mitochondrial biology, transcription, lipid and carboxylic acid metabolism, cell-cycle control, and ER stress **(Supplementary Fig. 4)**.

**Fig. 4.**
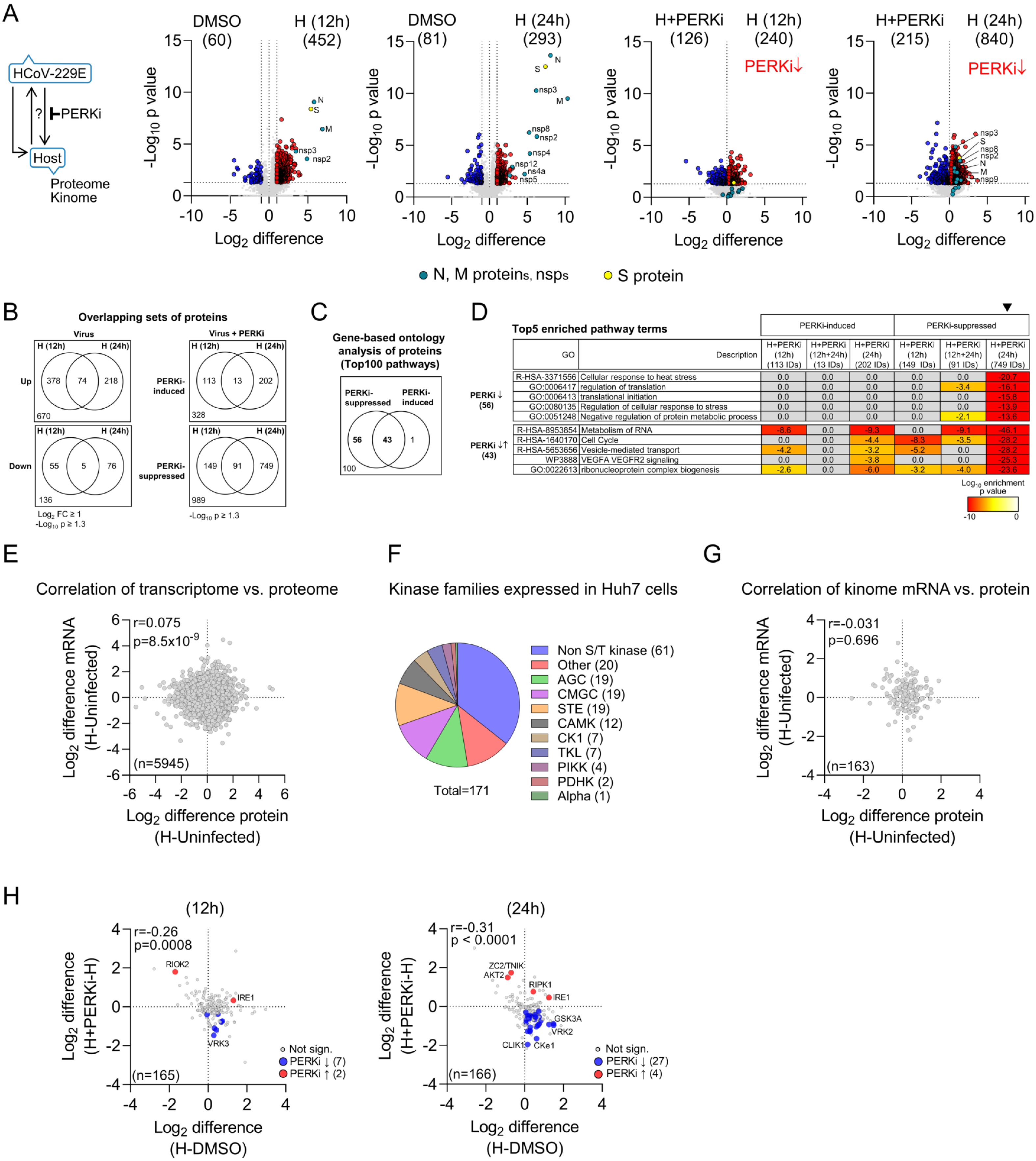
Protein level regulation of the HCoV-229E host response and the kinome by inhibition of PERK. (A) Huh7 cells were infected for 12 h or 24 h with HCoV-229E (H, MOI=l) and treated with of PERK inhibitor GSK2656157 (10 µM) or solvent (DMSO). Proteome analysis was performed from whole cell extracts identifying 6488 protein IDs. Volcano plots show mean Log_2_-fold changes and their significance from four biologically independent experiments and two technical replicates. Numbers in brackets and red or blue colors indicate up- or down-regulated proteins in response to CoV (LFC ≥ 1, - Log_10_ value ≥ 1.3) or PERKi treatment (-Log_10_ p value ≥ 1.3, student’s t-test). Individual CoV proteins and protein sets suppressed by PERKi treatment in infected cells are indicated. (B) Venn diagrams showing the overlap of proteins differentially expressed in response to CoV or PERKi. (C) Overrepresentation analysis was performed side by side for the six groups of proteins that were up- or down-regulated in response to PERKi after 12 h or 24 hpi as shown in (B). The Venn diagram shows the overlap of the top100 enriched pathway terms across all conditions. (D) Heatmap showing the top5 most strongly enriched pathway terms representing exclusively suppressed protein sets or those with a mixed pattern of regulation in response to PERKi. Pathways are sorted according to enrichment values marked by the arrowhead. (E) Correlation analysis of the regulation of the mRNA versus the protein levels of all 5945 factors with both mRNA and protein measurements in response to HCoV-229E (H) 24 hpi. Coefficients of correlation (spearman r) and p values for significance are indicated. (F) Family distribution of 171 out 536 protein kinases detected by LC-MS/MS. The projection of all kinases on the entire kinome tree is shown in Supplementary Fig. 5. (G) Correlation analysis of the regulation of mRNA versus the protein levels of 163 kinases with both mRNA and protein measurements in response to HCoV-229E (H). Coefficients of correlation (pearson r) and p values for significance are indicated. (H) Correlation of changes of protein kinase levels in HCoV-229E-infected cells (H-DMSO) or infected cells treated with PERKi (H+PERKi-H) 12 hpi or 24 hpi. Coefficients of correlation (spearman r) and p values for significance are indicated. Blue or red colors indicated significant differences in ratio values (-Log_10_ p value ≥ 1.3, student’s t-test).

PERK inhibition superimposed an additional layer of regulation, suppressing 989 proteins (–log10 p ≥ 1.3) while inducing 328 others, including both host and viral proteins **(Fig. 4A–B)**. These proteins mapped to 100 enriched pathways, with 43 shared between up- and downregulated sets across both time points **(Fig. 4C)**. At 24 hpi, PERKi most strongly suppressed pathways associated with translation, stress responses, and metabolism—paralleling the reversal of CoV-driven translational shutdown observed in **Fig. 3**. Conversely, pathways related to RNA metabolism, vesicle trafficking, and cell-cycle control contained both induced and suppressed proteins, suggesting a remodeling process that enforces an antiviral state **(Fig. 4D)**.

Transcript–protein comparisons revealed an unexpected disconnect: among 5,945 matched mRNA– protein pairs, there was essentially no correlation during infection **(Fig. 4E)**. The kinome subset mirrored this uncoupling. Of the 6,488 proteins detected, 171 were kinases spanning 11 families **(Supplementary Fig. 5)**. However, mRNA and protein levels for 167 matched kinase pairs were also uncorrelated **(Fig. 4G)**.

Despite this transcription–translation uncoupling, PERKi exerted additional effects on the kinome proteome in infected cells. After 24 h of treatment, 27 kinases were progressively downregulated, while only four were induced **(Fig. 4H)**.

Together, these proteome- and kinome-wide analyses reveal that PERK inhibition exerts broad effects on host–virus networks. Beyond counteracting the coronavirus-induced translational block, PERKi attenuates expression of more than two dozen kinases critical for the infected cell state, highlighting PERK as a central node in shaping both viral replication and host signaling.

### PERK inhibition rewires the phospho-proteome to disrupt CoV replication

To determine how PERK inhibition alters phosphorylation during coronavirus infection, we profiled 12,585 IMAC-enriched phospho(P)-peptides from the same samples used for proteomics. In line with immunoblotting **(Fig. 2)**, LC–MS/MS confirmed phosphorylation of the N protein at S145, which was blocked by PERKi at 12 hpi **(Fig. 5A)**. Additional N protein P-sites (S192, S208) and S protein phosphorylation (S210) were also suppressed, albeit with distinct temporal dynamics. Five further phosphorylation clusters were identified in N protein, though peptides containing SS364/367 were undetected **(Fig. 5A)**.

**Fig. 5.**
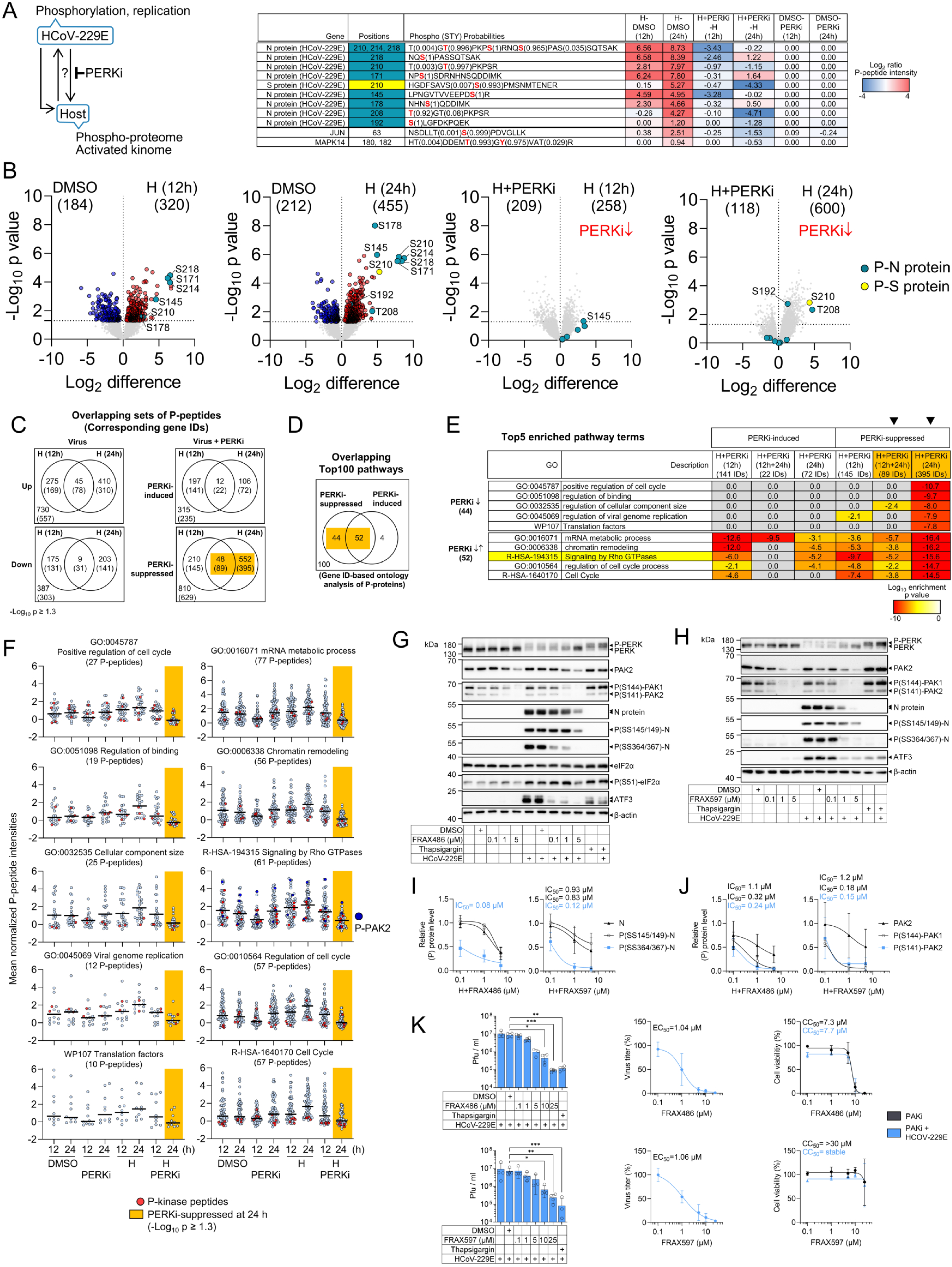
Widespread effects of PERK inhibition on the HCoV-229E-regulated phospho-proteome. (A) P-peptides from the same samples described in Fig. 4A were extracted by IMAC and analyzed by LC-MS/MS resulting in 12585 peptide IDs. The heatmap demonstrates mean changes of P-peptides, their sequences and the identified P-sites (in red) corresponding to the N or S proteins (green or yellow colors) and to host proteins c-Jun and p38 MAPK (MAPK14) for each of the six conditions analyzed. Data are from four biologically independent experiments and two technical replicates. (B) Volcano plots show mean Log_2_-fold changes of all P-peptides and their significance from four biologically independent experiments and two technical replicates. Numbers in brackets and red or blue colors indicate up- or down-regulated P-peptides in response to CoV or PERKi treatment (-Log_10_ p value ≥ 1.3, student’s t-test). P-peptides corresponding to N or S protein and individual P-sites are highlighed in green or yellow, respectively. Red text indicates sets of P-peptides suppressed by PERKi treatment in infected cells. (C) Venn diagrams indicating the overlap of P-peptides differentially phosphorylated in response to CoV or PERKi based on (-Log_10_ p value ≥ 1.3, student’s t-test). Numbers in brackets indicate the overlap of gene IDs corresponding to the identified P-peptides. (D) Overrepresentation analysis was performed side by side based on gene IDs corresponding to P-peptides for factors that were up- or down-regulated in response to PERKi after 12 h or 24 hpi. The Venn diagram shows the overlap of the top100 enriched pathway terms across all conditions. (E) Heatmap showing the top5 most strongly enriched pathway terms representing exclusively suppressed P-protein sets (upper table) or those with a mixed pattern of regulation in response to PERKi (lower table). Yellow color highlights the Rho GTPase pathway. (F) Scatter plots showing mean abundancies of the P-peptides that are suppressed by PERKi 24 hpi, segregated according to the pathways identified in (E), along with their values across all conditions. Data are from four biological replicates and two technical replicates, black lines indicate medians. Red colors mark P-peptides belonging to protein kinases. Blue colors highlight P-peptides of PAK2. (G-J) Huh7 cells were infected for 24 h with HCoV-229E (H, MOI=1) and treated with group I PAK inhibitors FRAX469 (G, I) or FRAX597 (H-J) at the indicated concentrations, thapsigargin (1 µM), solvent (DMSO) or were left untreated. (G-H) Whole cell extracts were used to analyze phosphorylation and expression of proteins by Western blotting. One representative out of four biologically independent experiments is shown. (I-J) Quantification of the indicated (P)-proteins from four replicates. Shown are mean values ± s.d. including non-linear regression lines used to estimate IC_50_ values. (K) Experiments were performed as described in (G-J), including two additional concentrations of PAKi (10 µM, 25 µM). Supernatants of cells were used to analyze viral titers by plaque assays and parallel cell cultures were used to assess cell viability. Left graphs show mean titers ± s.d. from four biologically independent experiments. The same data were used to estimate EC_50_ values by non-linear regression (middle graphs). Right graphs show mean normalized cell viability values ± s.d. including non-linear regression lines to estimate CC_50_ values as indicated, obtained from three biologically independent experiments. Asterisks indicate p values (*p ≤ 0.05, **p ≤ 0.01, ***p ≤ 0.001) derived from Kruskal Wallis tests. (C-F) Amber rectangles denote the sets of P-peptides and their corresponding gene IDs suppressed significantly by PERKi at 24 hpi (-Log_10_ p value ≥ 1.3, student’s t-test).

Host factors mirrored these effects. CoV-induced phosphorylation of c-Jun (S63) and p38 MAPK (T180/Y182) was strongly attenuated by PERKi **(Fig. 5A)**.

Globally, infection upregulated 730 and downregulated 387 P-peptides, corresponding to 557 and 303 proteins, respectively. These factors mapped to multiple pathways involved in chromatin remodeling and transcription, metabolism of RNA and translation, protein localization to organelles and Rho GTPase signaling and significantly overlapped with P-proteome studies of SARS-CoV-2-infected cells (Supplementary Fig. 6A-B).

PERKi inverted this balance, downregulating 810 and upregulating 315 phospho-peptides, corresponding to 629 and 235 proteins, respectively **(Fig. 5B–C)**.

Pathway enrichment revealed a clear bias: Among the top 100 enriched terms, 96 were linked to downregulated P-proteins after PERKi treatment, 52 showed mixed regulation, and only 4 were upregulated, revealing a broadly suppressive effect of PERKi on host phosphorylation **(Fig. 5D)**.

At 24 hpi, PERKi most strongly suppressed phosphorylation of specific subsets of proteins linked to cell-cycle progression, general terms (binding, cell size), viral replication and translation, supporting its antiviral phenotype shown in **Fig. 2 (Fig. 5E, Supplementary Fig.7)**. In contrast, pathways associated with RNA metabolism, chromatin remodeling, and Rho GTPase signaling contained both up- and downregulated substrates, suggesting a rewiring of host phosphorylation networks that culminates in an antiviral state **(Fig. 5E)**.

Accordingly, the Top10 pathway–resolved pattern of PERKi-suppressed P-peptides at 24 hpi **(highlighted in amber in Fig. 5C–F)** revealed a progressive, virus-driven increase in phosphorylation alongside an early suppression of basal phosphorylation by PERKi **(Fig. 5F)**. This individual P-peptide level high-resolution view underscores that PERKi exerts distinct effects on basal versus virus-induced cellular states, a pattern also evident in the analysis of nascent protein synthesis **(Fig. 3)**.

Notably, nearly all affected pathways contained kinase-derived P-peptides, pointing to upstream regulatory nodes directly sensitive to PERKi. Rho GTPase signaling (R-HSA-194315) contained most P-kinase peptides, including three ones representing p21-activated protein kinase (PAK) 2 **(Fig. 5F)**.

Independent pharmacological inhibition of PAKs with group I PAK inhibitors FRAX486 or FRAX597 confirmed this prediction: both compounds dose-dependently suppressed PAK phosphorylation at a critical serine residue activated by small GTPases (S141 in PAK2; S144 in PAK1) in non-infected and CoV-infected cells, inhibited PERK activation, reduced N protein expression and phosphorylation, and blocked ATF3 induction **(Fig. 5G–H)**. Thapsigargin, by contrast, did not affect PAK phosphorylation, highlighting a distinct PERK-dependent antiviral pathway **(Fig. 5G–H)**.

SS364/367 N protein phosphorylation was most strongly suppressed by PAK inhibition with an estimated IC_50_ of ~0.2 µM, while reduction of N protein, P(SS145/149)-N and PAK2 protein levels required considerably higher doses **(Fig. 5I, J)**.

PAK inhibition also potently suppressed viral replication (IC_50_ ~1 µM), with FRAX486 (CC_50_ ~7 µM) showing greater cytotoxicity than FRAX597 (CC_50_ > 30 µM) **(Fig. 5K)**.

Together, these results reveal that PERK inhibition broadly suppresses coronavirus-induced phosphorylation, including regulatory sites on viral proteins and host kinases. The discovery that PAK activity is a critical downstream effector in this network provides mechanistic insight into how PERKi exerts unconventional antiviral effects.

### Kinome motif analysis identifies CMGC family kinases as major PERKi targets

Because direct validation of kinase–substrate relationships remains technically limited, we used the recently published human kinome motif atlas to infer kinase activities from our phospho-proteome data. Of 12,585 P-peptides, 10,952 (87%) matched the individually phosphorylated residue as well as its surrounding ± 7 amino acids, confirming the robustness of our dataset **(Fig. 6A)**.

**Fig. 6.**
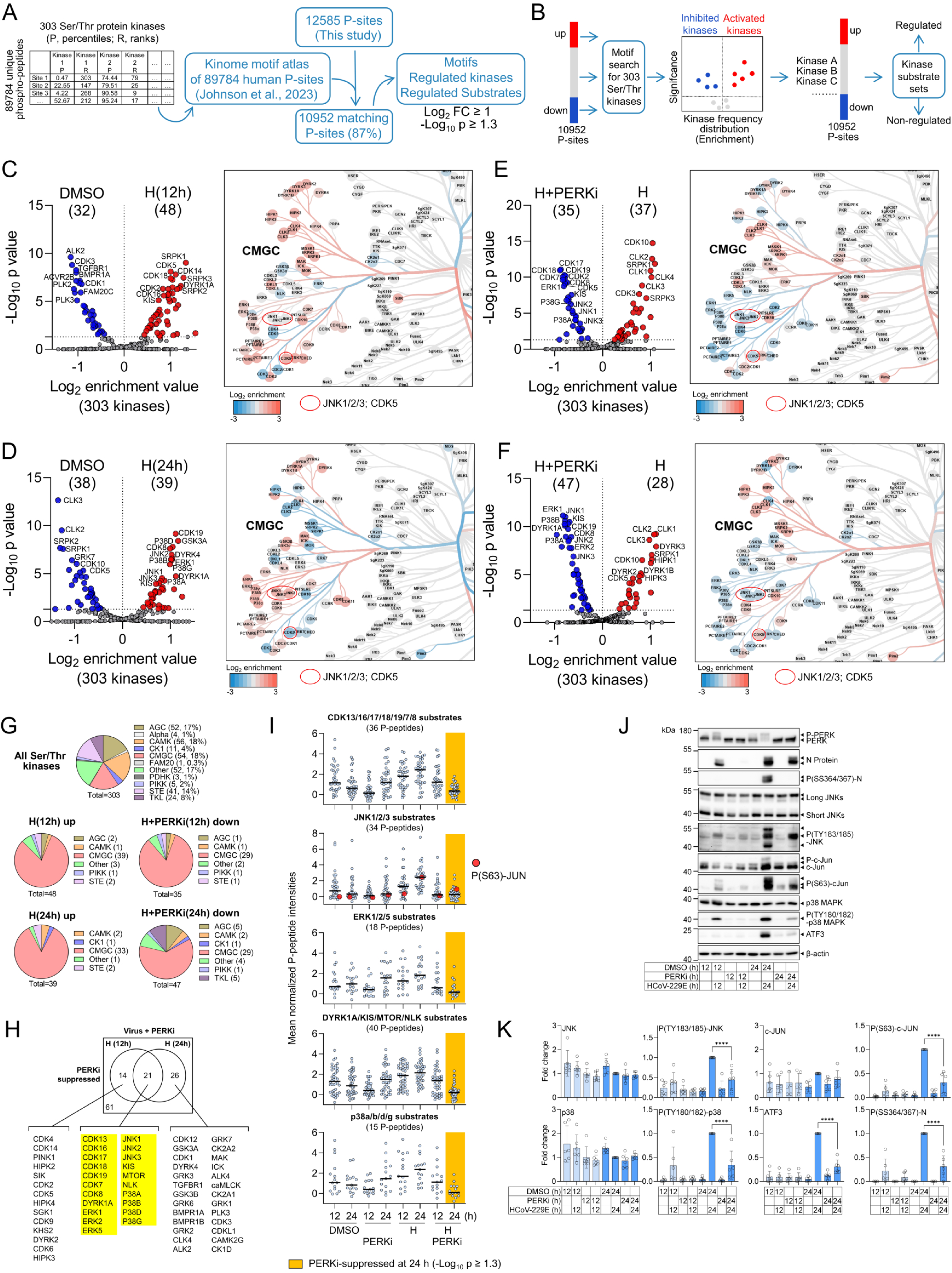
PERK inhibition antagonizes HCoV-299E-mediated activation of multiple CMGC kinases and the phosphorylation of specific sets of their substrates. (A) Schematic showing the integration of data from the human kinome motif atlas including the experimentally derived P-motifs of 303 Ser/Thr kinases with the P-proteome data sets of this study resulting in the identification of 10952 matching P-motifs and the associated ranked list of Ser/Thr kinases phosphorylating these P-sites. (B) Strategy to infer enriched (i.e. activated) or depleted (i.e. inactivated) kinases in CoV-infected or PERKi-treated cells by determining the frequency distribution of motifs and their associated kinases in up- or down-regulated compared to non-regulated, background P-peptide sets using centered motifs (P- site ± 7 amino acids), mean Log_2_-fold changes of P-peptide abundancies and their significance of all 10952 P-peptides as input data. Regulated P-peptides were defined by Log_2_ FC ≥ 1 and -Log_10_ p value ≥ 1.3. Data are from four biological replicates. (C-F) Volcano plots showing the time-wise relative enrichment / activation (red) or depletion / inhibition (blue) of 303 Ser/Thr kinases and the significance in HCoV-229E-infected cells (H) compared to solvent-treated control cells (DMSO) (C, D), or the effects of PERK inhibition in infected cells (E, F). The inserts show the projection of enrichment values on the CMGC part of the kinome tree using Coral software (Metz *et al*., 2018). Red circles mark JNKl/2/3 and CDK5 kinases as examples of differently regulated kinases. The projection of all regulated kinases on the entire kinome tree is shown in Supplementary Fig. 9. (G) Pie charts showing proportional changes of regulated kinase families and their kinetics in CoV- infected and PERKi-treated cells as derived from P-motif analyses shown in (C-F). The upper diagram depicts the principal family distribution of the set of 303 Ser/Thr kinases. Numbers in brackets indicate kinases and percentage per family. (H) Overlapping and distinct sets of 61 Ser/Thr kinases downregulated by PERKi in CoV-infected cells at 12 h compared to 24 h based on the analyses shown in (C-G). Yellow color highlights kinases chosen for further analysis of substrates. (I) Scatter plots showing mean abundancies of the P-peptide substrates that are suppressed by PERKi 24 hpi grouped into five sets representing the 21 kinases that are consistently suppressed by PERKi at 12 hpi and 24 hpi as shown in (H). For each peptide, the topl-ranked Ser/Thr kinase was selected and the values for all conditions are shown. Data are from four biological replicates, black lines indicate medians. Amber colors highlight the sets of P-peptides suppressed significantly by PERKi at 24 h p.i. (-Log_10_ p value ≥ 1.3, student’s t-test). (J-K) Huh7 cells were infected for 24 h with HCoV-229E (H, MOI=l) and treated with PERKi GSK2656157 (10 µM), solvent (DMSO) or were left untreated. (K) Whole cell extracts were used to analyze phosphorylation and expression of proteins by immunoblotting. One representative out of six biologically independent experiments is shown. (J) Bar graphs show mean relative changes ± s.d. of (P)- protein levels compared to the 24 h infection condition from six independent experiments. Asterisks indicate p values (**** p ≤ 0.0001) derived from one-way ANOVA.

As outlined in **Fig. 6B**, the overlap of our P-site datasets enabled the identification of regulated kinase activities for 303 Ser/Thr kinases in response to CoV infection and PERKi treatment.

The only well characterized direct substrates of PERK are Ser51 of eIF2α and PERK itself (Marciniak *et al*, 2006). However, the entire human consensus P-proteome contains > 300 P-sites with PERK listed as rank1 kinase, leaving the possibility that activated PERK has as yet undiscovered direct substrates beyond P-eIF2α **(Supplementary Fig. 8A)**. Modified peptides from the four eIF2α kinases PERK, GCN2, PKR, HRI and eIF2α itself were not detected in our P-proteomic data sets, likely due to low abundance. In total, we identified 21–54 motifs assignable to these kinases in the Huh7 phospho- proteomes, which were not reproducibly regulated across conditions, suggesting that PERK substrates are underrepresented in the Huh7 P-proteome and kinases beyond PERK dominate the (measurable) phosphorylation landscape **(Supplementary Fig. 8B–C)**

Systematic motif and kinase enrichment analyses of all 10,952 P-peptides revealed 28–48 kinases activated or inhibited across conditions **(Fig. 6C–F)**. As exemplified by Volcano plots and projection onto the human kinome tree, CMGC family members were strongly enriched and displayed dynamic regulation: CDK5 was upregulated at 12 hpi but downregulated at 24 hpi, whereas JNK1–3 were induced only at 24 hpi. PERKi reversed many of these virus-induced changes **(Fig. 6C–F, Supplementary Fig. 9)**.

The 303 Ser/Thr kinases distribute to 11 protein kinase families, of which 8 were modulated by CoV infection **(Fig. 6G)**. Remarkably, CMGC family kinases, which account for 18% of the 303 kinases tested, made up > 80% of all activated or inhibited kinases at 12 h and 24 h after infection **(Fig. 6G)**. PERKi had little effect on this distribution at 12 hpi but shifted it at 24 hpi, increasing contributions from TKL and miscellaneous kinases **(Fig. 6G)**.

In total, 61 kinases were suppressed by PERKi in infected cells, including multiple CDKs, JNKs, p38 MAPKs, ERKs, and a diverse set of others (DYRK1A, KIS, mTOR, NLK) **(Fig. 6H)**. At 24 hpi, PERKi strongly suppressed CoV-induced phosphorylation of substrates from these kinase groups, including c-Jun at P(S63), a canonical JNK target **(Fig. 6I)**. The specificity of our motif-based approach was validated by the presence of numerous unchanged substrates from proline-directed kinases with similar sequence preferences **(Supplementary Fig. 10)**.

Consistent with these phospho-proteomic results, immunoblotting confirmed that PERK inhibition suppresses CoV-induced phosphorylation of multiple JNK isoforms, c-Jun (P-S63), and p38 MAPK, while also blocking PERK autophosphorylation and ATF3 induction **(Fig. 6J–K)**.

Thus, motif-based analysis uncovered a PERKi-sensitive signaling network dominated by CMGC kinases providing a mechanistic framework for how PERKi exerts broad antiviral effects.

### PERK ablation identifies on and off-target effects of PERK inhibitors

To reveal whether PERKi effects were specific, or were affecting the activation of stress kinase systems during the infection cycle, with the inhibitor binding to the ATP pocket of additional kinases, we investigated key parameters of our analyses in genome-edited PERK-deficient cells.

PERK suppression reduced short-term Tg-inducible eIF2α phosphorylation strongly (from nine- to twofold), consistent with data from PERK-deficient murine embryonic fibroblasts (Harding *et al*, 2000b) **(Supplementary Fig. 11A)**. In contrast, long-term Tg treatment with or without CoV infection for 24 h, or, as a control, proinflammatory cytokine stimulation with IL-1α, induced eIF2α phosphorylation less strongly, with eIF2α phosphorylation only partially reduced in cells lacking PERK, suggesting that in these conditions alternative eIF2α kinases such as GCN2 contribute to the induction or maintenance of the ISR (Harding *et al*, 2000a) **(Supplementary Fig. 11A)**.

In the absence of PERK or IRE1α, viral titers were significantly reduced, but less strong as with PERKi **(see Fig. 2C)** or with Tg, the latter causing complete suppression of replication via suppression of additional pathways **(Supplementary Fig. 11B)**.

In line with the effects on P(S51)-eIF2α, lack of PERK alleviated the short-term Tg-mediated translational shutdown completely, again consistent with previous studies (Harding *et al*., 2000b) **(Supplementary Fig. 11C-D)**. Absence of PERK had no global effect on protein biosynthesis in basal or IL-1α-regulated conditions. However, CoV-mediated translational shutdown in the absence or presence of additional Tg treatment was reduced, indicating the requirement of activated PERK for the CoV-induced ISR **(Supplementary Fig. 11C-D)**.

These data suggested that PERKi-mediated modulation of the translational landscape occuring through on-target effects is a substantial part of the overall antiviral effects.

In contrast, the inhibitory effects of PERK on CoV-induced phosphorylation of JNK, p38 MAPK and c-Jun were largely maintained in PERK-deficient cells, demonstrating off-target activity on key steps of CMGC kinome activation that additionally contribute to the antiviral state **(Supplementary Fig. 12A-B)**.

### Meta-analyses across coronaviruses reveal a conserved CMGC kinase signature

To assess whether the PERKi- and kinase-dependent regulation observed in HCoV-229E extended to highly pathogenic coronaviruses, we reanalyzed published phospho-proteomic datasets from A549-ACE2 cells infected with SARS-CoV or SARS-CoV-2 (Stukalov *et al*, 2021). At the proteome level, A549-ACE2 cells expressed a similar number and family distribution of kinases as Huh7 cells **(Fig. 7A)**. Among the 183 expressed kinases, 18 to 46 were regulated by SARS-CoV or SARS-CoV-2 **(Fig. 7B)**. However, CoV infection remodeled the phospho-proteome far more extensively: >2,500 sites changed upon infection, confirming that the major regulatory layer resides in kinase activity rather than protein abundance **(Fig. 7B)**.

**Fig. 7.**
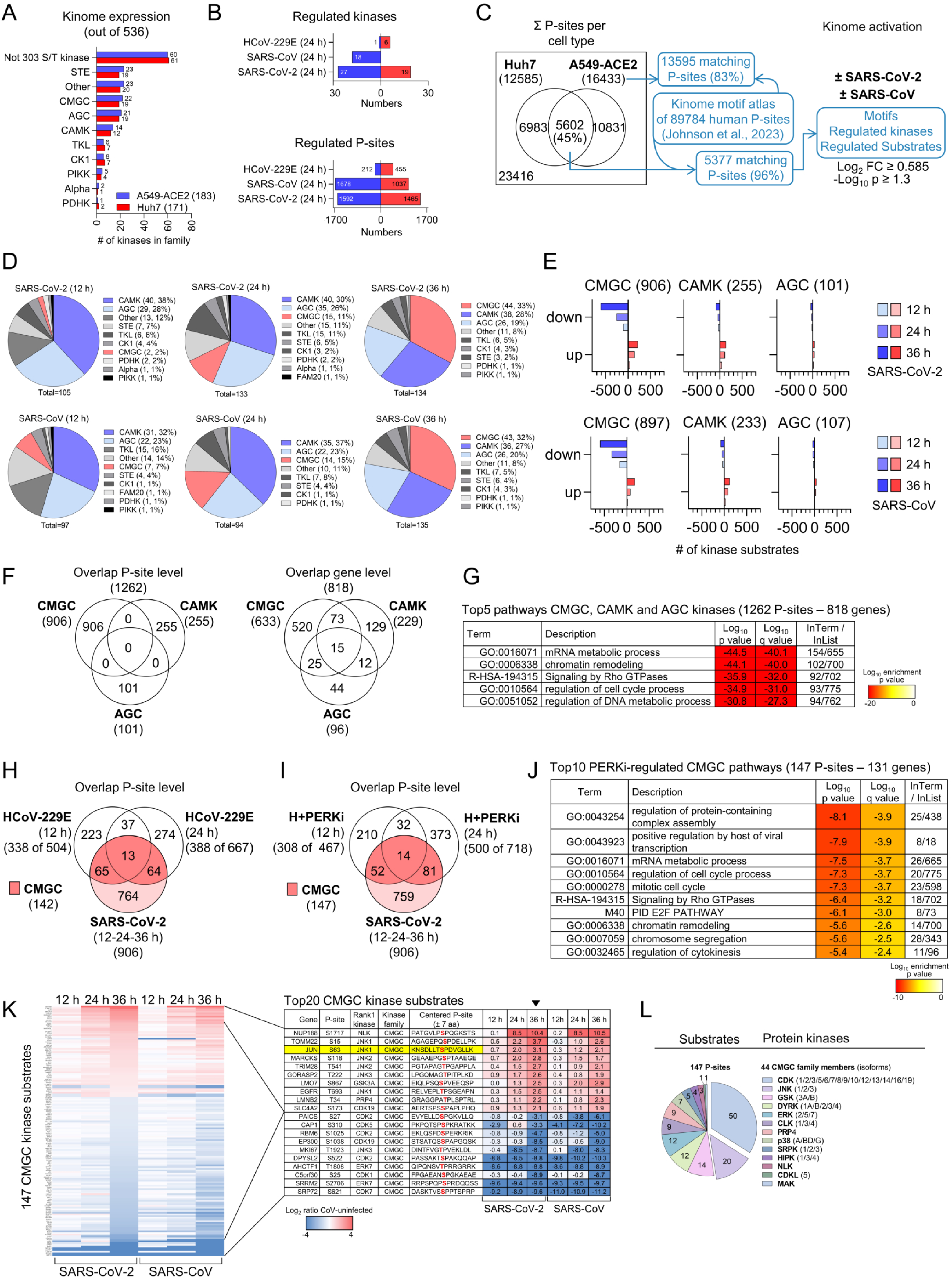
Identification of a PERKi-sensitive conserved set of CoV-regulated CMGC kinases and their substrates. (A-C) Published (phospho)-proteome data from A549-ACE2 cells infected with SARS-CoV or SARS-CoV-2 for 12 h, 24 h and 36 h were downloaded from the supplement of Stukalov et al. (Stukalov *et al*., 2021). Data were derived from four biological replicates. Data concerning Huh7 cells are from this study. (A) Proteomic data were searched for the number and family distribution of protein kinases expressed in A549-ACE2 cells compared with Huh7 cells from this study using the kinhub data bases of 536 kinases. (B) Upper bar graphs show numbers of kinases regulated at protein level (difference > or < 0, -Log_10_ p value ≥ 1.3) and lower bar graphs show differentially regulated P-sites in A549-ACE2 cells infected with SARS-CoV or SARS-CoV-2 for 24 h (LFC ≥ 0.585, -Log_10_ p value ≥ 1.3) compared to values from Huh7 cells infected with HCoV-229E for 24 h in this study. (C) The Venn diagram shows distinct and overlapping sets of P-proteins identified in Huh7 and A549-ACE2 cells. The joint set of P-peptides matching the human kinome motif atlas (5377 P-sites) was used as input data to perform kinome enrichment analysis of infected A549-ACE2 cells. Regulated P-peptides were defined by LFC ≥ 0.585 and -Log_10_ p value ≥ 1.3. (D) Pie charts showing proportional changes of regulated Ser/Thr kinase families and their kinetics in SARS-CoV / SARS-CoV-2-infected A549-ACE2 cells derived from P-motif analyses shown in (C). Numbers in brackets indicate number of identified kinases and percentage per family. (E) For each P-peptide of A549-ACE2 cells, the topl-ranked Ser/Thr kinase was selected. Bar graphs show the total numbers of unique regulated P-sites (LFC ≥ 0.585 and -Log_10_ p value ≥ 1.3) for the three most strongly regulated CMGC, CAMK and AGC kinase families according to time point of infection and virus. (F, G) Venn diagrams show the overlap of P-peptides pooled from all time points of infection or of the corresponding genes and the heatmap depicts the top5 most strongly enriched pathway terms for all 818 P-proteins assigned to CMGC, CAMK and AGC families. (H, I) Venn diagrams indicating the overlap of differentially regulated P-peptides from HCoV-229E-infected or PERKi-treated infected Huh7 cells (as shown in Fig. 5) that are also present in A549-ACE2 cells, with the 906 P-sites regulated by CMGC kinases in SARS-CoV-2-infected cells as shown in (E). (J) Overrepresentation analysis was performed for 131 gene IDs derived from 147 PERKi-sensitive P-sites shared between HCoV-229E and SARS-CoV-2-infected cells as shown in (I). The table shows the top10 enriched pathway terms sorted by p-value. (K) Heatmap showing the fold change of regulation by SARS-CoV-2 or SARS-CoV across all time points of the conserved set of 147 PERKi-sensitive P-sites. The table shows zoomed in views of the top10 up- or downregulated P-sites. Phosphorylated residues are marked in red. Yellow colors highlight the inducible phosphorylation of Ser63 of c-Jun. Ratio values are sorted according to the 24 h time point. (L) Pie chart illustrating the distribution of rank1 kinase families and individual kinases that phosphorylate 147 P-sites in SARS-CoV-2 or SARS-CoV-infected A549-ACE2 cells.

Out of 16,433 identified P-sites, 83% mapped to the human kinome motif atlas, and nearly half overlapped with sites detected in Huh7 cells, highlighting both conserved and cell-type-specific phosphorylation patterns **(Fig. 7C)**. Comparative motif enrichment of the 5,377 common P-sites revealed 94–135 kinases altered across time points in SARS-CoV and SARS-CoV-2 infected cells, with AGC and CAMK kinases dominating constitutive activity, while CMGC kinases progressively expanded to comprise ~45% of the regulated kinome by 36 h **(Fig. 7D)**. CMGC kinases were threefold more frequently ranked as primary regulators than CAMK or AGC members, accounting for ~900 dynamic P-sites in both infection models **(Fig. 7E)**.

Upon SARS-CoV-2 infection, these three kinase families collectively phosphorylated 1,262 unique sites, corresponding to 818 proteins, of which 125 (15%) overlapped and were associated with the same top overrepresented pathway terms as the HCoV-229E-regulated phospho-proteins shown in **Fig. 5** and in **Supplementary Fig. 6 (Fig. 7F-G)**.

Integration of SARS-CoV-2 and HCoV-229E phospho-proteomes identified 142–147 P-sites overlapping with CoV-regulated, PERKi-sensitive CMGC substrates in Huh7 cells **(Fig. 7H-I)**. The PERK-dependent subset mapped to pathways governing viral transcription, cell cycle control, mRNA processing, chromatin organization, and Rho GTPase signaling-paralleling, patterns observed in HCoV-229E infection and highlighting a conserved PERK–CMGC signaling axis **(Fig. 7J)**.

Phosphorylation of the canonical JNK substrate c-Jun at serine 63 exemplified this pattern: it was among the most strongly upregulated sites in both SARS-CoV and SARS-CoV-2 infected cells. More broadly, one-third of CMGC substrates were upregulated while two-thirds were suppressed, mirroring the remodeling observed under PERK inhibition in HCoV-229E infection **(Fig. 6K)**. In total, 44 CMGC family members were ranked as the top kinases phosphorylating one of the 147 identified CMGC substrates **(Fig. 6L)**. Notably, nearly half of these 147 substrates were attributed to multiple CDK or JNK isoforms as rank 1 kinases, for example, the motif surrounding serine 63 of c-Jun was assigned to JNK1 **(Fig. 6K-L)**. Network views emphasized the connections between related protein kinase subgroups and their substrates **(Supplementary Fig. 13)**.

In conclusion, these meta-analyses reveal a conserved coronavirus kinome signature dominated by CMGC kinases, including a PERKi-dependent subset. Rather than relying on a single kinase system, coronaviruses appear to exploit a large coordinated network of proline-directed kinases to orchestrate host cell remodeling and sustain viral replication.

## Discussion

In this work, we present a comprehensive kinome-scale dissection of kinase–substrate networks and their pharmacological modulation during RNA virus infection. Our results uncover that inhibition of the ER stress sensor PERK orchestrates profound, multilayered reprogramming of host cell regulation. PERK blockade not only reshapes the translational landscape but also remodels a coordinated network of proline-directed kinases and selectively alters functionally coherent modules within the CMGC kinase family. Integrating chemical and genetic perturbations, we reveal the intrinsic polypharmacology of PERK inhibitors and establish a conceptual framework for leveraging kinase network rewiring to drive an antiviral state, offering broad implications for therapeutic targeting of virus–host interactions.

Exploratory proteomic profiling of post-translational modifications (PTMs) uncovered extensive remodeling of both HCoV-229E and host proteins. Serine and threonine phosphorylation dominated the landscape; however, hundreds of lysine residues were also modified by ubiquitination or acetylation in response to coronavirus infection. Distinct linkage-specific ubiquitination patterns, including K48- and K63-linked chains, likely reflect cellular defense strategies, either by promoting ER-associated proteasomal degradation of viral or misfolded proteins, or by activating innate immune signaling (Mello-Vieira *et al*, 2023). Acetylation events may arise from metabolic reprogramming and, in turn, modulate the epigenome, for example through histone acetylation during the pronounced transcriptional activation previously observed in coronavirus-infected cells (Poppe *et al*., 2017). Notably, 19% of all identified proteins carried two distinct PTMs, and 3% harbored three, underscoring the prevalence of combinatorial regulation **(Supplementary Fig. 1B)**. While our study prioritized phosphorylation to define kinome-driven mechanisms, these findings highlight the importance of dissecting PTM crosstalk in future investigations.

We identified multiple post-translational modifications (PTMs) in HCoV-229E proteins, with ubiquitination and phosphorylation of the nucleocapsid (N) protein being the most prevalent. The multi-domain, multifunctional N protein is the most abundant coronavirus protein and plays central roles in binding, condensing, and encapsidating viral genomic RNA, as well as modulating immune evasion through sequestration of host factors (McBride *et al*, 2014). Structurally, the N protein comprises two folded domains: an N-terminal RNA-binding domain (N2) and a C-terminal dimerization / oligomerization domain (N4), flanked by three intrinsically disordered regions (N1, N3, and N5) (McBride *et al*., 2014). Hyperphosphorylation of N proteins from transmissible gastroenteritis virus (TGEV), mouse hepatitis virus (MHV), infectious bronchitis virus (IBV), and SARS-CoV has been documented, though its functional significance remained poorly understood (Calvo *et al*, 2005; Chen *et al*, 2005; Lin *et al*, 2007; White & Hogue, 2006).

The central N3 domain contains a serine–arginine (SR)–rich region and two leucine- or lysine-rich helices separated by a glutamine-rich strand. In SARS-CoV-2, this domain contacts nonstructural protein 3 (nsp3) at the replication organelle (RO) exit channel, positioning N protein adjacent to nascent viral RNA (Bessa *et al*, 2022; Wolff *et al*, 2020). In vitro studies indicate that multisite phosphorylation within the SR region disrupts RNA binding and N2:N4 interactions, functioning as a conformational switch that regulates genomic RNA packaging and unpackaging (Botova *et al*, 2024). Here, we mapped multiple phosphorylation sites within the SR region of HCoV-229E N protein, including S149, corresponding to S184 of SARS-CoV-2, which is the most strongly conserved phosphorylated residue, present in > 80% of 82 examined coronavirus strains (Yaron *et al*, 2022). In contrast, S145 is conserved only in HCoV-OC43 and HCoV-HKU1, but not in MERS-CoV, SARS-CoV, or SARS-CoV-2 (Yaron *et al*., 2022). Using mutant viruses and phosphorylation-specific antibodies, we provide orthogonal evidence for the functional importance of endogenous N protein phosphorylation at SS145/149 in supporting coronavirus replication.

Additionally, we identified C-terminal phosphorylation sites (SS364/S367) that are conserved in HCoV-229E and close relatives from the two species *Alphacoronavirus chicagoense* and *Alphacoronavirus amsterdamense* **(Supplementary Fig. 1C)**. While these sites are dispensable for replication, they appear to suppress ATF3 expression, potentially representing a viral strategy to attenuate host responses in this group of alphaviruses.

Motif analysis revealed that phosphorylation sites within the N protein represent consensus substrates for multiple kinases, including serine–arginine protein kinases (SRPK), glycogen synthase kinase 3 (GSK-3), and casein kinase 1 (CK1), which together constitute a hierarchical kinase cascade. This cascade, defined by in vitro assays, orchestrates clustered phosphorylation within the SARS-CoV-2 N protein SR region (Yaron *et al*., 2022). Strikingly, HCoV-229E N protein lacks both a CK1 priming phosphorylation site and the GSK-3β docking motif within its SR domain. This absence suggests a mechanistic divergence in alphacoronaviruses, implying that alternative kinases or phosphorylation cascades may govern SR domain regulation during infection, thereby contributing to species-specific modulation of viral replication and host interaction (Yun *et al*, 2022).

A kinome-centered transcriptomic analysis revealed that Huh7 cells express over 75% of the annotated human kinome at the mRNA level. Notably, mutation of the N protein exerted minimal impact on the cellular transcriptome and mRNA kinome, suggesting that N protein phosphorylation–mediated modulation of the host response is most likely to manifest at the translational level. This analysis further identified PERK as the sole endoplasmic reticulum (ER) stress / unfolded protein response (UPR) kinase transcriptionally upregulated in response to coronavirus infection.

PERK activation is canonically triggered by disturbances in ER protein folding capacity, alterations in membrane lipid bilayer composition, and changes in intracellular calcium levels, such as those induced by thapsigargin, conditions likely recapitulated during coronavirus-induced formation of replication organelles (Mazzolini & Touriol, 2025; Volmer *et al*, 2013). However, the precise role of PERK during coronavirus infection remains unresolved. Notably, we observed no HCoV-229E–mediated autophosphorylation of PERK at T980, a hallmark of canonical activation observed following ectopic expression and oligomerization of wild type PERK, but absent in catalytically inactive mutants (Carrara *et al*, 2015) **(Supplementary Fig. 14)**. This suggests that coronaviruses may activate PERK via a non-canonical mechanism, the nature of which remains to be defined.

Medicinal chemistry has produced thousands of PKi as chemical probes to interrogate the kinome (Reinecke *et al*, 2024). Among these, two PERK inhibitors, GSK2606414 and GSK2656157, have been extensively characterized. Initial profiling against panels of ~300 kinases, including the three additional integrated stress response kinases, demonstrated that 15–20 kinases were inhibited by > 80% at 10 µM, with IC_50_ values differing by several hundredfold, reflecting the high in vitro selectivity of these compounds (Axten *et al*., 2012; Axten *et al*., 2013). Nevertheless, their specificity has been questioned following reports of effective RIPK1 inhibition (Rojas-Rivera *et al*, 2017). In our system, PERK inhibitors potently suppressed HCoV-229E N protein phosphorylation at SS364/367 and virus-induced ATF3 expression, with low IC_50_ values of ~1–2 µM in intact cells, values congruent with those required for PERK inhibition. These observations prompted a detailed investigation of the global effects of PERK inhibition during coronavirus infection.

Coronaviruses impose a robust translational shutdown primarily through nsp1-mediated blockade of capped host mRNA entry into the ribosome, while allowing viral mRNA translation to proceed (Schubert *et al*, 2023). In parallel, infection activates the integrated stress response via Ser51 phosphorylation of eIF2α. Consistent with this, we observed a profound global reduction in translation rates for over 1,000 proteins spanning stress response, transcriptional regulation, and metabolic pathways. In non-infected cells, PERK inhibition reduced protein biosynthesis, in line with recent evidence that PERK inhibitors modulate the ATP affinity of the ISR kinase GCN2, paradoxically eliciting partial ISR activation under tunicamycin-induced ER stress (Szaruga *et al*, 2023). Strikingly, in coronavirus-infected cells, both pharmacological inhibition and genetic ablation of PERK significantly alleviated HCoV-229E–induced translational shutdown, indicating an on-target rescue effect that contributes to the antiviral state. Notably, P-eIF2α levels were not fully abrogated upon PERK inhibition, implying alternative mechanisms of ISR activation and the existence of alternative PERK-dependent pathways that are specifically engaged during coronavirus infection. Indeed, PERK has been reported to target additional substrates, including NRF2, FOXO transcription factors, and the lipid diacylglycerol, highlighting its broader regulatory role beyond canonical ER stress signaling (Bobrovnikova-Marjon *et al*, 2012; Cullinan & Diehl, 2004; Perea *et al*, 2023; Zhang *et al*, 2013).

Likely reflecting coronavirus-induced translational shutdown, our data revealed a striking uncoupling between mRNA expression and the proteome in infected cells. Notably, approximately two dozen protein kinases were downregulated in response to PERK inhibition, raising important questions about how these changes manifest within the phosphorylation landscape. Consistent with reports from SARS-CoV-2 studies across diverse cell types, we observed time-dependent remodeling of hundreds of phospho-proteins during HCoV-229E infection. Prior phospho-proteomic investigations have leveraged phosphorylation site data to infer limited sets of coronavirus-regulated kinases for targeted antiviral testing using small-molecule inhibitors (Bouhaddou *et al*, 2020; Fritch *et al*, 2023; Hekman *et al*, 2020; Higgins *et al*., 2023; Kohli *et al*, 2022; Stukalov *et al*., 2021; Wu *et al*, 2023). However, these studies did not examine PERK inhibitors, and to our knowledge, no previous work has systematically interrogated the global phenotypic consequences of a single kinase inhibitor at kinome-wide scale, as we have achieved here.

Integration of translational, proteomic, and phosphoproteomic datasets revealed profound molecular reprogramming at the level of the top 20 most significantly enriched Gene Ontology terms. This analysis identified eight coronavirus-modulated pathways that were consistently altered in both the translatome and phosphoproteome, encompassing RNA metabolism, RNA localization, ribosome biogenesis, chromatin remodeling, DNA metabolism, cell cycle regulation, growth factor signaling, and Rho GTPase signaling. In contrast, proteome-level changes were dominated by processes related to metabolism, mitochondrial function, and intracellular transport and trafficking **(Supplementary Fig. 16)**. These findings align with previous proteomic and phospho-proteomic studies of SARS-CoV-2 as shown by our bioinformatics analyses. Crucially, PERK inhibition broadly suppressed or remodeled gene sets across all of these major functional categories, impacting numerous sub-processes detailed throughout this study. Among these, suppression of Rho GTPase signaling emerged as a particularly prominent and consistent effect of PERK inhibition.

Rho GTPases constitute a family of twenty small GTP-binding proteins that function as molecular switches to orchestrate cytoskeletal dynamics, cell polarity, migration, and cell cycle progression (Coleman *et al*, 2004; Heasman & Ridley, 2008). Beyond their roles in cytoskeletal regulation, Rho GTPases integrate actin dynamics with the spatiotemporal control of membrane and vesicle trafficking, including processes between the Golgi apparatus and endoplasmic reticulum during exocytosis (Olayioye *et al*, 2019; Ridley, 2006). In their active GTP-bound state, Rho GTPases engage a diverse set of downstream effectors, notably including six p21-activated kinases (PAKs 1–6), which are multifunctional STE family kinases implicated in oncogenic signaling (Kumar *et al*, 2017; Sankaran *et al*, 2023). Although PAKs are not represented among the 303 Ser/Thr kinases of the human motif atlas, limiting direct substrate discovery, phosphorylated PAK2 was detected in our phospho-proteomic datasets. This finding prompted a targeted investigation of the Rho GTPase pathway using two ATP-competitive inhibitors selective for group I PAKs (PAK1–3), FRAX486 and FRAX597, which were originally developed as therapeutic agents for neurofibromatosis, certain inherited cancers, and fragile X syndrome (Dolan *et al*, 2013; Licciulli *et al*, 2013).

Group I PAKs function as homodimers that are activated through conformational rearrangements and multi-site phosphorylation within their N-terminal regulatory and C-terminal kinase domains. Binding of GTP-loaded Rho GTPases to the p21-binding domain (PBD) relieves autoinhibition, which arises when the auto-inhibitory domain (AID) of one PAK molecule interacts with the PBD of another (Kumar *et al*., 2017). Autophosphorylation at Ser144 in PAK1 or Ser141 in PAK2 is a hallmark of active PAK dimers (Gatti *et al*, 1999). In Huh7 cells, both residues were constitutively phosphorylated, consistent with basal PAK1/2 activation. Upon HCoV-229E infection, treatment with either of two PAK inhibitors markedly suppressed PAK autophosphorylation, virus-induced ATF3 expression, N protein phosphorylation, and viral replication at low micromolar concentrations (1–2 µM), establishing a central role for Rho GTPase signaling in CoV replication and host response modulation.

Supporting these findings, recent studies demonstrated that FRAX486 inhibits SARS-CoV-2 replication in diverse cell lines, primary nasal epithelial organoid cultures, and an in vivo hamster model (Liu *et al*, 2023; Wu *et al*., 2023). These studies linked PAK1 (and group II member PAK4) to microvillar remodeling and viral egress, as well as to cytoskeletal rearrangements and autophagy-dependent endocytosis (Liu *et al*., 2023; Wu *et al*., 2023). Notably, other work reported FRAX486’s antiviral activity to be cell-type dependent, effective in Huh7.5 cells but not in Calu-3 lung cells, and associated with cytotoxicity at concentrations exceeding 10 µM (Dittmar *et al*, 2021; Liu *et al*., 2023). In contrast, FRAX597 exhibited antiviral activity without prior characterization and appears to possess a more favorable cytotoxicity profile. Collectively, these data highlight the potential of PAK inhibitors, originally developed as anticancer agents, as promising antiviral strategies against CoVs and possibly 12 other viruses, warranting further investigation into their mechanisms of action and therapeutic windows (Aiqui-Reboul-Paviet *et al*, 2024; Casado *et al*, 2025; Van den Broeke *et al*, 2010).

Our findings further reveal that CoV-induced PERK activation is attenuated by PAK inhibitors, uncovering a previously unrecognized, indirect crosstalk between PERK signaling and Rho GTPase pathways during CoV infection. PERK activation is known to be modulated by actin cytoskeleton remodeling and plays a critical role in coordinating ER–plasma membrane apposition through cytoskeletal regulation (van Vliet *et al*, 2017). Recent evidence highlights that ER stress orchestrates ER architecture in concert with cell morphology via PERK-dependent mechanisms (Sanchez-Alvarez *et al*, 2025). These observations raise the intriguing possibility that perturbation of Rho GTPase signaling, thereby linking cytoskeletal dynamics to ER stress and PERK function, could modulate the viral life cycle. Targeted disruption of these pathways using emerging allosteric GTPase inhibitors represents a promising avenue for future investigation into their antiviral potential (Morstein *et al*, 2024).

By integrating our comprehensive analysis of PERK inhibition with motif-based interrogation of phospho-proteomes, we delineate the regulatory phosphorylation landscape of CoV-infected cells from a kinome-centric perspective. This analysis leverages a compendium of ~89,000 experimentally characterized human phosphorylation sites and consensus motifs for 303 Ser/Thr kinases.

A principal finding of our study is that CoV infection drives dynamic activation of the CMGC kinase family in its entirety. This family comprises 63 kinases, including cyclin-dependent kinases (CDKs), glycogen synthase kinases (GSKs), mitogen-activated protein kinases (MAPKs), and CDK-like kinases (CDKLs), as well as smaller subgroups such as HIPKs, DYRKs, and SRPKs (Chowdhury *et al*, 2023).

Of particular significance are CDKs, a large and multifunctional kinase family of 20 members, which coordinate cell cycle progression, regulate RNA polymerase II-mediated transcriptional initiation, pause release, and elongation, and are intensively investigated for their oncogenic roles (Pluta *et al*, 2023). Notably, infection with SARS-CoV and SARS-CoV-2 has been associated with downregulation of specific cyclins, the regulatory subunits of CDKs, thereby inducing cell cycle arrest (Gupta & Mlcochova, 2022; Surjit *et al*, 2006). Consistently, dinaciclib, a multi-CDK inhibitor targeting CDK1, 2, 5, and 9, has been reported to suppress SARS-CoV-2 replication across independent studies, albeit with pronounced cytotoxicity (Bouhaddou *et al*., 2020; Fritch *et al*., 2023). These observations underscore a critical gap in understanding the roles of CDKs within key pathways perturbed during viral replication, including cell cycle regulation and nuclear processes such as DNA metabolism and chromatin remodeling, warranting further mechanistic investigation (Pellarin *et al*, 2025).

Our findings corroborate robust CoV-mediated activation of the JNK and p38 MAP kinase signaling pathways, consistent with prior observations in infectious bronchitis virus (IBV), SARS-CoV, and SARS-CoV-2 infections, and extend these insights by linking both pathways to the coordinated activation of the broader CMGC kinase network (Fritch *et al*., 2023; Liao *et al*, 2011; Mizutani *et al*., 2004, 2005). Recent work from our group demonstrated that pharmacological inhibition of JNK markedly suppresses HCoV-229E replication and N protein phosphorylation (Bruggemann *et al*, 2025). Notably, c-Jun, a highly specific JNK substrate and a member of the AP-1 family of basic leucine zipper transcription factors, functions as a pivotal regulator of chromatin-dependent processes relevant to cancer, inflammation, and host-pathogen interactions (Bejjani *et al*, 2019). Activation of the JNK pathway by stressors, including small Rho GTPases such as Cdc42 and Rac1, suggests an uncharacterized mechanistic link to the CoV-induced Rho GTPase signaling axis (Minden *et al*, 1995). c-Jun is a multi-site phosphorylated and strongly regulated protein (Derijard *et al*, 1994; Morton *et al*, 2003). Phosphorylation of Ser63 within c-Jun’s N-terminal transactivation domain emerges as one of the most prominent and consistently regulated phosphorylation events in CoV-infected cells, highlighting its potential centrality in the viral host response. Elucidating the transcriptional programs governed by the JNK–P-c-Jun axis, in conjunction with AP-1 partner proteins such as ATF3, will be critical to resolving their integrated roles in shaping the transcriptional landscape of CoV infection.

Our study uncovers a conserved core module of ~150 functionally interconnected substrates within the CMGC kinase network that is consistently activated across HCoV-229E, SARS-CoV, and SARS-CoV-2 infections and suppressed by PERK inhibition. This defines a fundamental regulatory phosphorylation landscape central to the CoV host response. While the absence of direct PERK phosphorylation sites and the persistence of CMGC kinase regulation in PERK-deficient cells indicate that many PERKi effects are off-target, this finding reframes PERK inhibition as a powerful strategic lens to interrogate, and pharmacologically modulate, the CMGC kinase network. The phosphorylation sites revealed here, integrated with translational and kinome dynamics, constitute a rich atlas for future targeted functional dissection of RNA virus infection biology, including through genome-scale CRISPR base editing (Kennedy *et al*, 2024). Beyond their virological relevance, these findings establish a conceptual framework for deciphering the intricate interplay between translatomes, proteomes, and phospho-proteomes that governs the phenotypic outcomes of small-molecule kinase inhibition, with broad implications for antiviral drug development.

## Limitations

This study was conducted in a single cell type and focused on one human coronavirus to ensure experimental consistency and enable integration across multiple molecular layers. While our label-free LC-MS/MS approach provided deep coverage, it did not capture the full proteome or phospho-proteome due to inherent detection limits. Nonetheless, we identified over 12,000 phosphorylation sites, representing substantial overlap with the recently reported human consensus phospho-proteome (Johnson *et al*., 2023). Despite affinity enrichment strategies, many low-abundance regulatory factors, including phosphorylated PERK, eIF2α, and JNK, remain underrepresented, reflecting a known limitation in phospho-proteomics.

Our analyses were performed on whole-cell extracts, precluding insight into subcellular kinome regulation, for example at the endoplasmic reticulum, which may explain the absence of certain specifically regulated PERK substrates. Kinase–substrate assignments based on the human motif atlas improve upon previous purely in silico approaches; however, restricting analyses to rank 1 kinases inevitably underrepresents the inherent redundancy of kinase substrate recognition. Generation of regulated nascent and phosphorylated protein lists required user-defined filtering criteria based on controls, enrichment levels, and statistical thresholds. As with all bioinformatics workflows, adjusting these criteria influences the resulting datasets. Likewise, functional annotations depend on underlying databases and default parameters in tools such as Metascape and STRING.

Importantly, we provide the complete raw dataset and source data to enable reproducibility, transparency, and independent reanalysis. This resource will allow the field to refine filtering strategies, expand kinase–substrate relationships, and generate alternative interpretations tailored to specific biological questions.

## Materials and Methods

### Cell lines and viruses

Huh7 human hepatoma cells used in this study were obtained from the Japanese Collection of Research Bioresources (JCRB) cell bank (Nakabayashi *et al*, 1982). The cells were maintained in Dulbecco’s modified Eagle’s medium (DMEM) with the following contents: 3.7 g / 1 NaHCO_3_; PAN Biotech Cat No P04-03550), 10% filtrated bovine serum (FBS Good Forte; PAN Biotech, Cat No. P40-47500), 2 mM L-glutamine (Gibco, 21935-028), 100 U/ml penicillin and 100 μg/ml streptomycin. Standard trypsinization and passaging protocols were followed.

CV-1 (from African green monkey kidney) and Hela-D980R cells (Weissman & Stanbridge, 1980) were used to generate and plaque-purify recombinant vaccinia viruses containing genome-length HCoV-229E cDNA as described previously (Thiel *et al*, 2001). High-titer virus stocks of recombinant vaccinia viruses containing HCoV-229E cDNA were prepared using CV-1 or BHK-21 cells (kidney of a Syrian hamster) (Thiel *et al*., 2001). The cells were cultured in Dulbecco’s Modified Eagle’s Medium (DMEM) supplemented with 10% [v/v] fetal bovine serum (FBS), 100 U/mL penicillin and 100 µg/mL streptomycin.

The genome sequence of the human coronavirus (HCoV)-229E used in this study is available from NCBI, accession number of AF304460.1, and reference sequence of NC_002645.1. Huh7 cells were used to propagate wild type (wt) HCoV-229E and HCoV-229E mutants. Virus titers were determined on Huh7 cells using TCID_50_/ml (for stocks) or by plaque assay (see below).

### Plasmid vectors, CRISPR-CAS9-mediated genome editing and transfections

To generate cell lines lacking either PERK or IRE1α, Huh7 cells were stably transfected with CRISPR-CAS-9 plasmids carrying sgRNAs targeting genes of interest (*EIF2AK3*, *ERN1*) with the following sequences: sg1_EIF2AK3:se, 5’-CACCGTGGAGCGCGCCATCAGCCCG-3’; as 5’-AAACCGGGCTGATGGCGCGCTCCAC-3’; sg1_ERN1: se, 5’-CACCGTCACCGCCTCGCTGTCGTCG; as 5’-AAACCGACGACAGCGAGGCGGTGAC-3’. Briefly, the sgRNAs (designed online using https://crispr.mit.edu from the Zhang lab, now discontinued, see https://zlab.squarespace.com/guide-design-resources) were synthesized by Eurofins Genomics as sense and antisense strands. The obtained oligonucleotides were re-suspended in appropriate volumes of water to reach a final concentration of 100 μM as indicated by the synthesis report and then annealed using the following amounts and components: 1 μl of sgRNA sense and antisense oligonucleotides (100 μM), 5μl of T4 PNK buffer A (10x) and 43 μl ddH_2_O using the following temperatures profile: 95 °C for 4 minutes, 70 °C for 10 minutes, 37 °C for 15 minutes and 4 °C as pause temperature. The annealed oligonucleotides were then phosphorylated using the following components: 2 μl Annealed oligonucleotides, 1 μl T4 PNK (polynucleotide kinase) buffer A (10x), 1 μl 1 mM ATP (10 mM), 1 μl T4 PNK and 5 μl ddH_2_O. The temperatures profile were as follow: 37 °C for 30 minutes, 70 °C for 10 minutes (heat inactivation of T4 PNK enzyme) and 4 °C as pause temperature. The now double stranded oligonucleotides were then cloned into compatible pSpCas9(BB)-2A-Puro (pX459) vector using appropriate restriction enzymes (following the manufacturer-provided protocols). The double digest reaction and ligation were carried out together as follow: 2 μl diluted oligonucleotides, 100 ng vector, 2 μl Fast digest buffer (10x), 1 μl Fast digest Bbs1, 1 μl ATP (10 mM), 1 μl T4 ligase and ddH_2_O adjusted to 20 μl with the following temperatures profile for 6 cycles: 37 °C for 30 minutes and 21°C for 10 minutes. A final step included the use of Plasmid-Safe exonuclease (Lucigen #E3101K) to digest residual, linearized DNA as follow: 10 μl The ligation reaction from the previous step, 1.5 μl Plasmid-Safe buffer (10X), 1.5 μl ATP (10 mM) and 1 μl Plasmid-Safe exonuclease (10 U/μl) using the following temperature settings: 37 °C for 30 minutes and 70 °C for 30 minutes. The ligated plasmid DNA was then transformed into chemically competent bacterial strains (E. coli TOP10, XL1-Blue from Thermo Fisher Scientific or E. coli bacteria from NEB # C3040H). Plasmids-producing bacteria were then allowed to grow overnight in LB medium at 37 °C (continuous shaking) and harvested for plasmid isolation. The DNA plasmids were then extracted using the NucleoSpin® plasmid kit (Machery & Nagel) following the manufacturer’s protocol. The success of the cloning process was confirmed using Sanger sequencing. All thermal reactions were carried out on Thermocycler T-Professional.

The successfully cloned plasmids were transfected into parental Huh7 cells using calcium phosphate (CaCl_2_) method. Briefly: 9x10^5^ Huh7 cells were incubated in a solution containing 1500 μl H_2_O, 1350 μl 2x HEBS, 30 μg plasmid and 189 μl 2 M CaCl_2_ for 5 hours followed by a glycerol shock (3 ml of DMEM with 10% glycerol) for 3 minutes. Thereafter, the cells were returned to normal growth medium and subsequently selected for the successful integration of the plasmids using puromycin (10 µg/ml). All cell lines (parental or engineered) used in this study were tested regularly for mycoplasma contamination using PCR-Test kit (#A3744, AppliChem).

Published plasmid vectors encoding murine wild type (#21814, Addgene) or catalytically inactive Myc-tagged PERK (#21815, Addgene) and were transfected using calcium phosphate as described above (Harding *et al*., 1999).

### Generation of mutant viruses

The two HCoV-229E mutants used in this study, HCoV-229E_N-SS145/149AA and HCoV-229E_N-SS364/367AA, were generated using the recombinant vaccinia virus vHCoV-inf-1 containing a full-length cDNA of the HCoV-229E genome (NCBI reference sequence NC_002645) as described previously (Thiel *et al*., 2001).

First, vRec-3’GPT, a derivative of vHCoV-inf-1, was generated by recombination using pBS-3′GPT plasmid DNA. This pBluescript II-derived plasmid was constructed to contain the E. coli gpt gene flanked by (i) a 503-bp fragment representing the cDNA sequence of nts 24,473–24,976 of the HCoV-229E genome and (ii) a 500-bp fragment representing the vaccinia DNA sequence located downstream of the HCoV-229E cDNA insert in vHCoV-inf-1. The gpt-positive vHCoV-inf-1 derivative vRec-3′GPT was selected from CV-1 cells that had been infected with vHCoV-inf-1 and transfected with pBS-3′GPT plasmid DNA.

For the second recombination step, D980R cells were infected with vRec-3′GPT and transfected with the appropriate pBS-3’_mut plasmid DNA containing the desired mutations in the N gene. The pBS-3′_mut plasmid constructs used to produce the vHCoV-inf-1 derivatives with the desired N gene mutations contained a cDNA of HCoV-229E nts 24,473-27,317 with the desired mutations (introduced by PCR-based mutagenesis) and the 500-bp vaccinia virus sequence downstream of the HCoV-229E polyA sequence present in vHCoV-inf-1. Recombined gpt-negative vHCoV-inf-1 derivatives containing the desired mutations in the cDNA sequence of the N gene were isolated using appropriate selection conditions (Hertzig et al., 2004). Southern blot and sequence analyses were used to confirm the correct assembly of the HCoV-229E cDNA in the vHCoV-inf-1 derivatives (Thiel et al., 2001).

Next, 5’-capped genome-length HCoV-229E (wt and mutants) RNAs were prepared by in vitro transcription using T7 RNA polymerase (RiboMAX Large Scale RNA Production System, Promega) using purified vaccinia virus genomic DNA isolated from the respective vHCoV-inf-1 derivatives. 1.25 µg of in vitro-transcribed genome-length RNAs and 0.75 µg of in vitro-transcribed mRNA encoding the HCoV-229E N protein (Schelle et al., 2005, Almazan et al., 2006) were used to co-transfect 1x10^6^ Huh7 cells using the TransIT® mRNA transfection kit according to the manufacturer’s instructions (Mirus Bio LLC). At 72 h post-transfection (p.t.), cell culture supernatants were collected and virus titers were determined by plaque assay for three independent transfection experiments. Additionally, total RNA was isolated from the cells and used for genome sequence analyses.

The following oligonucleotides were used for site-directed mutagenesis of the pBS-3’ plasmid DNA: S145+149A: se, 5’-AGAACCTGACgCCCGTGCTCCTgCCCGGTCTCAGTCGAGGTCG-3’; as, 5’-GACTGAGACCGGGcAGGAGCACGGGcGTCAGGTTCTTCAACAACAGTAAC-3’; S364+367A: se, 5’-ATTCAACCCAgCTCAAACTgCACCTGCAACTGCTGAACCAGTGCGTGA-3’; as, 5’-CAGTTGCAGGTGcAGTTTGAGcTGGGTTGAATTCTAGTGCACTAGGGTT-3’.

For further analysis and validation using Western blot (**Fig. 1C**), 0.5-l x10^6^ Huh7 cells were either mock infected or infected with HCoV-229E wt or one of the HCoV-229E N gene mutants at 33 °C using a multiplicity of infection (MOI) of 1 or 0.1 plaque-forming units (pfu)/cell, respectively. Infection experiments were done using virus stocks obtained from two independent virus rescue (i.e, RNA transfection) experiments. For each of those, two independent biological replicates were performed, resulting in four independent samples. At one hour p.i., the virus inoculum was removed and replaced with fresh medium. After incubation for 24 hours at 33 °C, the culture medium was removed and the cells were either lysed as described below or total RNA was isolated using the IntronBio total RNA extraction kit according to the manufacturer’s instructions and used for microarray experiments.

### Virus infections and assessments of antiviral activity

To understand the role of PERK or indicated downstream effectors in viral replication, the Huh7 (parental or mutants) cells were infected with HCoV-229E (wild type or mutant) at the indicated multiplicities of infection (MOI). The virus-infected cells were incubated at 33°C in the presence or absence of indicated inhibitors, chemicals or corresponding amounts of DMSO as a solvent control. At the end of indicated incubation times (12 or 24 h p.i) cells were harvested for downstream analysis and supernatants were collected and stored at −80 °C for viral titer determinations using plaque assay.

The plaque assay was performed as follows: Briefly, a confluent monolayers of Huh7 cells were incubated with serial dilutions of HCoV-229E-containing supernatants, derived from indicated experimental conditions. The virus was diluted 10^1^ to 10^7^ and incubated at 33 °C. After 1 h, the virus inoculum was replaced with fresh medium (MEM, Gibco) containing 1.25% Avicel® (FMC Biopolymer, #RC591), 100 U/ml penicillin, 100 μg/ml streptomycin, and 10% FBS. At 72 hpi. cell culture supernatants were removed and, the cell layer was washed with PBS and stained with 0.15% (w/v) crystal violet (diluted in 20% ethanol) and plaques were counted. All experiments involving HCoV-229E and recombinant HCoV-229E N gene mutants were performed in biosafety level 2 (BSL-2) containment laboratories approved for such use by the German authorities.

## Materials

Thapsigargin (Cay10522-1), GSK2656157 (Cay17372) and GSK2606414 (Cay17376) were purchased from Cayman Chemicals. FRAX486 (TGM6840) and FRAX597 (TGM6014) were obtained from TargetMol and KIRA6 (532281) was purchased from Calbiochem through Merck-Millipore. All these chemicals were dissolved in DMSO as 10 mM or 5 mM (for FRAX597) stock solutions. Appropriate DMSO concentrations served as vehicle controls in indicated experiments. The following inhibitors were used: leupeptin hemisulfate (Carl Roth, #CN33.2), microcystin (Enzo Life Sciences, #ALX-350012-M001), pepstatin A (Applichem, #A2205), PMSF (SigmaAldrich, #P-7626). Pepstatin A, PMSF and microcystin were dissolved in ethanol and leupeptin in water. Other reagents were from Sigma-Aldrich or Thermo Fisher Scientific, Santa Cruz Biotechnology, Jackson ImmunoResearch, or InvivoGen and were of analytical grade or better.

Primary antibodies against the following proteins or peptides were used: anti β-actin (Santa Cruz, #sc-47778), anti PERK (Santa Cruz, #sc-377400), anti PERK (Abcam, #ab65142), anti P(T980)-PERK (Cell Signaling, #3179, 16F8), anti eIF2α (Cell Signaling #9722), anti P(S51)-eIF2α (Cell Signaling #9721), anti P(S724)-IRE1α (Novus Biologicals, #NB100-2323), anti IRE1α (Santa Cruz, #sc-390960), anti ATF3 (Santa Cruz, #sc-188), anti HCoV-229E N protein (Ingenasa, Batch 250609), anti-HCoV-229E P(S145/S149)N protein (Eurogentec, On-demand raised AB), anti HCoV-229E P(S364/S367)N protein (Eurogentec, On-demand raised AB), anti puromycin [3RH11] (Kerafast Inc., EQ0001), anti PAK2 (Cell Signaling, 2608), anti P(S144)-PAK1/P(S141)-PAK2 (Cell Signaling, 2606). anti c-Jun (Invitrogen; # MA5-15889), anti P(S63)-c-Jun (Cell Signaling; #9261), anti JNK(Cell Signaling, 9252), anti P(T183/Y185) JNK (Cell Signaling, 9251), anti p38 MAPK (Cell Signaling, 9212), anti P(T180/Y182)-P38 (Zymed, 36-8500), anti Myc (Roche, #11667149001, 9E10).

Purified rabbit polyclonal antibodies against HCoV-229E N protein phosphorylation sites (marked by asterisks) as identified by mass spectrometry were raised against P(SS145/147) (peptide LPNGVTVVEEPDS*RAPS*R) and P(SS364/367) (peptide EMQQHPLLNPSALEFNPS*QTS*PATAEPVRDEVSIETDIIDEVN) using a commercial immunization program by the company Eurogentec (4102 Seraing, Belgium).

First, synthetic peptides were generated for modified (peptide 1511422, h-C+EEPD-S(PO3H2)-RAP-S(PO3H2)-RSQ-nh2) and unmodified SS145/149 residues (peptide 1511423) or modified (peptide 1511424, h-C+EFNP-S(PO3H2)-QT-S(PO3H2)-PATA-nh2) and unmodified SS364/367 residues (peptide 1511425). All peptides were validated by MALDI-TOF mass spectrometry and each P-peptide plus proprietary adjuvants including keyhole limpet hemocyanin (KLH) were injected into two rabbits **(Supplementary Fig. 16A)**.

After 3 weeks, pre-immune sera and large bleeds from both rabbits were analyzed for the presence of high titer antibodies against the P-peptides or the carrier by specific ELISA using plates coated with P-peptides 1511422, 1511424 and carrier protein **(Supplementary Fig. 16B)**.

One week later, the final bleeds from one rabbit (1507 for P(SS145/19)-N) and 1510 for P(SS364/367)-N) were used to purify high titer P-specific antibodies from the sera: First, N protein-specific antibodies were adsorbed to P-peptide affinity columns, the flow through fraction F1 was discarded and the peak fraction P1 eluted. Next, P1 was further purified by non-phospho-peptide coupled affinity chromatography and the flow-through F2 collected, which contained the double purified highly specific antibody fractions recognizing either P(SS145/149) or P(SS364/367). Each purification step was validated by specific ELISA against P-peptide, carrier or unmodified peptide. Purity of the final IgG fraction was 87.2% for P(SS145/149) and 83.4% for P(SS364/367) antibodies and antibody titers for both P-peptide antibody preparations were > 1x10^5^ as summarized in **(Supplementary Fig. 16C)**.

The horseradish peroxidase (HRP)-coupled secondary antibodies used in this study were the following: Dako P0447; polyclonal goat anti-mouse immunoglobulins/HRP, Dako P0448; polyclonal goat anti-rabbit immunoglobulins/HRP.

### Cell lysis, in vivo puromycinylation, and Western blotting

Whole cell extracts from samples created for Western blot analysis were lysed in Triton X-based cell lysis buffer with the following components: 10 mM Tris, pH 7.05, 30 mM NaPPi, 50 mM NaCl, 1% Triton X-100, 2 mM Na_3_VO_4_, 50 mM NaF, 20 mM ß-glycerophosphate and freshly added 0.5 mM PMSF, 2.5 µg/ml leupeptin, 1.0 µg/ml pepstatin, 1 µM microcystin. Briefly: At the end of indicated incubation/treatment time; cells were scraped in ice-cold PBS and pelleted at 500 x g (5 min at 4°C). Cells were either snap frozen in liquid nitrogen (N2) or immediately lysed as follow: Appropriate volumes of the lysis buffer were added to the centrifuged cell pellets and incubated for 10-15 min on ice. The lysates were then centrifuged at 15000 x g for 15 min (4°C) to clear cellular debris. Bradford assay was then used to measure the concentrations of protein in each sample and to determine loading-on-gel volumes.

For all samples created for Western blotting, aliquots corresponding to 20-50 µg protein (per lane) were mixed with 4 x SDS sample buffer (ROTI^®^Load, Roth, #K929) and either stored at −80 °C or loaded immediately on SDS-PAGE. The loaded samples were subjected to SDS-PAGE with varying concentrations of acrylamide (8-12.5%) gels depending on the antibodies used after blotting. The Thermo Scientific PageRuler™ prestained protein ladder (#26616) was used as molecular weight marker.

Imunoblotting was carried out as follows: first; proteins were separated on SDS-PAGE and electrophoretically transferred to PVDF membranes (Roti-PVDF, #T830 Roth or Immobilon-P, IPVH00010). To confirm efficient protein transfer and equal loading, the membranes were stained with Ponceau S (Sigma, 0,1 % (w/v), dissolved in 5% acetic acid). Blocking of the membranes was done with 5% dried milk in Tris-HCl-buffered saline/0.05% Tween (TBST) for 1 h and then they were incubated for 12-24 h with primary antibodies. Thereafter, the membranes were washed in TBST and incubated for 1-2 h with compatible peroxidase-coupled secondary antibody. Proteins were detected using enhanced chemiluminescence (ECL) systems from Millipore or GE Healthcare. Images were acquired with the ChemiDoc Touch Imaging System (BioRad) and quantified using the software ImageLab (versions V_5.2.1 or V_6.0.1, Bio-Rad).

Nascent polypeptides were metabolically labeled in intact cells as reviewed in (Iwasaki & Ingolia, 2017). Huh7 cells were seeded in appropriate cell culture dishes (3-9 x 10^5^ cells) and treated as indicated. Half an hour before harvesting, 3 µM of puromycin (InvivoGen, #ant-pr-1) was added to the medium. The cells were then lysed and immunoblotting was carried out as described above. Membranes were initially stained with Coomassie brilliant blue (CBB) or Ponceau S as control for steady-state protein levels and then hybridized with the anti puromycin antibody to detect nascent polypeptides.

Total cell lysates used for the total proteome or phospho-proteomics mass spectrometry experiments were prepared as follows: Cells were scraped in ice-cold PBS and pelleted at 500 x g for 5 min at 4°C and stored in liquid N2. After thawing, cell pellets from all biological replicates were lysed together (at the same time) using urea-based lysis buffer composed of 20 mM HEPES, pH 8.0, 9.0 M urea (Thermo Scientific, Sequanal Grade, #t29700), 1 mM activated sodium orthovanadate, 2.5 mM sodium pyrophosphate, 1 mM ß-glycerophosphate, and water for volume adjustments. The samples were then sonicated using Bioruptor sonicater for 4 min with 30 sec on; 30 sec off, high power settings and centrifuged for 15 min (4°C) at 20000 x g. Supernatants were then transferred to a fresh tubes and protein concentrations were determined with the detergent compatible Bradford assay kit (Pierce™, #23246) using a 300-fold dilution.

### Click chemistry-based OPP labelling of nascent polypeptide chains

To understand the role of PERK in host response to HCoV-229E infection on the level of the nascent proteome, newly synthesized polypeptide chains were labelled with OPP (O-propargyl-puromycin) using a click chemistry-based approach based on (Forester *et al*., 2018).

Briefly, 3x10^6^ Huh-7 cells were seeded in appropriate dishes 24 hours before the beginning of the infection or the inhibitor treatment. Cells were then moved to 33 °C incubator for half an hour and subsequently HCoV-229E or GSK2656157 were added at the indicated MOI and concentrations. Cycloheximide (10 µg/ml), as a negative control for the global arrest of translational elongation, was added 3 hours before harvesting. 2 hours before the end of the 24 hours infection/treatment period (i.e. 2 hours before harvesting) the cells were treated with either 30 μM of OPP (Hycultec, #HY-15680-5) or 0.3% DMSO (as solvent control). Cells were then harvested as previously described and either stored at −80 °C or directly lysed. The cells were lysed with 1 ml of click chemistry lysis buffer (100 mM HEPES pH 7.5, 150 mM NaCl, 1% NP-40, 2 mM PMSF, 1X cOmplete Protease Inhibitor Cocktail (Roche; #11836170001) with incubation on ice for 15 minutes with intermittent vortexing followed by centrifugation at 16.000 x g (also for the following steps unless otherwise stated) for 30 minutes at 4°C.

The supernatants were transferred to new tubes and protein concentrations were then determined using PierceDetergent Compatible Bradford Assay Kit (Thermo Scientific, # 23246) with a dilution of 1:300 in PBS. At this point, 25 μg of the lysate was saved from each sample and used later for Western blot.

Subsequently, to remove endogenously biotinylated proteins, a pre-clearing step of the lysates was carried out as follows: For each sample, 60 μl of high capacity streptavidin agarose resin (Thermo Scientific # 20361) were equilibrated with 500 μl click-chemistry lysis buffer in 1.5 ml tubes. The beads-lysis buffer mixture was then centrifuged at 1000 x g for 2 min at 4°C. Supernatants were then discarded carefully and volumes of lysates corresponding to 1.8 mg proteins (per Bradford assay) were added to the beads. Varying amounts of click-chemistry lysis buffer were added to make a total volume of 1 ml in each sample followed by overnight 22 RPM rotation at 4 °C.

Samples were then centrifuged (1000 x g, 2 minutes, 4 °C), beads were discarded and the supernatants (the now pre-cleared lysates) were collected in preparation for the conjugation of biotin-azide (Caymanchemical, #13040) to OPP. The click reaction for the conjugation consisted of the following reagents: 0.625 mM biotin-azide, 6.25 mM tris(2-carboxyethyl)phosphine (TCEP, Sigma, #C4706), 0.625 mM tetraethylene pentamine (TBTA, Sigma, #678937), 6.25 mM CuSO4*5H_2_O and 20% SDS. The stock solutions of biotin-azide and TBTA were degassed through argon to enhance their stability. 160 μl of the click reaction was mixed with 1 ml of pre-cleared lysate (for each sample). The conjugation mixture was then allowed to rotate (22 RPM) at room temperature for 1.5 hour. Thereafter, the now conjugated lysate (“click-lysate” or OPP-biotin labeled lysate) was transferred to 5 ml Greiner tubes and mixed with 4 ml acetone (cooled down to −20 °C) to precipitate proteins. The mixture was then incubated overnight at −20 °C.

In the following day, the acetone-click-lysate mixture was centrifuged at 3500 x g for 5 minutes at 4 °C. The supernatants (containing Acetone) were discarded and the pellets were washed two times with 2 ml methanol cooled down to 4°C. Vortexing in-between washes and centrifugation at 3500 RPM for 5 minutes at 4°C were carried out. After the final washing step all remaining methanol was carefully removed by pipetting and drying the pellet for 5 minutes at room temperature.

The pellets were then re-suspended in 120 μl PBS with 1% SDS at room temperature followed by desalting and re-buffering of the solution using Zebra Spin Columns (Thermo Scientific, #89882 7k MWCO) according to the manufacturer’s guide. This step ensures an efficient removal of excess biotin-azide, preventing the streptavidin beads from saturating (see below). First, the columns were equilibrated twice with 300 μl of washing buffer consisting of PBS with 1% NP-40 & 0.1% SDS. In each step, the flow through was removed through one minute centrifugation at 1500 x g. The now equilibrated columns were placed onto new collection tubes and the re-suspended pellets were added. The samples were then centrifuged at 1500 x g for 2 minutes. The flow through is the click-eluate and was kept on ice.

The protein concentrations in the click-eluate were determined (Pierce Bradford assay) using dilutions of 1:400 in PBS. 25 μg of the click-eluate was collected from each sample for Western blot analysis. The streptavidin agarose beads pulldown was then carried out as follows: 60 μl of high capacity streptavidin agarose beads were equilibrated with 500 μl of 1% NP-40, 0.1% SDS in PBS buffer, followed by centrifugation at 2000 x g for one minute at 4°C. The supernatant was discarded and the beads were then incubated with volumes of click-eluates corresponding to 600 μg of proteins (per Bradford assay) and up to 700 μl of 1% NP-40, 0.1% SDS in PBS was added to each tube. The beads-click-eluates mixtures were then allowed to rotate (22 RPM) overnight at 4°C.

The following day, the mixtures were centrifuged at 1000 x g for 2 minute and supernatants were discarded. The beads were then washed 2 times with 500 μl of PBS with 1% NP-40, 0.1% SDS, 2 times with 500 μl of ice-cold PBS with 6 M Urea and 3 times with 500 μl of ice-cold PBS. Each washing step was carried out with 10 minutes rotation (22 RPM) at 4°C. Centrifugation at 1000 x g, 4°C for 2 minutes was done to remove the buffer from the previous wash step.

After the final washing step, the beads were incubated with mass spectrometer buffer (20 mM Tris-HCL pH 8.0, 2 mM CaCl_2_) and shipped on ice to the mass spectrometer facility (never frozen) for on-beads trypsin digestion.

For Western blot analysis, 50 μl ROTI®Load 1 (2x) was added and the samples were heated at 95°C for 20 minutes. Eluted samples were either stored at −20°C or loaded directly on SDS-PAGE.

### Cell viability assays

MTS assays were performed using the The CellTiter 96® AQueous One Solution Cell Proliferation Assay kit (Promega, #G3582) as described by the manufacturer protocol. Briefly: 1.2 x 10^4^ Huh7 cells were seeded in 96-well plates for 24 hours and thereafter treated with indicated chemicals, virus alone or virus plus chemical for 24 hours. Then, the medium was replaced by 100 µl complete cell culture medium including 20 µl CellTiter 96® AQueous one solution reagent and the cells were further incubated for 1 hour at 33 °C. Absorbance values were then measured at 490 nm. Background absorbance was corrected for through medium-only wells measurements. CC_50_ values were then calculated by non-linear regression using GraphPadPrism 9.5.1.

### Genomic DNA extraction and sequencing to validate PERK stable knockouts cell lines

The extraction of genomic DNA from PERK KO was carried out using the NucleoSpin® Tissue kit (Macherey-Nagel; #740952.50) according to manufacturer protocol. Briefly, Huh7 cells with PERK KO were seeded to around 10 million cells for 24 hours. The cells were suspended in 200 µl buffer T1 (kit provided) and 25 µl of proteinase K solution was added. The sample was then incubated at 70 °C for 15 minutes. Thereafter 210 µl of ethanol was added followed by vigorous vortexing. The sample was then loaded on the kit-provided column for DNA binding and centrifuged for one minute at 11000 x g. Flow-through was discarded and the silica-membrane column was washed with multiple kit-provided buffers (500 µl of buffer AW and 600 µl buffer B5) with centrifugation in-between. DNA was thereafter eluted in 100 µl pre-warmed (at 70 °C) BE buffer (kit provided). Purified genomic DNA was subsequently submitted for targeted sequencing (Eurofins Genomics) using the primer (5’-ATGTTGACCACCAGGGAAAG-3’), designed to amplify the genomic region proximal to the first designated Cas9 cleavage site. Results were then analysed using DNAStar Navigator aligned to the *EIF2AK3* reference sequence to validate InDel mutations.

### Viral N protein alignment

N protein sequences of HCoV-229E (NC_002645) and two other viruses representing the species *Alphacoronavirus chicagoense* (camel alphacoronavirus, KT368903; 229E-related bat coronavirus, KT253269) were downloaded from the Genbank/NCBI database, along with N protein sequences of two members of the species *Alphacoronavirus amsterdamense* (human coronavirus NL63, AY567487; NL63-related bat coronavirus, YP_009328939). A multiple sequence alignment was generated using Clustal Omega (Sievers *et al*, 2011) and rendered using ESPript version 3.0 (Robert & Gouet, 2014). Information on secondary structure elements was derived from crystal structure analyses of the HCoV-NL63 N-and C-terminal domains (Szelazek *et al*, 2017) (pdb 5N4K, 5EPW).

The FASTA-format files containing the amino acid sequences of the following coronaviruses nucleocapsid (N) protein were downloaded from the UniProt database: HCoV-229E (sp|P15130|NCAP_CVH22), MERS-CoV (sp|K9N4V7|NCAP_MERS1), MHV (sp|P03416|NCAP_CVMA5), CoV-NL63 (sp|Q6Q1R8|NCAP_CVHNL), CoV-OC43(sp|P33469|NCAP_CVHOC), SARS-CoV (sp|P59595|NCAP_SARS), SARS-CoV-2 (sp|P0DTC9|NCAP_SARS2) and TGEV (sp|P04134|NCAP_CVPPU). Phylogenetic tree and percent identity matrix of aligned amino acid residues of different viral N proteins were visualized using DNASTAR navigator alignment software, version 11.1.0.54.

### Microarray transcriptomics and subsequent analysis

Purified total RNA from Huh7 cells was amplified and Cy3-labeled using the LIRAK kit (Agilent) following the kit instructions. Per reaction, 200 ng of total RNA was used. The Cy3-labeled aRNA was hybridized overnight to 8x60K 60-mer oligonucleotide spotted microarray slides (Agilent Technologies, design ID 072363). Hybridization and subsequent washing and drying of the slides were performed in accordance with the Agilent hybridization protocol. The dried slides were scanned at 2 µm/pixel resolution using the InnoScan is900 (Innopsys, Carbonne, France). Image analysis was performed with Mapix 8.7.2 software, and calculated values for all spots were saved as GenePix results files.

Stored data were evaluated using the R software and the limma package from BioConductor. Mean spot signals were background-corrected with an offset of one using the NormExp procedure on the negative control spots. The logarithms of the background-corrected values were quantile-normalized. The normalized values were then averaged for replicate spots per array. From different probes annotated the same NCBI gene ID, the probe showing the maximum average signal intensity over the samples was used in subsequent analyses. Genes were ranked for differential expression using a moderated t-statistic. Pathway analyses were done using gene set tests on the ranks of the t-values. Pathway annotations were obtained from KEGG (Kanehisa et al, 2016). The genes assigned to these annotations, including the signal intensity values (E values) and the differential expression between samples (Log_2_ fold change, LFC) with the associated significance (-Log_10_ p value) were listed in an Excel file and used for further filtering steps.

### Mass spectrometry analysis of proteins and protein modifications

#### Exploratory PTM analyses

Huh7 cells were left untreated or were infected with HCoV-229E (MOI=1) for 12 h or 24 h. Cells were washed twice with PBS (without Ca/Mg), and directly lysed on the plate upon addition of fresh urea lysis buffer (20 mM HEPES (pH 8), 9 M urea, 1 mM activated sodium orthovanadate, 2.5 mM sodium pyrophosphate, 1 mM β-glycerophosphate. The cells were carefully resuspended without foaming and extracts were sonified with a Branson device for 4 x 30 s. Samples were centrifuged for 15 min at 4°C and 20,000 x g, and the protein concentration was determined in the supernatant using a Bradford assay. 12 to 13 mg of frozen cell extract were transferred to Cell Signaling Technology (Danvers, MA 01923) for commercial analysis of post-translational modifications by the PTMScan® Discovery proteomics service.

Samples were reduced at 55°C with 4.5 mM dithiothreitol for 30 min, followed by alkylation with iodoacetamide (19 g/l H_2_O) for 15 min at room temperature in the dark. After 1:4 dilution with 20 mM HEPES (pH 8.0), the proteins were digested overnight with 10 g/ml trypsin-TPCK (tolylsulfonyl phenylalanyl chloromethyl ketone). Digested peptides were acidified with 1% trifluoroacetic acid (TFA) and desalted over 360-mg Sep Pak Classic C18 columns (catalog no. WAT051910; Waters, Milford, MA). Elution of peptides was performed with 40% acetonitrile in 0.1% TFA, followed by drying under vacuum. Modified acetylated, ubiquitinylated or phosphorylated peptides were enriched from 0.5 to 1 mg of protein extract by immunoprecipitation using anti acetyl-lysine antibody (#13420, Cell Signaling Technology (Danvers, MA 01923)) or anti ubiquitin branch motif (KεGG) antibody (#3925 Cell 21 Signaling Technology (Danvers, MA 01923) or by IMAC-Fe3+ columns (Cell Signaling Technology (Danvers, MA 01923), respectively.

LC-MS/MS analysis was done using a LTQ-Orbitrap-Elite instrument. Peptides were loaded directly onto a PicoFrit capillary column (10 cm by 75 µm) packed with Magic C18 AQ reversed-phase resin. The column was developed by using a 90 to 150-min linear gradient of acetonitrile in 0.125% formic acid delivered at 280 nl/min. The MS parameter settings were as follows: MS run time, 96 or 156 min; MS1 scan range, 300.0 to 1,500.00; and top 20 MS/MS (min signal, 500; isolation width, 2.0; normalized coll. energy, 35.0; activation-Q, 0.250; activation time, 20.0; lock mass, 371.101237; charge state rejection enabled; charge state 1 +rejected; dynamic exclusion enabled; repeat count, 1; repeat duration, 35.0; exclusion list size, 500; exclusion duration, 40.0; exclusion mass width relative to mass; exclusion mass width, 10 ppm). MS/MS spectra were evaluated using SEQUEST (Eng *et al*, 1994). Each sample was run twice non-sequentially on the instrument both to maximize the number of identifications and also to provide a basis for measurement of analytical reproducibility on the instrument. For each identified peptide, % CV (percent coefficient of variation) values were generated by dividing the standard deviation of the replicate peak area measurements by the average peak area and converting to percentage. For each peptide and fold change reported, the individual % CV values for each sample can be found in **Supplementary Table 1**, as indicated by the headers. Some peptides map to more than one protein due to sequence identity. In these cases, to generate the most complete view of the data, we have listed all possible matching proteins separated by semicolon. No approach was taken to try and resolve to a single protein for these cases since, in the absence of further criteria, it is impossible to unambiguously assign the peptide to one specific protein isoform or paralog. Modification sites are reported as determined by the SEQUEST search. For each peptide, the site localization scores (AScore) were calculated. Score values of 13 were considered of sufficient confidence to correctly assign a modification (Beausoleil *et al*, 2006). Searches were performed against NCBI homo sapiens and coronavirus databases, versions 2015, with a mass accuracy of ± 50 ppm for precursor ions and 1 Da for product ions. Results were filtered with a mass accuracy of 5 ppm on precursor ions and the presence of the intended motif. A 5% default false-positive rate was used to filter the Sequest results. The results were given as spreadsheets with information on the peptide measurements. There are cases where more than one peptide matches to a LC-MS/MS peak due to ambiguous site localization, and in those cases both possibilities are reported and flagged in the data with red text in the intensity columns.

Data were filtered for acetylation, ubiquitination and phosphorylation sites of human viral proteins as reported in **Fig. 1A-B and Supplementary Fig. 1A-B.**

The centered 32 P-sites of HCoV-229E proteins **(Fig. 1B)**, including the seven surrounding N- and C- terminal amino acids, were scored for ranked phosphorylating protein kinases and kinase families online using the human kinase-library, v1.5.0 (library.phosphosite.org/kinase-library/score-site). The top 3 kinases were selected.

#### Parallel proteome/phospho-proteome analyses

Total cell lysates used for parallel proteome or phospho-proteomics mass spectrometry experiments were prepared as follows. Cells were scraped in ice-cold PBS and pelleted at 500 x g for 5 min at 4°C and stored in liquid N2. After thawing, cell pellets from all biological replicates were lysed together (at the same time) using urea-based lysis buffer composed of 20 mM HEPES, pH 8.0, 9.0 M urea (Thermo Scientific, Sequanal Grade, #29700), 1 mM activated sodium orthovanadate, 2.5 mM sodium pyrophosphate, 1 mM ß-glycerophosphate, and water for volume adjustments. The samples were then sonicated using Bioruptor sonicater for 4 min with 30 sec on; 30 sec off, high power settings and centrifuged for 15 min (4°C) at 20000 x g. Supernatants were then transferred to a fresh tubes and protein concentrations were determined with the detergent compatible Bradford assay kit (Pierce™, #23246) using a 300-fold dilution.

Proteomes or phospho-proteomes were determined starting from the same cell extracts. For phospho-peptide enrichment, aliquots containing 600 µg of protein were prepared. 8 M urea was added to each to a final volume of 300 µl. If necessary, the pH was adjusted to 7–9 using 0.1 M HCl or NaOH. Subsequently, 3.75 µl of a 0.2 M TCEP solution in 0.1 M NH₄HCO₃ was added, and the samples were then incubated for 1 hour at 37 °C with shaking at 1000 rpm.

Samples were then cooled down to 25 °C, followed by the addition of 3.75 µl of a 0.4 M iodoacetamide solution in bidistilled water. Incubation was performed in the dark at 25 °C for 45 minutes under gentle shaking (500 rpm). Afterwards, 3.75 µl of a 0.5 M N-acetylcysteine solution in 0.1 M NH₄HCO₃ was added, and the samples were incubated for 10 minutes at 25 °C, 500 rpm. If necessary, the pH was readjusted to 8–9.

Thereafter, the urea concentration was reduced to 6 M by dilution with 0.1 M NH₄HCO₃ and, 7.5 µl of a 0.2 µg/µl LysC solution in 0.1 M NH₄HCO₃ and 1 µl Turbonuclease (Sigma-Aldrich, T4330; 200 Units) were added. Samples were incubated for 3 hours at 37 °C.

Afterwards, the samples were diluted with 0.1 M NH₄HCO₃ to a final urea concentration of 1.6 M. 12 µl of a 0.5 µg/µl solution of Sequencing Grade Modified Trypsin (Serva) in 0.1 M NH₄HCO₃ were added, followed by overnight incubation at 37 °C.

TFA was added to each sample to a final concentration of approximately 1%, lowering the pH below 2. Samples were then centrifuged at 14,000 x g.

Peptides were desalted and concentrated using Chromabond C18WP spin columns (Macherey-Nagel, Part No. 730522). The dried eluates were dissolved in 1 mL of water containing 5% acetonitrile and 0.1% formic acid. Peptide concentrations were determined using a NanoDrop spectrophotometer (Thermo Scientific). A small aliquot (less than 5 µg) was set aside for whole-proteome analysis and diluted to a concentration of 100 ng/µl. The remaining samples were dried in a vacuum concentrator (Eppendorf).

Phospho-peptide enrichment was performed using magnetic MagReSyn Ti-IMAC beads (PELOBiotech) following the manufacturer’s instructions. Per sample, 50 µl (of beads equivalent to 1 mg of binding capacity) were freshly prepared. Beads were stored at 4 °C for no longer than six months and not frozen. Beads were thoroughly resuspended by vortexing, and 50 µl were transferred to 2 mL microcentrifuge tubes. After placing the tubes on a magnetic rack, the storage solution was removed. Beads were washed twice with 200 µl of 70% ethanol by vortexing, shaking at 600 rpm for 5 minutes, placing on the rack, and removing the supernatant. Subsequently, beads were treated with 100 µL of 1% ammonium hydroxide, vortexed, placed on the magnetic rack, and then the supernatant was discarded. This was followed by three washes with 50 µl of loading buffer (1 M glycolic acid, 80% acetonitrile, 5% TFA), each including a 60-second incubation at room temperature, magnetic separation, and removal of the supernatant.

Dried peptide samples were then reconstituted in 100 µl of loading buffer and dissolved in an ultrasonic water bath. The peptide solution (corresponding to ~600 µg, see above) was added to the equilibrated beads, vortexed, and incubated for 20 minutes at room temperature with shaking at 400 rpm. Beads were collected on a magnetic rack, and the supernatant was discarded. Unbound peptides were removed by washing with 100 µl of loading buffer, followed by vortexing, magnetic separation, and removal of the supernatant. This was followed by two washes with 100 µl of wash buffer (80% acetonitrile, 1% TFA) with 2 minutes shaking at 400 rpm and magnetic separation. Beads were additionally washed twice with 100 µL of 10% acetonitrile/0.2% TFA by vortexing, magnetic separation, and removal of the supernatant.

Phospho-peptides were eluted by adding 80 µl of 1% aqueous ammonium hydroxide to the beads, followed by a 15-minute incubation at room temperature. After magnetic separation, the supernatant was collected in fresh tubes. This elution step was repeated twice, and the three eluates were combined to a total volume of 240 µl. The combined eluates were acidified with 60 µl of 10% formic acid.

#### Nascent proteome analyses

OPP-samples were digested on-beads by adding 0.1 µg trypsin in 50 µl ammonium-bicarbonate buffer to the beads and samples were incubated at 37°C for 45 minutes. Subsequently, the supernatant was transferred into fresh tubes and digested overnight at 37°C to completeness.

For the reduction of disulphide bridges, 5 mM DTT (Dithiothreitol) was added. Samples were then incubated for 15 minutes at 95°C. Subsequently, the resulting sulfhydryl-groups were chemically modified by adding iodoacetamide to a final concentration of 25 mM and incubating samples for 45 minutes at room temperature (RT) in the dark. Excess iodoacetamide was quenched by the addition of 50 mM DTT and incubation for one more hour at RT.

### Mass spectrometry of tryptic peptide mixtures

Peptide, phospho-peptide or nascent peptide mixtures were desalted and concentrated using Chromabond C18WP spin columns. Finally, dried peptides were dissolved in 15 µl of water containing 5% acetonitrile and 0.1% formic acid. Peptide concentrations were determined using a NanoDrop spectrophotometer.

Mass spectrometric analysis was performed in technical duplicates on a timsTOF Pro mass spectrometer (Bruker Daltonics) coupled to a nanoElute 2 HPLC system (Bruker Daltonics). An Aurora Ultimate C18 RP column (25 cm × 75 µm, 1.7 µm beads; IonOpticks) was used. Approximately 200 ng of peptides (2 µl) were injected directly onto the column. Sample loading was performed at a constant pressure of 800 bar. Separation was carried out at 60 °C with a gradient of water/0.1% formic acid (solvent A) and acetonitrile/0.1% formic acid (solvent B) at a flow rate of 400 nl/min: linear increase from 2% to 17% B over 60 minutes, from 17% to 25% B over 30 minutes, and from 25% to 35% B over 10 minutes. Subsequently, solvent B was increased to 95% within 10 minutes and held for an additional 10 minutes.

The built-in “DDA PASEF-standard_1.1sec_cycletime” method (Bruker Daltonics) was applied for data acquisition. Raw data analysis was conducted using MaxQuant version 2.5.1.0.7 (Tyanova *et al*, 2016a). For peptide/protein assignments, two FASTA databases were used. A human UniProt FASTA database from July 2024 and a UniProt FASTA database from March 2021, representing human coronavirus 229E, TaxonId 11137, where the protein sequence of the polyprotein R1AB was split into 16 posttranslationally processed nsp proteins.

For the detection of phosphorylated peptides, the raw data were processed using the same settings as before plus an additional search for phospho (STY) sites as provided by MaxQuant and the results were extracted from the MaxQuant output files “modificationSpecificPeptides.txt”, “Phospho (STY)Sites.txt” and “proteinGroups.txt”.

### Bioinformatic analyses

Perseus software (version 1.6.15) was used for further analyses of protein intensity values (Tyanova *et al*, 2016b). IDs only identified by site, reverse sequences and contaminates were omitted from further analysis.

For nascent proteome analyses, measurements from cell lysates of 24 samples and the according 24 purified nascent protein fractions were combined into one Perseus analysis file, resulting in 96 measurements, including technical duplicates, and 6505 protein IDs. Samples from lysates were normalized by width adjustment, whereas streptavidin-purified nascent proteomes were not normalized to preserve the anticipated differences between conditions, but in this protein set missing values were imputed using a Log_2_ intensity value of 9, which was below the lowest intensity value measured across all samples.

For proteome analyses, measurements from 32 cell lysates were combined into one Perseus analysis file, resulting in 64 measurements, including technical duplicates, and 6488 protein IDs. Samples were normalized by width adjustment.

For phospho-proteome analyses, measurements from 32 cell lysates were combined into one Perseus analysis file, resulting in 64 measurements, including technical duplicates, and 12766 unique P-peptide IDs, which were collapsed to 12585 ID by omitting contaminants, reverse and only identified by site sequences. Samples were normalized by width adjustment and missing values were replaced from normal distribution using Perseus settings (width 0.3, down shift 1.8).

All protein intensity values were Log_2_-transformed. For calculation of ratio values, biological and technical replicates from each condition were assigned to one analysis group using Perseus tools for categorical annotation rows.

Enriched proteins between pair-wise comparisons were identified by Student’s t-tests using Perseus functions and were visualized by Volcano plots.

Interesting groups of regulated proteins/P-proteins were defined by Log_2_ fold change (LFC) and statistical significance of changes based on -Log_10_ p values ≥ 1.3 as indicated in the Figure legends.

Subsequent filtering steps and heatmap visualizations were performed in Excel 2016 according to the criteria described in the figure legends.

Venn diagrams were created with tools provided at http://bioinformatics.psb.ugent.be/webtools/Venn/.

Overrepresentation analyses of gene sets were done using the majority protein IDs or gene IDs of differentially enriched proteins or mRNAs uploaded to Metascape software and processed with the predefined express settings (Zhou *et al*, 2019).

### Kinome motif analyses, kinome annotations and visualizations

First, centered motifs containing the phosphorylated residue plus the seven surrounding N- and C- terminal amino acids for 303 human serine/threonine kinases assigned to 89784 P-sites from the human kinome motif atlas were mapped onto the Huh7 and A549-ACE2 phospho-proteomes including the information on ranks and percentiles of the top three protein kinases assigned to each motif by previous systematic in vitro kinase assays as published by Johnson et al. (Johnson *et al*., 2023).

Second, to map protein kinase motifs within identified P-proteomes from Huh7 **(Fig. 6)** or A549-ACE2 cells **(Fig. 7)**, and deduce sets of activated or suppressed protein kinases by kinase frequency enrichment analysis, tools from the kinase library, version 0.011 (Huh7) or 1.0.0 (A549-ACE2), provided at (https://kinase-library.mit.edu/home) by the Yaffe and Cantley labs, respectively, were used as published in (Johnson *et al*., 2023). 10952 (Huh7 cells) or 5377 (A549-ACE2 cells) centered P-peptide motifs, along with Log_2_ ratio and –Log_10_ p values for each pairwise condition of interest, were used as input data sets for phospho-proteomic enrichment analyses. For Huh7, the following analysis parameters were applied: Foreground LFC: 1, p value threshold: 1.3, serine/threonine kinases, Benjamini-Hochberg procedure for FDR, include Ser vs Thr favorability for the central phospho-acceptor and non-canonical kinases. For A459-ACE2 cells, the settings were the same except that LFC threshold was set to 0.585. The output data were transferred to Excel and Log_2_ transformed dominant enrichment and -Log_10_ transformed dominant p values were plotted as Volcano plots **(Fig. 6C-F)** and used to define activated (Log_2_ ratio > 0, -Log10 p ≥ 1.3) or inhibited (Log_2_ ratio < 0, -Log10 p ≥ 1.3) sets of protein kinases across all conditions.

Note, that the human kinase library is constantly updated and a history is available at (https://pypi.org/project/kinase-library/1.5.0/).

For annotations of protein kinases to the kinome, we used the list of 536 protein kinases and their associated families, groups and alternative names as published in the kinhub data base (http://www.kinhub.org/)(Eid *et al*., 2017). This lists builds on the orginal kinome of 518 protein kinases published by Manning et al. (Manning *et al*., 2002). The nomenclature of kinase families **(Supplementary Fig. 5)** was further adapted from the kinomer resource (https://www.compbio.dundee.ac.uk/kinomer/families.html) and from resources available at Kinome.org which lists 716 human protein kinases (https://labsyspharm.shinyapps.io/kinome/). The kinome annotations used in this study are summarized in **Supplementary Table 10**.

For projection of kinases expression or activity values (i.e. dominant enrichment values) onto nodes and branches of the original phylogenetic kinome tree **(Fig. 6, Supplementary Fig. 5 and 9)**, online tools from Coral were employed (http://phanstiel-lab.med.unc.edu/CORAL/) (Metz *et al*, 2018).

De novo sequence motif logos from lists of regulated and non-regulated P-peptides **(Supplementary Fig. 5, 10)** were generated from lists of centered P-peptides at PosphoSitePlus online (https://www.phosphosite.org/sequenceLogoAction), focusing on phospho-serine and frequency changes as analysis parameters (Hornbeck *et al*., 2019). Motif logos for the 303 human Ser/Thr kinases as reported in the human kinome motif atlas were downloaded from the supplement of Johnson et al. (Johnson *et al*., 2023).

Protein networks were constructed using Cytoscape, version 3.10.0 or higher (Shannon *et al*, 2003).

### Quantification and statistical analysis

Protein bands detected by Western blotting were quantified using Bio-Rad Image Lab, version 5.2.1 (build 11) or version 6.1.0 (build 7). Statistics (t-tests, Mann-Whitney-Rank Sum Test, one-way ANOVA, Kruskal-Wallis tests, correlations) were calculated using GraphPad Prism 9.5.1, Perseus 1.6.14.0 or Microsoft Excel 2016.

All statistical tests for pathway enrichment analyses were calculated online by Metascape software (https://metascape.org/) (Zhou *et al*., 2019) using the ontology sources KEGG Pathway, GO Biological Processes, Reactome Gene Sets, Canonical Pathways, CORUM, TRRUST, DisGeNET, PaGenBase, Transcription Factor Targets, WikiPathways, PANTHER Pathway, and COVID and all genes in the genome as the enrichment background. P-values were based on the accumulative hypergeometric distribution and q-values were calculated using the Benjamini−Hochberg procedure to account for multiple testings. Terms with a p-value < 0.01, a minimum count of 3, and an enrichment factor > 1.5 were collected. Kappa scores were used as the similarity metric for hierarchical clustering on the enriched terms, and sub-trees with a similarity of > 0.3 were considered a cluster. The most statistically significant terms within a cluster were chosen to represent the cluster.

## Data availability

The nascent proteomic and proteomic/phospho-proteomic data sets of this study have been deposited to the ProteomeXchange Consortium via the PRIDE partner repository (Perez-Riverol *et al*, 2025) with the dataset identifiers PXD068700, project DOI:10.6019/PXD068700, or, PXD069083, project DOI: 10.6019/PXD069083, respectively (https://www.ebi.ac.uk/pride/login/PXD068700, https://www.ebi.ac.uk/pride/login/PXD069083).

The microarray data sets of this study have been deposited in NCBI’s Gene Expression Omnibus (Edgar *et al*, 2002) and are accessible through GEO Series accession number GSE306496 (https://www.ncbi.nlm.nih.gov/geo/query/acc.cgi?acc= GSE306496).

## Acknowledgments

This work was supported by the following grants from the Deutsche Forschungsgemeinschaft (DFG, German Research Foundation). SFB1213/3 (B03 (to M.K.); project 268555672); TRR81/3 (B02 (to M.K.); project 109546710); KR1143/9-2 (KFO309, P3 (to M.K. and J.Z.); project 284237345); KR1143/12-1 (to MK, project 554390251); SFB1021/3 (C02 (to M.K.), Z03 (to M.K., U.L); project 197785619); and GRK2573/2 (RP5 (to M.K.); project 416910386).

Work in the laboratory of M.K. is also supported by the LOEWE program of the state of Hesse (Coropan, P4 (to MK)), the IMPRS program of the Max Planck Society and the Excellence Cluster Cardio-Pulmonary Institute (EXC 2026: Cardio-Pulmonary Institute (CPI), project 390649896) and the DZL/UGMLC program. We are grateful for support from the Margarete Elisabeth Leib-Stiftung. We thank Helmut Muller for technical assistance.

## Author contributions

MSS performed experiments, generated genome-edited cell lines, analyzed data and drafted the Methods section, HW performed and analyzed experiments, UL performed mass spectrometry analysis, AW processed and analyzed MS data, JM-S established OPP labeling protocols, FD performed experiments, CM-B contributed to cytotoxicity assay, MP performed experiments, JZ and NK generated and functionally characterized mutant coronaviruses, JW performed and analyzed microarray experiments, MK conceived the study, performed and visualized all bioinformatics analyses, prepared figures and tables and wrote the initial draft.

---

**Supplementary Fig. 1.**
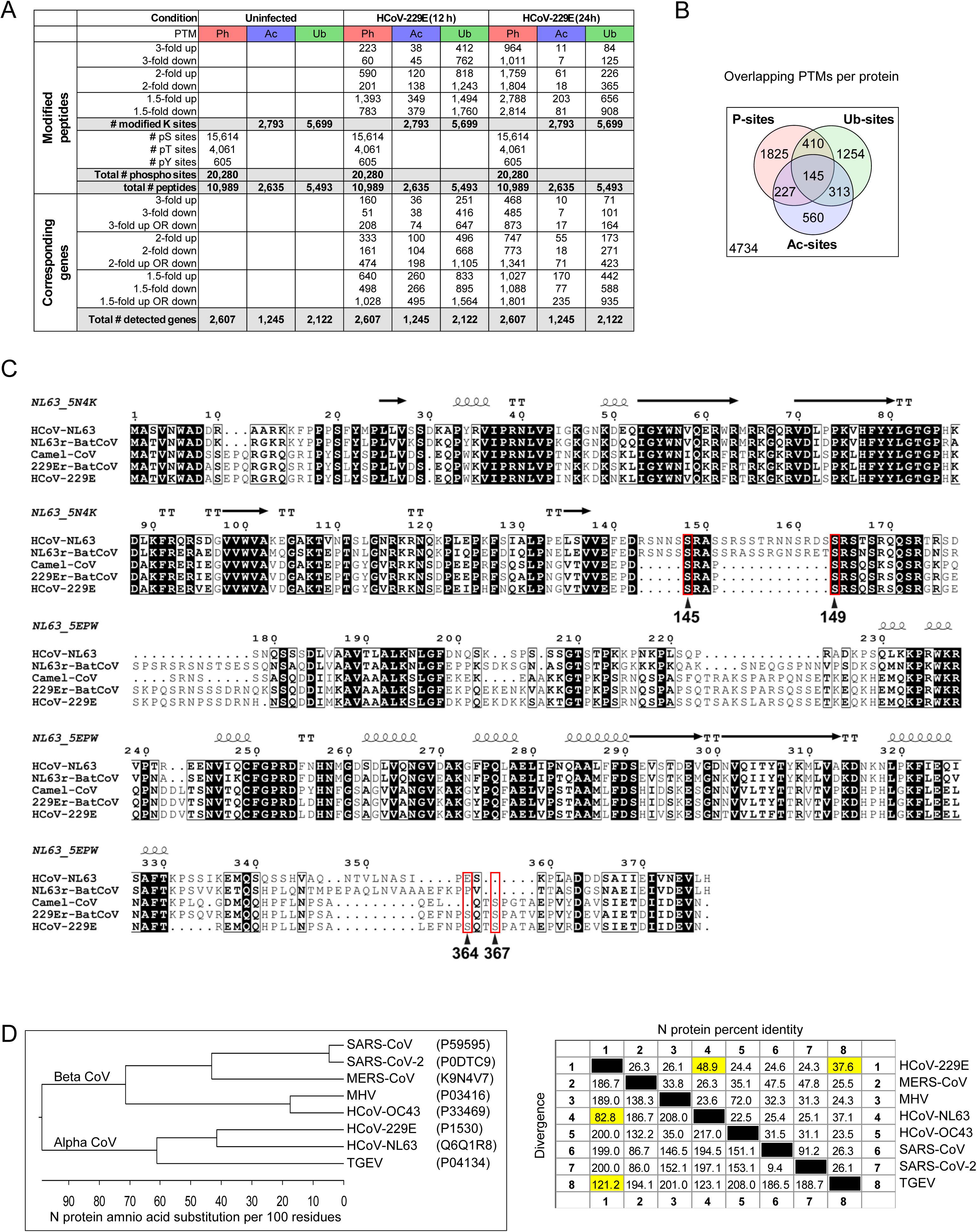
Changes of phosphorylation, acetylation and ubiquitination patterns in HCoV-229E infected cells and alignments of N protein phosphorylation sites. (A) Table summarizing the numbers of detected modified peptides and their regulation in the LC-MS/MS analysis shown in Fig. 1A-B. (B) Venn diagram summarizing overlapping and unique post-translational modifications in Huh7 cells. (C) Amino acid sequence alignment of HCoV-229E N protein with the most closely related HCoV-NL63 and bat CoV variants as indicated by their accession numbers. The alignment was based on primary sequence and secondary structure information from HCoV-NL63 N protein according to the annotation of the top rows. Red boxes mark tandems of phosphorylated serine residues within the conserved serine rich (SR) region in the middle part and the C-terminal region of N protein, respectively, that were investigated in this study. (D) Identity/divergence matrices demonstrating conservation of N protein residues at the amino acid level across different alpha-and betacoronaviruses compared with the N protein of HCoV-229E. Yellow colors highlight the most closely related HCoV-NL63 and TGEV.

**Supplementary Fig. 2.**
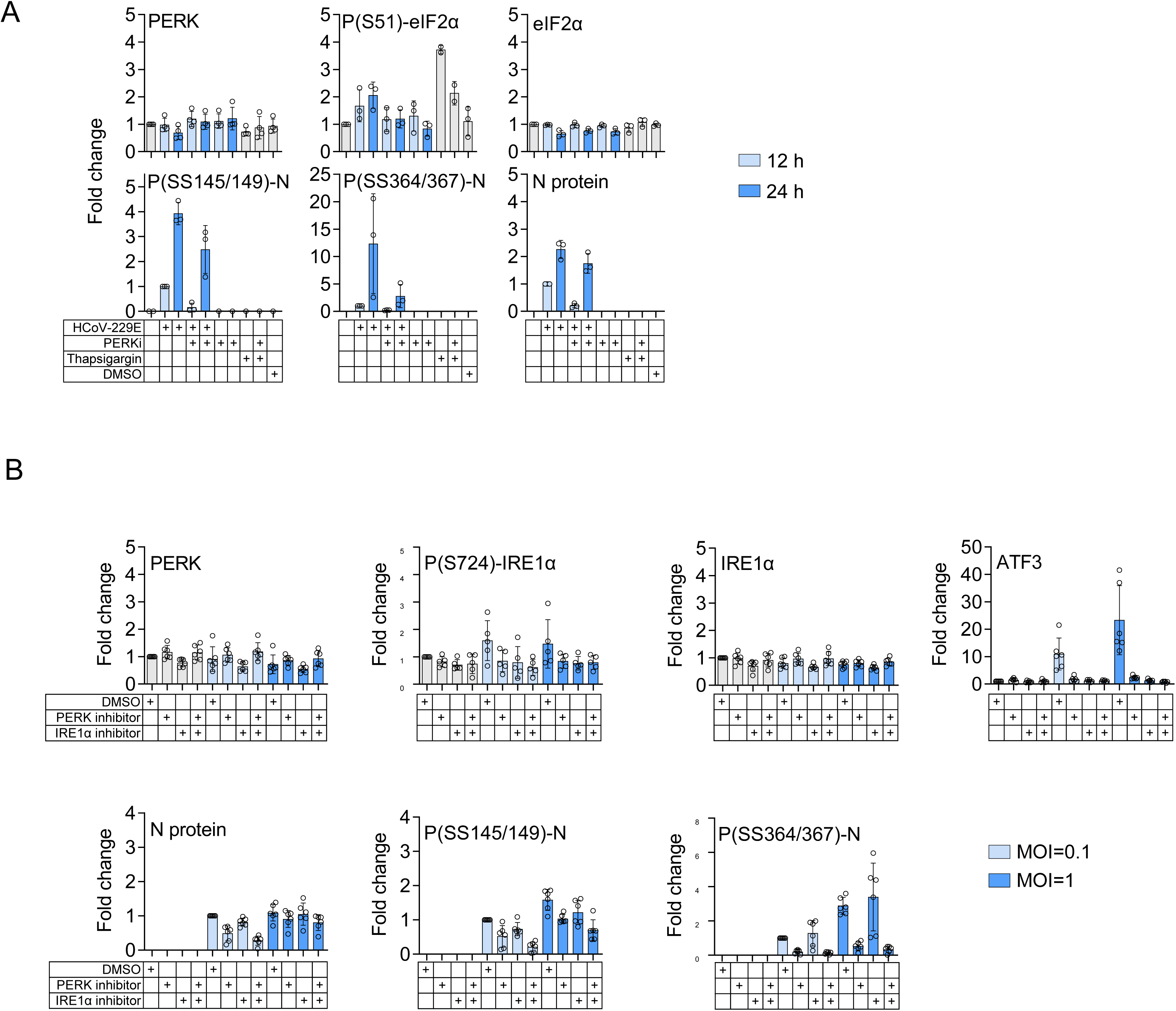
Quantification of effects of PERK inhibition on N protein levels and N protein phosphorylation. (A, B) Quantification of immunoblot experiments shown in Fig. 2B-C. Graphs shown relative mean changes ± s.d. compared to untreated control. Changes of N protein and N protein phosphorylation levels were compared to the 12 hpi condition (A) or cells infected with MOI of 0.1 (B). Data are from three (A) or five (B) biologically independent experiments.

**Supplementary Fig. 3.**
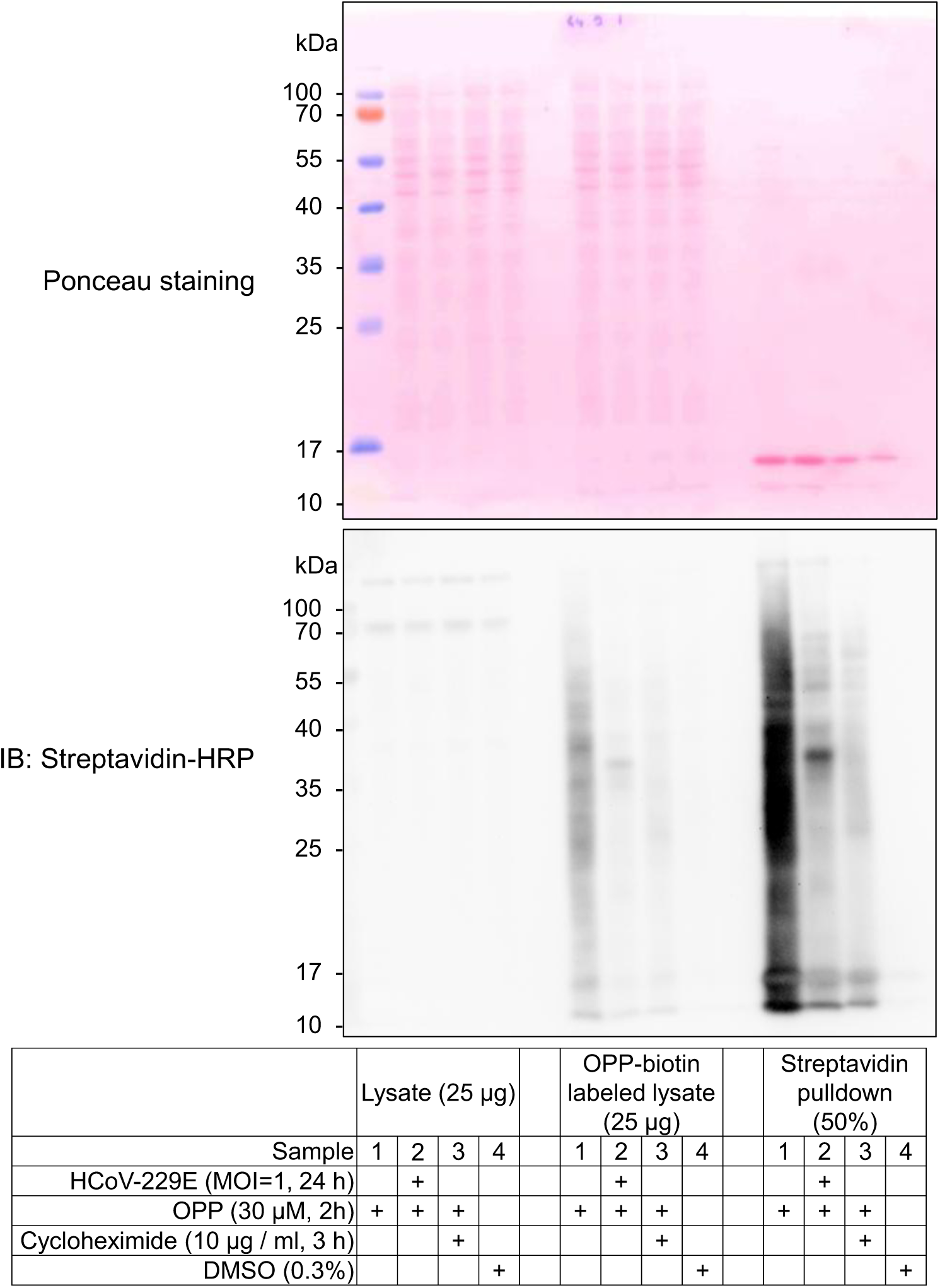
Validation of OPP-labelling in lysates and nascent protein fractions. Huh7 cells were left untreated or were infected with HCoV-229E (MOI=1, 24 h). For the last 2 h, cells were metabolically labeled with OPP (30 µM) or solvent (DMSO) as indicated. Cycloheximide (CHX, 10 µg / ml) was added to one sample 1 h before OPP addition. All samples were lysed simultaneously and total cell extracts were subjected to click biochemistry using biotin azide as substrate. Biotinylated nascent polypeptides were purified by streptavidin affinity chromatography. Lysates before and after the click-chemistry step (OPP-biotin labeled lysate), and purified nascent protein fractions were analyzed for the presence of biotinylated proteins by Western blotting using streptavidin-coupled horseradish peroxidase (HRP). Equal loading was confirmed by Ponceau red staining of the blot membrane.

**Supplementary Fig. 4.**
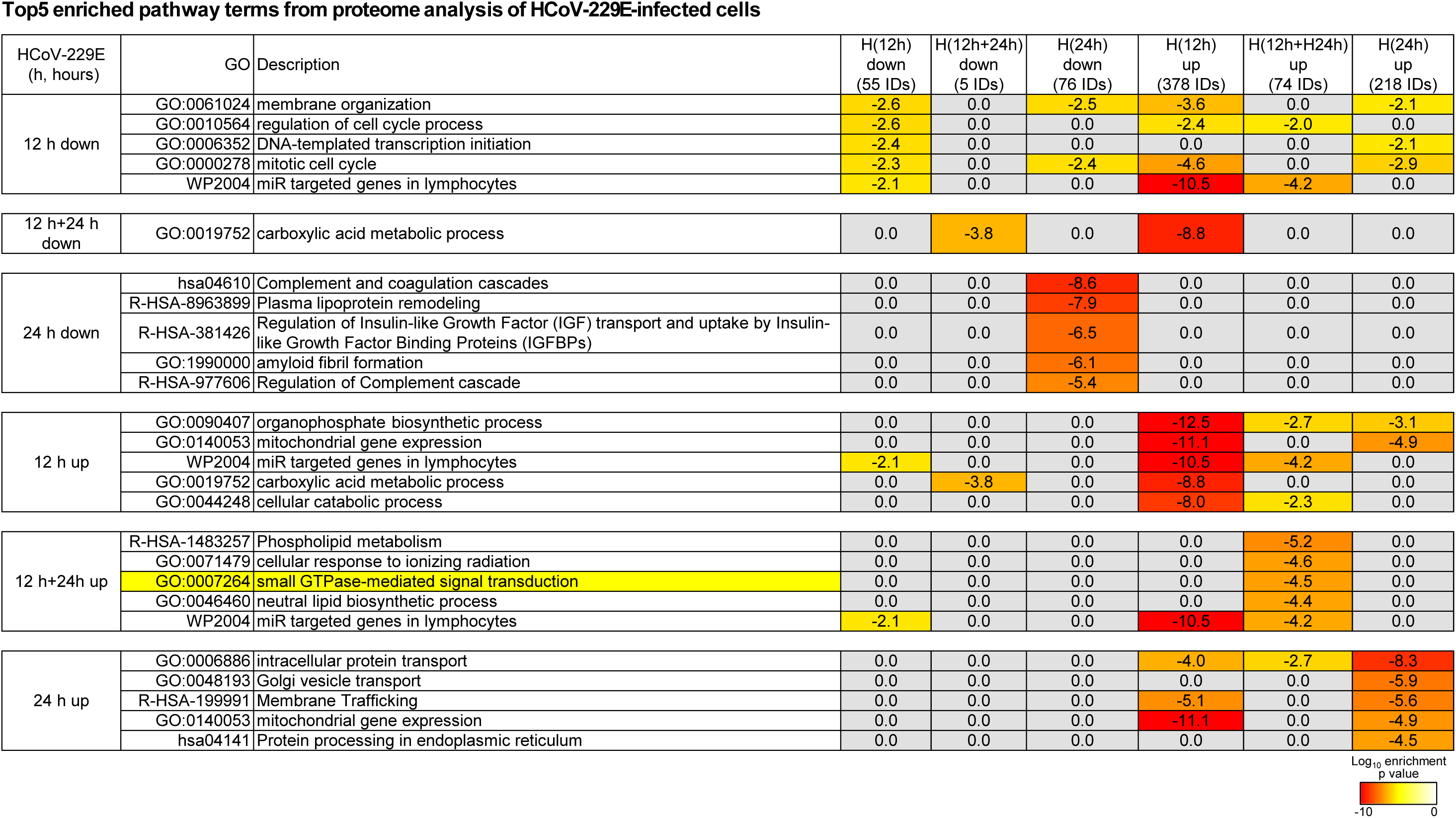
Top5 pathways associated with up- or downregulated proteins in cells infected with HCoV-229E for 12 h or 24. **h.** Metascape (Zhou *et al*., 2019) overrepresentation analysis was performed side by side for the six groups of proteins that were up- or down-regulated in response to HCoV-229E after 12 hpi or 24 hpi as shown in the left Venn diagrams of Fig. 4B. The heatmap shows the top5 most strongly enriched pathway terms for each condition along with the enrichment values of all other conditions. Yellow colors mark enrichment of small GTP-ase pathway components as highlighted in additional data sets of this study.

**Supplementary Fig. 5.**
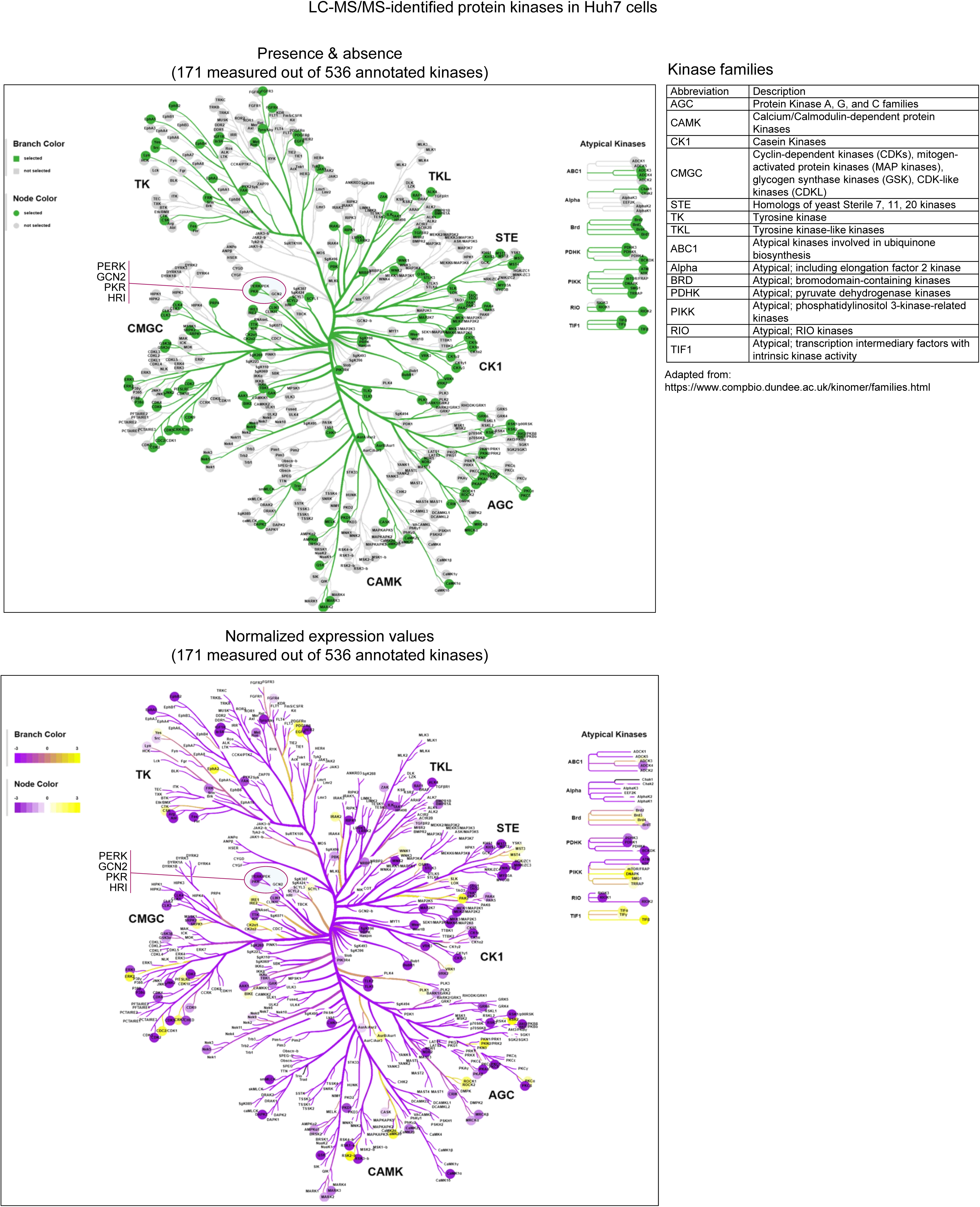
Distribution of protein kinases from Huh7 cells identified at the protein level on the kinome tree. 171 protein kinases out of 536 kinases annotated in the kinhub data base were found to be present in the proteome data sets shown in Fig. 4F (http://www.kinhub.org/index.html) (Eid *et al*., 2017). Coral software was used to project the distribution of these kinases on the kinome tree (green, upper graph) along with their normalized expression levels (yellow to purple, lower graph) using the values from the DMSO (12 h) control samples (http://phanstiel-lab.med.unc.edu/CORAL/) (Metz *et al*., 2018). The table explains the abbreviations of the major kinase families.

**Supplementary Fig. 6.**
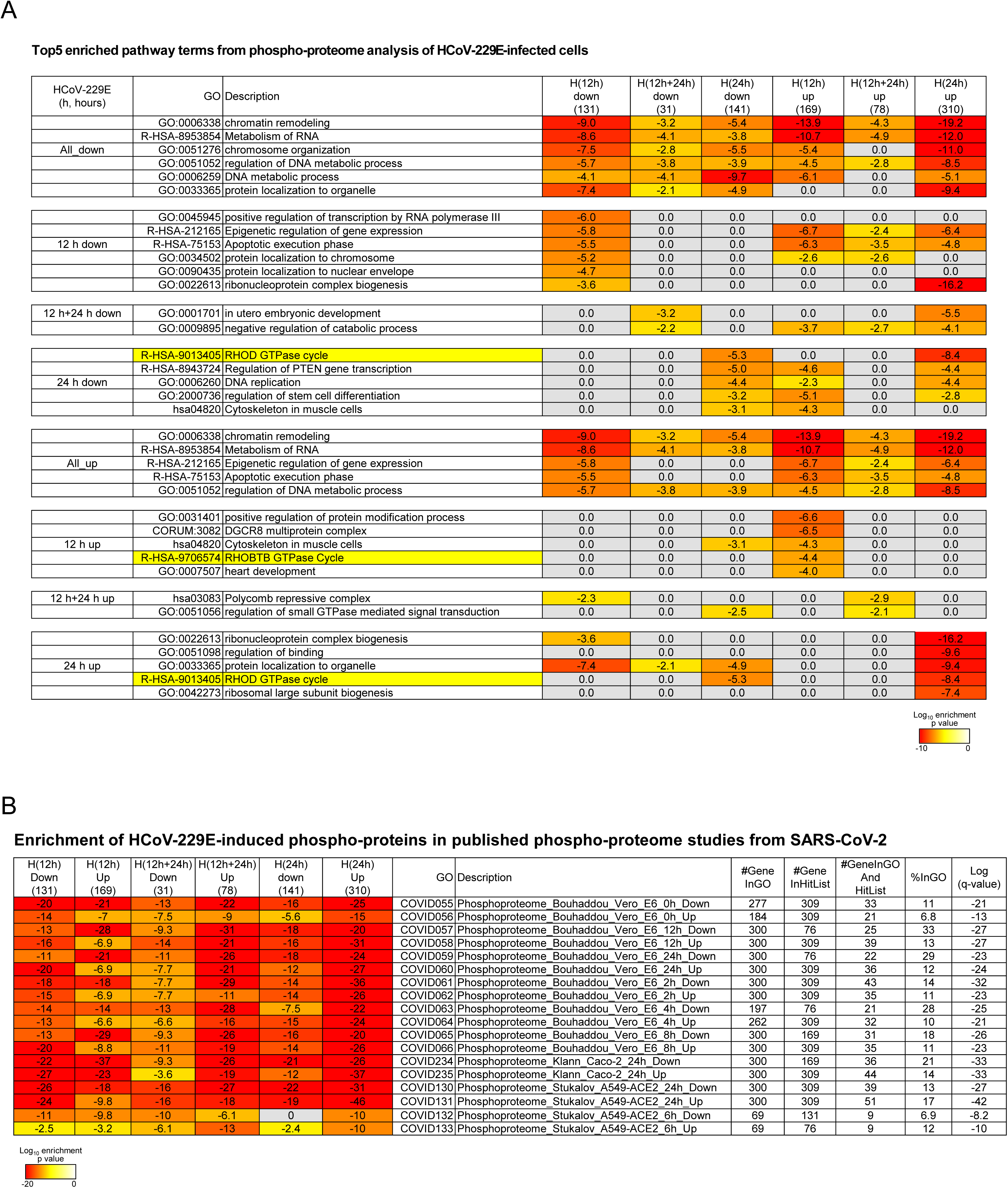
Top5 pathways associated with up- or downregulated P-proteins in cells infected with HCoV-229E for 12 h or 24 h. (A) Metascape (Zhou *et al*., 2019) overrepresentation analysis was performed side by side for the six groups of P-proteins that were up- or down-regulated in response to HCoV-229E after 12 hpi or 24 hpi as shown in the left Venn diagrams of Fig. 5C. The heatmap shows the top5 most strongly enriched pathway terms for each condition along with the enrichment values of all other conditions. Yellow colors mark enrichment of small GTP-ase pathway components as highlighted in additional data sets of this study. (B) Overrepresentation of HCoV-229E-induced or repressed P-proteins in three published P-proteomic data sets according to the same Metascape analysis performed in (A). The heatmap shows the significance of enrichment along with the identified overlapping pathway components and further analysis parameters.

**Supplementary Fig. 7.**
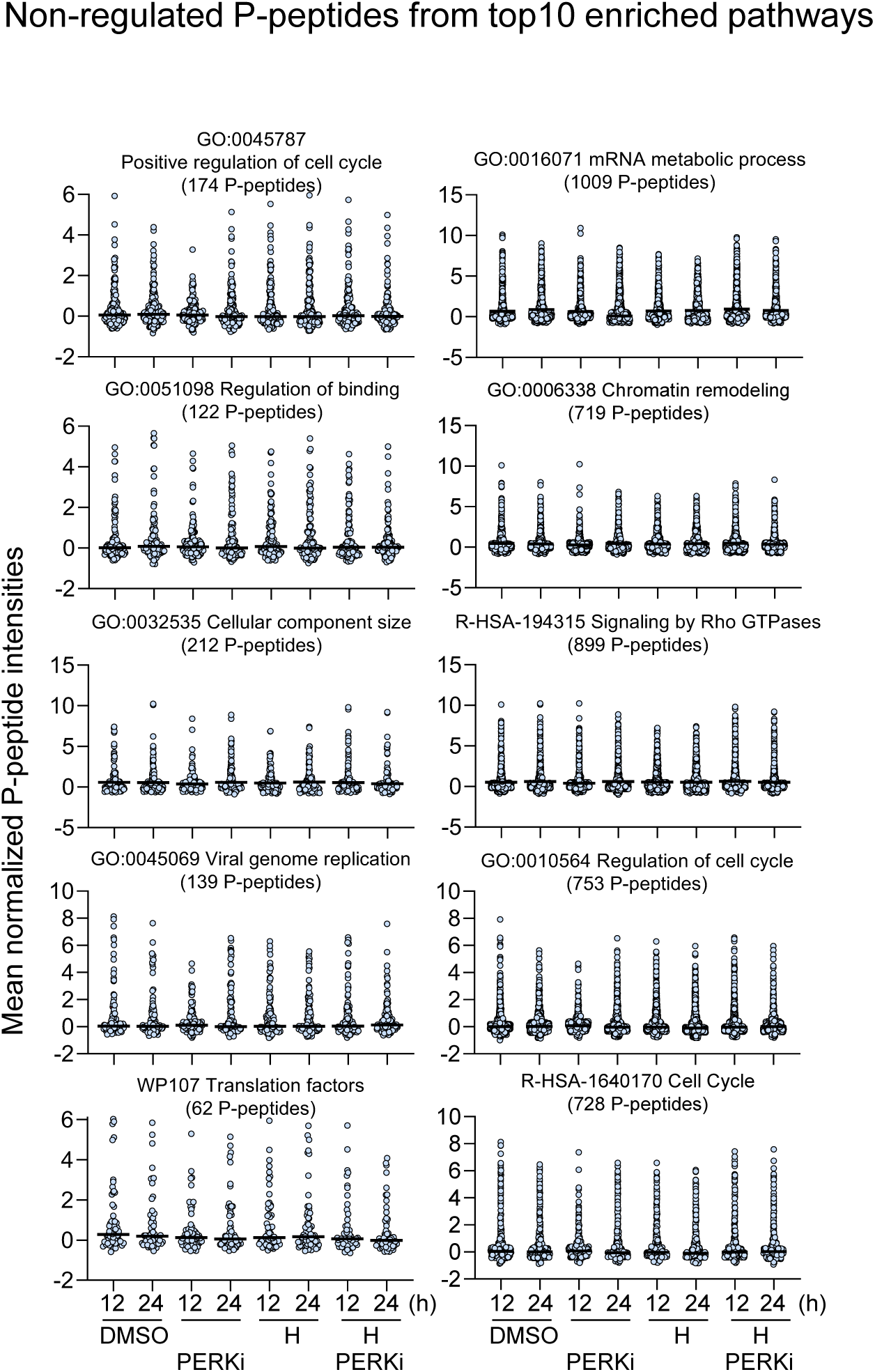
Non-regulated P-peptides from top10 enriched pathway terms. Scatter plots showing mean abundancies of all P-peptides, including non-regulated ones, belonging to the top10 pathways identified in P-proteomes of Huh7 cells as shown in Fig. 5E-F. Data are from four biological replicates and two technical replicates, black lines indicate medians.

**Supplementary Fig. 8.**
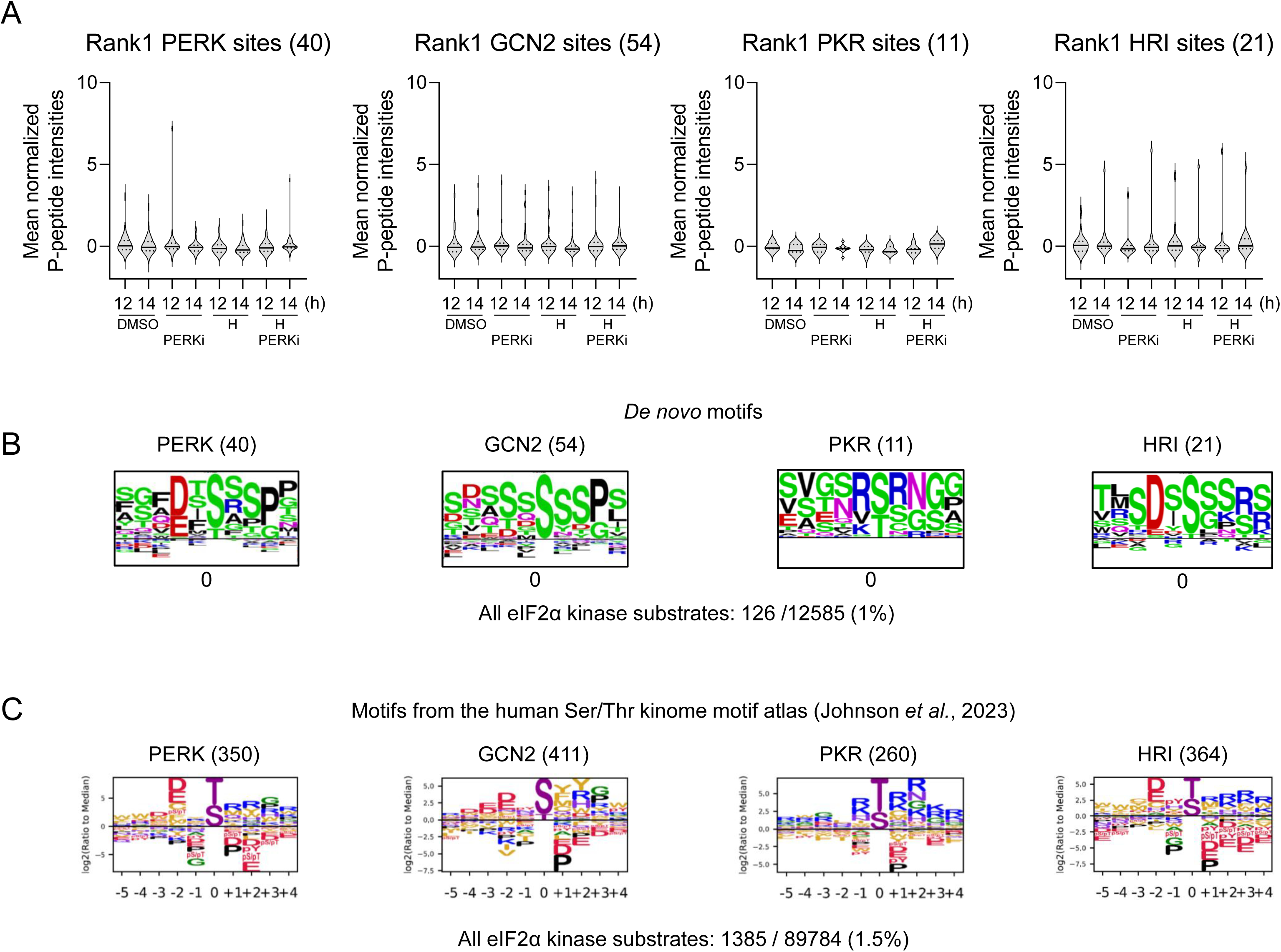
Phospho-proteome analysis of eIF2α kinase substrates. (A) All 12585 P-peptides described in Fig. 4 were analyzed for motifs with PERK (encoded by the *EIF2AK3* gene), GCN2 (encoded by the *EIF2AK4* gene), PKR (encoded by the *EIF2AK2* gene) or HRI (encoded by the *EIF2AK1* gene) as rank1 kinase according to the human motif atlas annotation (Johnson *et al*., 2023). Violin plots show mean P-peptide intensities across all conditions. Solid lines show medians and dashed lines show 1^st^ and 3^rd^ quartiles. Numbers in brackets indicate identified P-peptides per kinase. (B) *De novo* sequence logos of P-peptides, centered at the phosphorylated residue, were generated from the four kinase groups shown in (A) by tools from PhospoSitePlus (https://www.phosphosite.org/sequenceLogoAction) (Hornbeck *et al*., 2019). (C) Motif logo plots showing the preferred consensus phosphorylation motifs of the four kinases as published in the supplement of Johnson et al. for comparison (Johnson *et al*., 2023).

**Supplementary Fig. 9.**
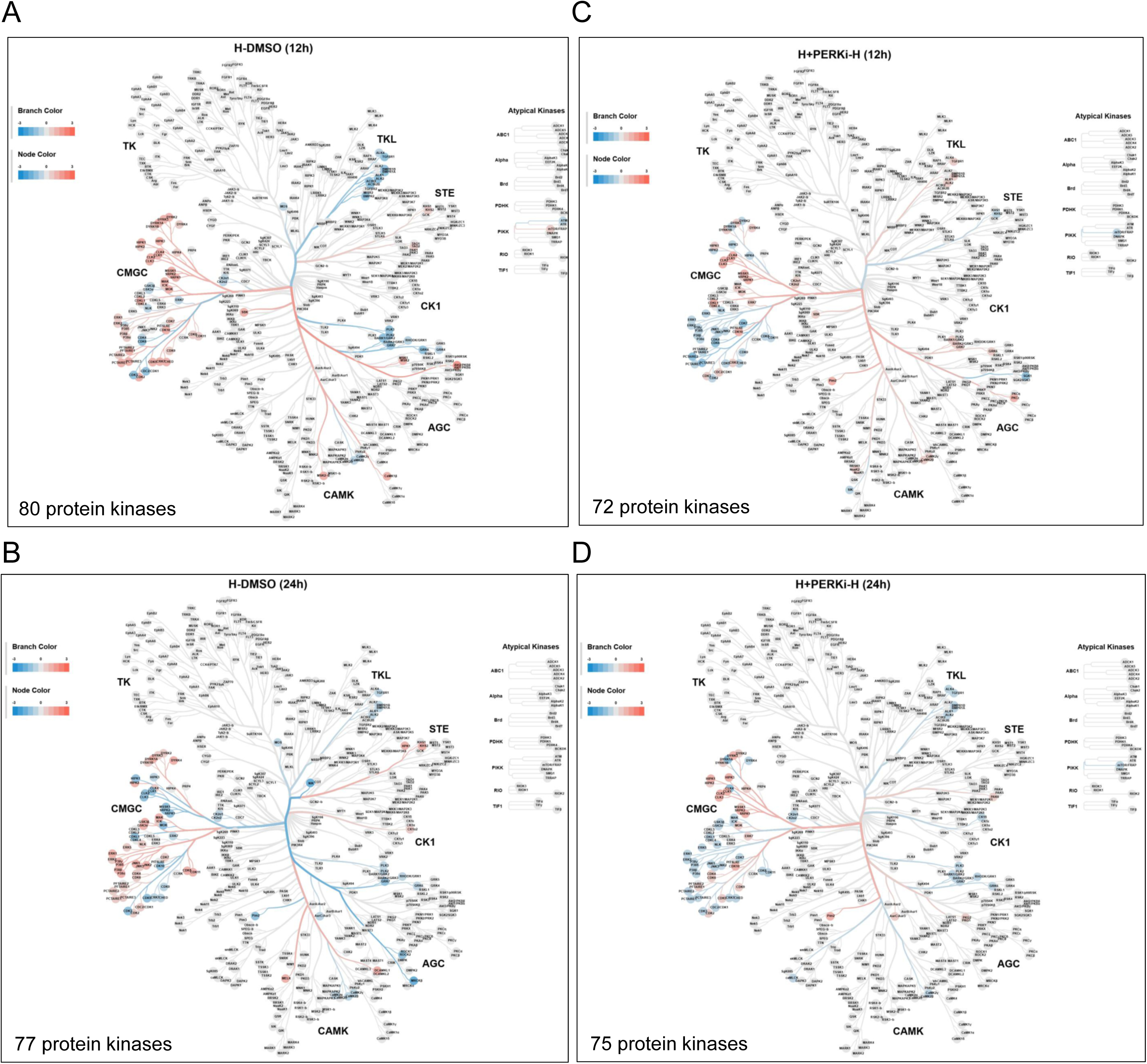
Projection of activated or suppressed protein kinases in HCoV-229E-infected and PERK-treated Huh7 cells on the kinome tree. Protein kinases found to be activated (red colors) or repressed (blue colors) in CoV-infected and PERKi-treated cells by kinome motif enrichment analysis as shown in Fig. 6C were projected onto the kinome tree along with their dominant Log_2_ enrichment values using Coral software tools (http://phanstiel-lab.med.unc.edu/CORAL/) (Metz *et al*., 2018).

**Supplementary Fig. 10.**
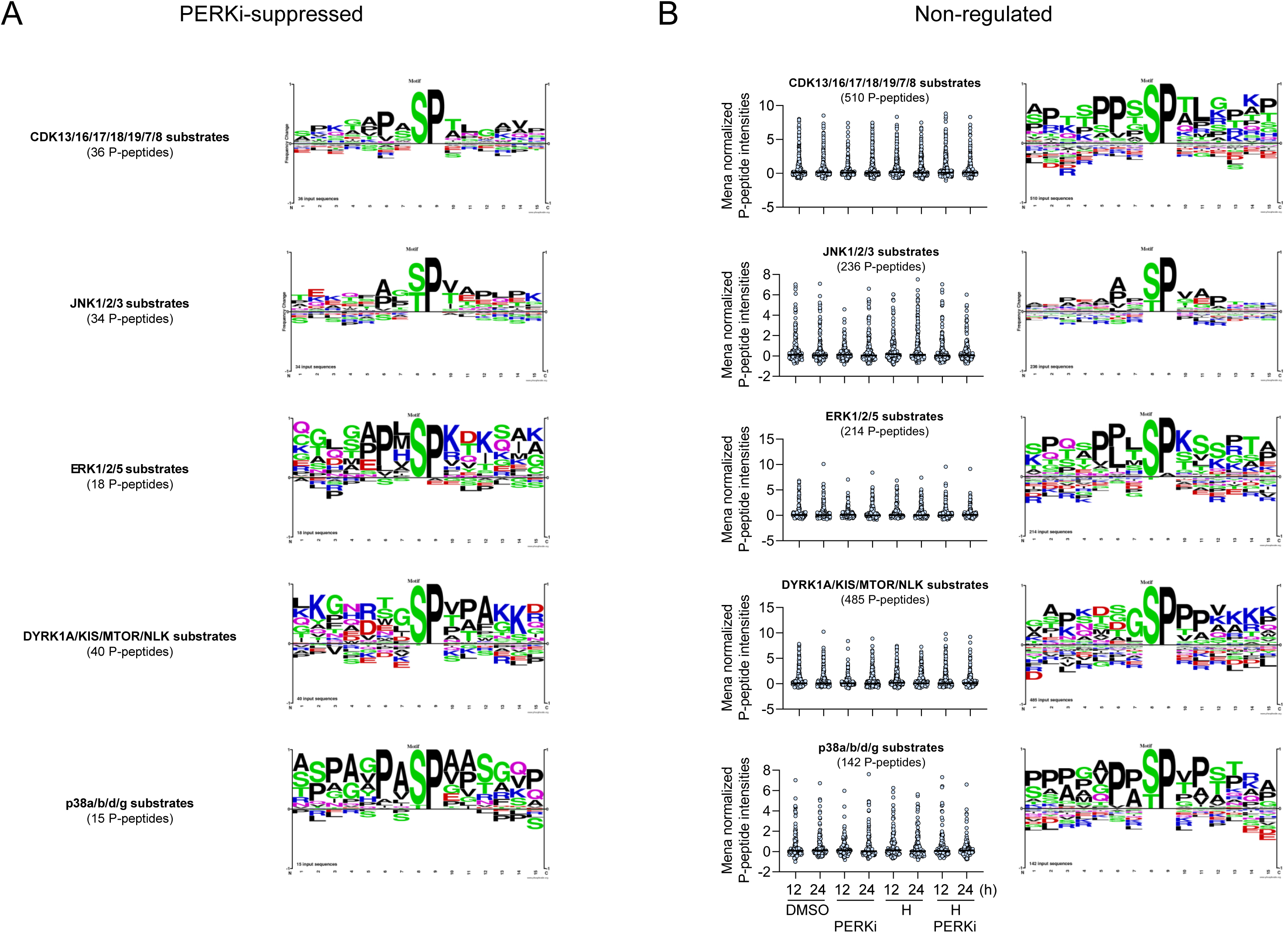
CMGC kinases sequence preferences in regulated and non-regulated sets of P-peptide substrates. (A) *De novo* sequence logos, centered at the phosphorylated residue, of all PERKi-suppressed P-peptides of the five kinase groups shown in Fig. 5I were generated by tools from PhospoSitePlus online (https://www.phosphosite.org/sequenceLogoAction) (Hornbeck *et al*., 2019). (B) Scatter plots showing mean abundancies of all non-regulated P-peptide substrates of the five groups comprising 21 kinases as shown in Fig. 6H-I. For each peptide, the top1-ranked Ser/Thr kinase was selected and the values for all conditions are shown. Data are from four biological replicates, black lines indicate medians. On the right, sequence logos generated as described in (A) are shown for each panel.

**Supplementary Fig. 11.**
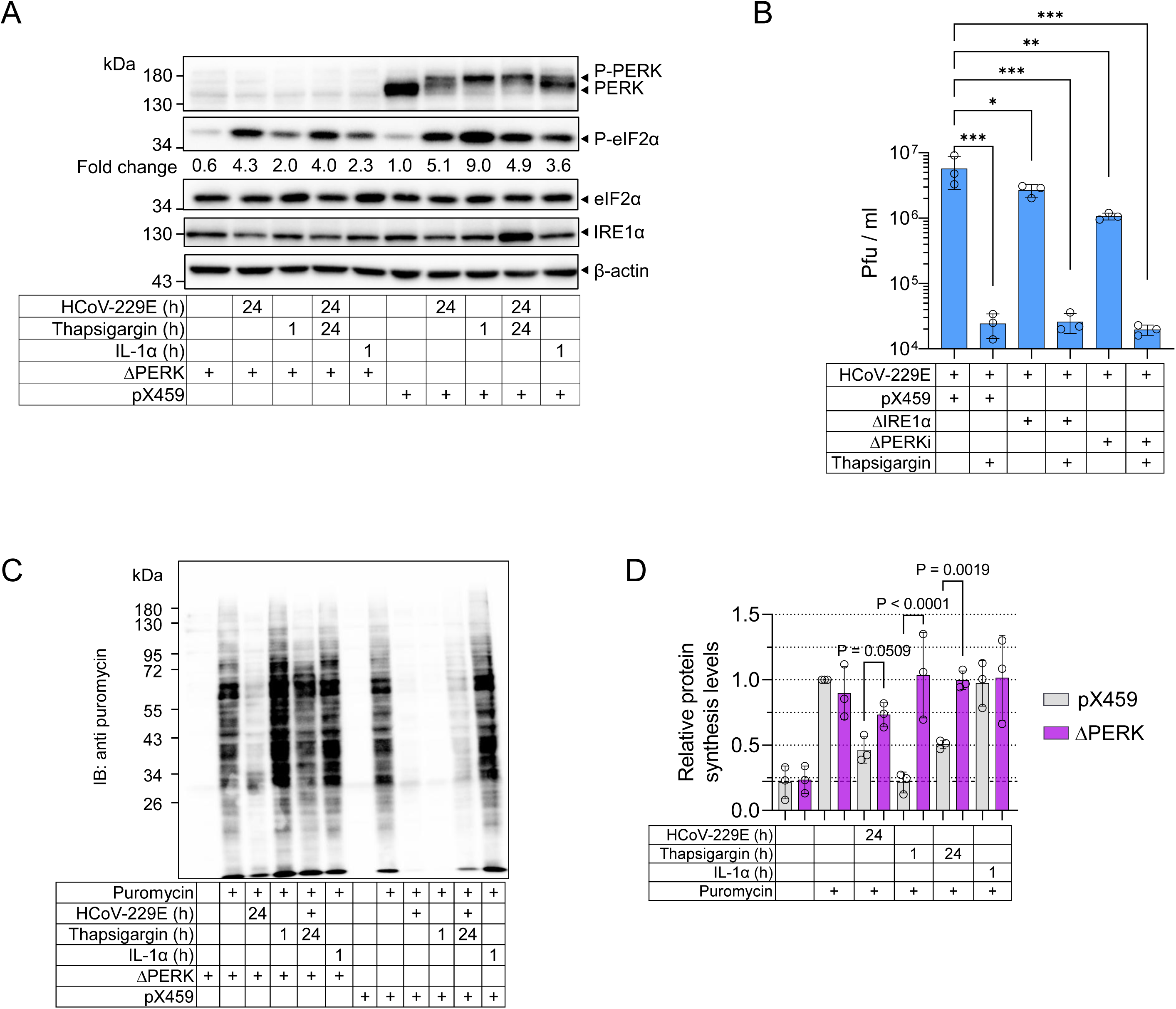
PERK deficiency suppresses viral replication and rescues CoV-mediated translational shutdown. (A) Huh7 cells stably transduced with Cas9 only (pX459) or with pX459 plus sgRNA against *EIF2AK3* (ΔPERK) were left untreated or were treated with 1 µM thapsigargin for 1 h or 24 h or with 10 ng/ml IL-1α for 1 h or were infected for 24 h with HCoV-229E at MOI of 1. Whole cell extracts were analyzed by immunoblotting for the phosphorylation or expression of the indicated proteins. (B) Huh7 cells stably transduced with Cas9 only (pX459) or with pX459 plus sgRNA against *EIF2AK3* (ΔPERK) or *ERN1* (ΔIRE1α) were left untreated or were treated for 24 h with thapsigargin (1 µM) or were infected for 24 h with HCoV-229E at MOI of 1. Viral titers were determined in supernatants by plaque assays. The graph shows the mean titers ± s.d. from three biologically independent experiments. Asterisks indicate p values (*p ≤ 0.05, **p ≤ 0.01, ***p ≤ 0.001) obtained by one-way ANOVA. (C-D) Huh7 cells were treated as in (A) and puromycin (3 µM) was added for the last 30 minutes. Total puromycinylation patterns were analyzed as described in Fig. 3. (C) Shown is one representative Western blot experiment out of three. (D) Graphs show mean levels of puromycinylation ± s.d. relative to the untreated control from three biologically independent experiments. P values obtained by one-way ANOVA are indicated.

**Supplementary Figure 12.**
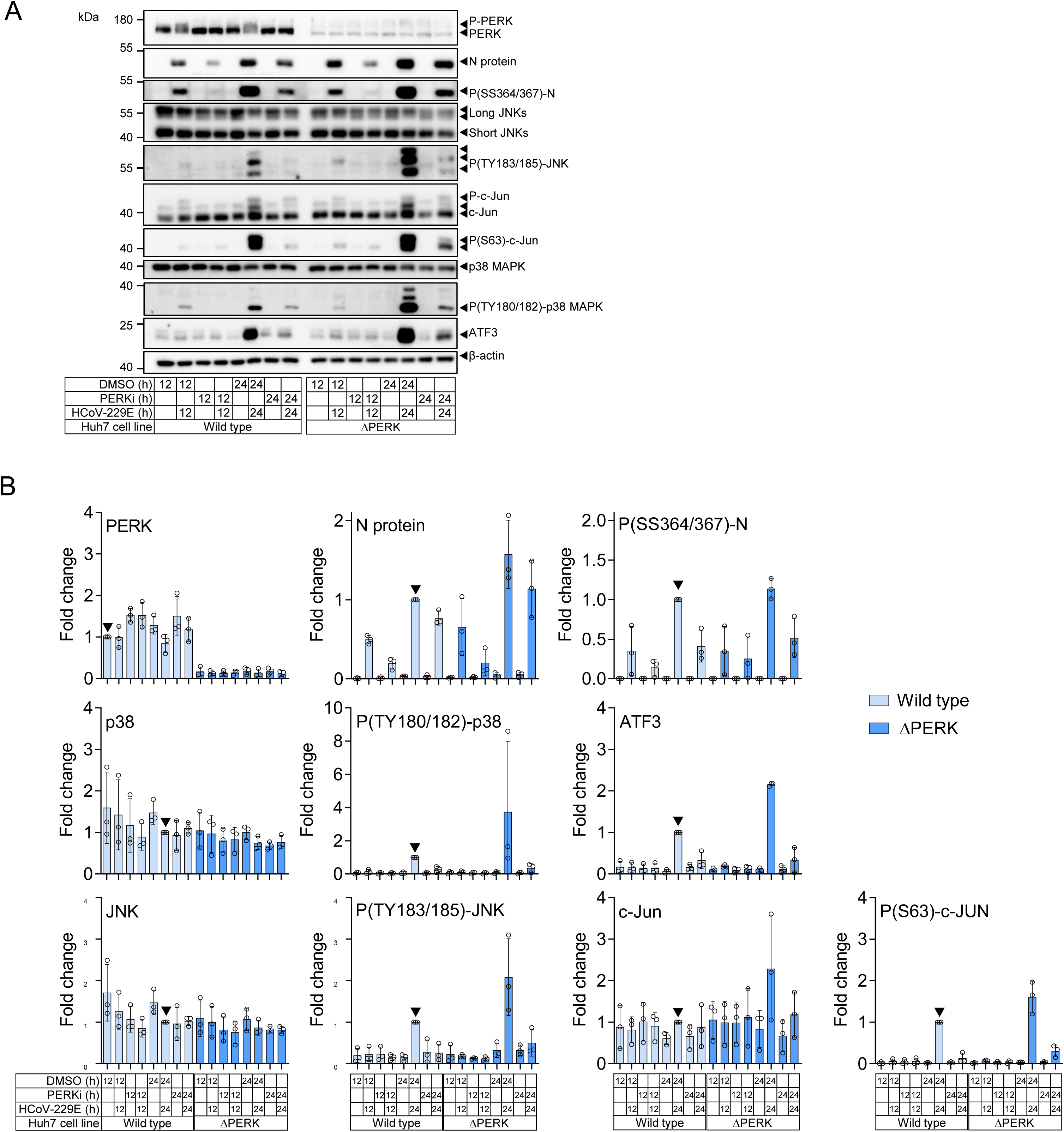
PERKi suppressive effects maintained in PERK-deficient cells define off-target effects. (A) Parental Huh7 cells (wild type) or stable cell lines lacking PERK (ΔPERK) were treated with solvent (DMSO) or were infected for 24 h with HCoV-229E at MOI of 1 in the presence or absence of 10 µM PERKi (GSK2656157). Whole cell extracts were analyzed by immunoblotting for the phosphorylation or expression of the indicated proteins. Shown is one representative Western blot experiment out of three. (B) Bar graphs showing mean changes of (P)-protein levels ± s.d. relative to the values from the 24 hpi time point (marked by black arrows) of the three experiments described in (A). PERK values were normalized relative to that of the DMSO control.

**Supplementary Figure 13.**
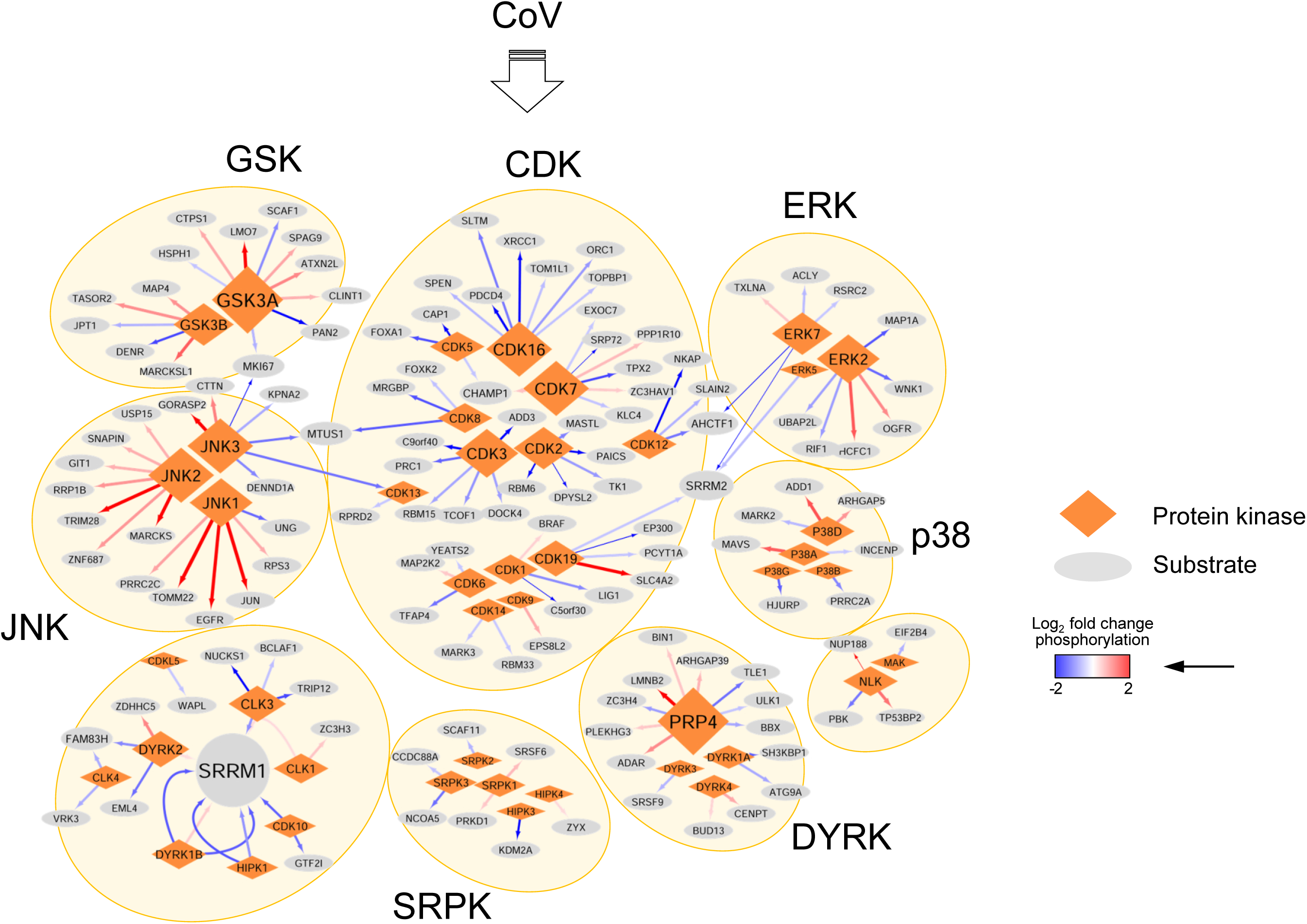
PERKi-sensitive CMGC kinase-substrate network in CoV-infected cells. Network reconstruction of kinase-substrate relations of the 44 protein kinases and 147 substrates shown in Fig. 7K-L. Protein kinases were arranged according to subgroups of the CMGC family shown in Fig. 7K and symbol size corresponds to number of substrates. Up- or downregulation of phosphorylated peptides is indicated by blue to red color and width of the arrows connecting a P-site to its rank1 kinase.

**Supplementary Figure 14.**
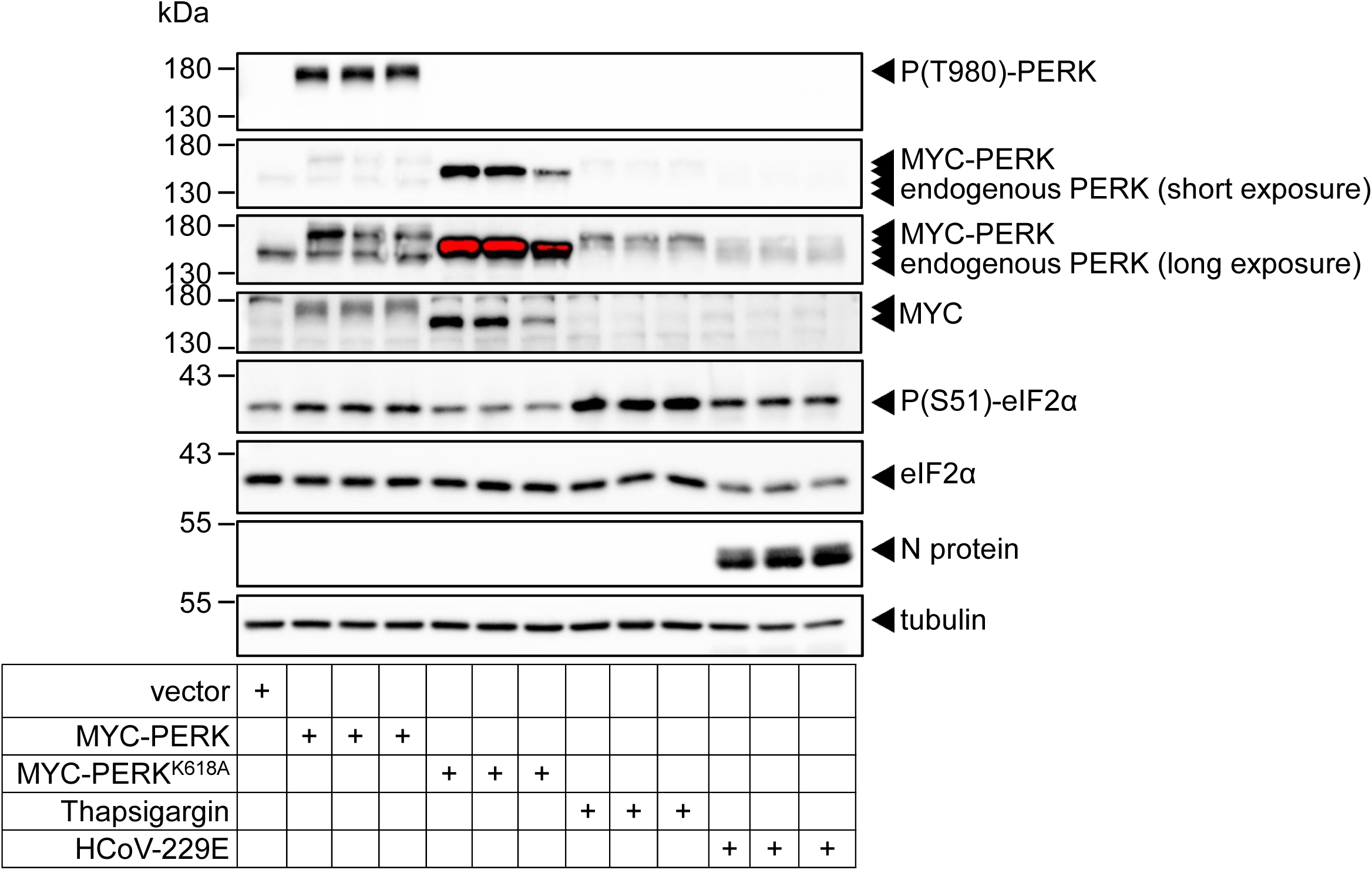
CoV infection does not activate Thr9S0 phosphorylation of PERK. Huh7 cells were transiently transfected with empty vector or expression vectors encoding Myc-tagged PERK or a catalytically inactive mutant (K618A) for 24 h. In parallel, cell cultures were treated for 1 h with thapsigargin (1 µM) or were infected with HCoV-229E for 24 h at MOI of 1. Cell extracts prepared at the end of the treatments were analyzed for the presence or regulation of the indicated (P)-proteins by Western blotting. Shown is one experiment performed with technical triplicates.

**Supplementary Fig. 15.**
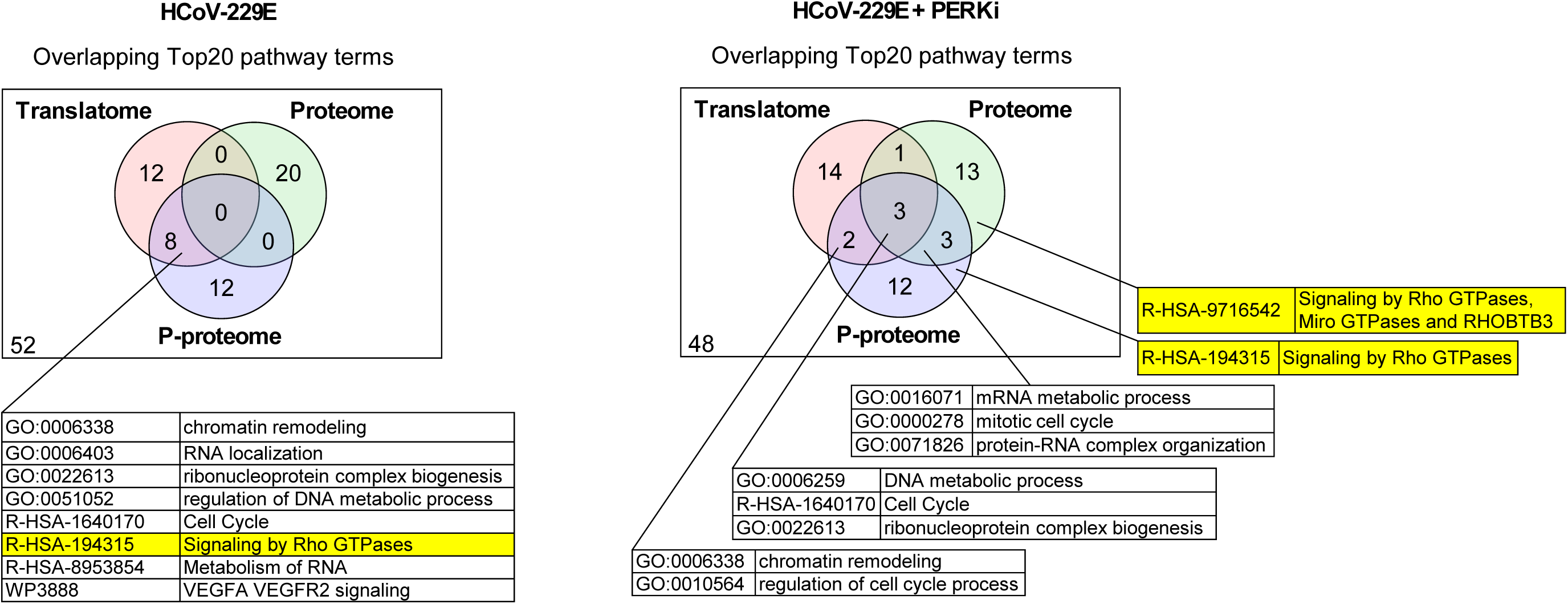
Pathway level overlap of translatomes, proteomes and P-proteomes. Overlap of the top20 pathway terms identified by Metascape summary analysis from all pooled differentially regulated nascent proteins (Fig. 3G), proteins (Fig. 4G and Supplementary Fig. 4) and P-proteins (Fig. 5E, Supplementary Fig.7A) in response to HCoV-229E at 12 hpi or 24 hpi (left Venn diagram) or in infected cells treated with PERKi (right Venn diagram). Yellow colors mark enrichment of small GTPase pathway components as highlighted in additional data sets of this study.

**Supplementary Figure 16.**
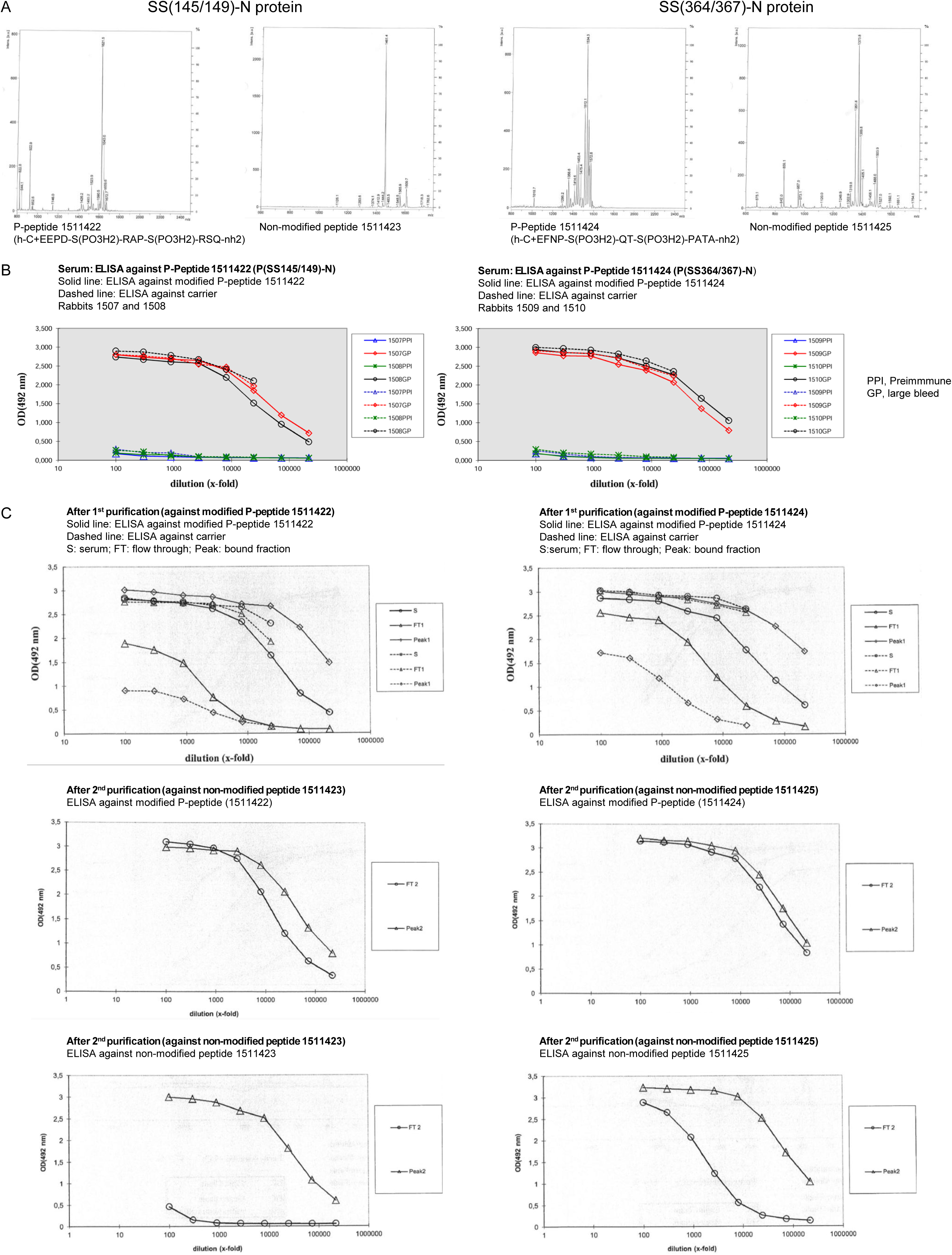
Generation and characterization of antibodies against phosphorylated HCoV-229E N protein. (A) Design and mass spectrometry-based quality control of bi-phosphorylated and non-phosphorylated immunogenic synthetic peptides corresponding to the HCoV-229E N protein. (B) Validation of antibody titers by ELISA in pre-immune sera and bleeds of two rabbits immunized against P-peptides shown in (A). (C) Validation of antibody titers in the serum of final bleeds (S) and after sequential affinity purification against columns coupled to P-peptide (1^st^ purification) and non-phosphorylated peptide (2^nd^ purification). Bound (peak) and unbound fractions (flow through, FT) were serially diluted and tested by ELISA against P-peptide or carrier. Peak2 fractions contain the purified P-specific polyclonal antibody preparations that were additionally depleted from antibodies binding to the non-phosphorylated N protein used in our study.

